# Decoding the Baltic Sea’s Past and Present: A simple Molecular Index for Ecosystem Assessment

**DOI:** 10.1101/2024.02.29.582456

**Authors:** Alexandra Schmidt, Juliane Romahn, Elinor Andrén, Anke Kremp, Jérôme Kaiser, Helge W. Arz, Olaf Dellwig, Miklós Bálint, Laura S. Epp

**Affiliations:** Department of Biology, University of Konstanz, Constance, Germany; International Max Planck Research School Quantitative Behaviour Ecology & Evolution, Constance, Germany; Senckenberg Biodiversity and Climate Research Centre, Frankfurt am Main, Germany; Loewe Center for Translational Biodiversity Genomics (LOEWE-TBG), Frankfurt am Main, Germany; Institute for Insect Biotechnology, Justus Liebig University, Giessen, Germany; Department of Natural Science, Technology and Environmental Studies, Södertörn University, Stockholm, Sweden; Leibniz Institute for Baltic Sea Research Warnemünde, Rostock, Germany

**Keywords:** ddPCR, biomonitoring phytoplankton, metabarcoding, dinoflagellate, diatom, sedimentary ancient DNA, sedaDNA, diatom/dinoflagellate index, ecological indicator, environmental impact

## Abstract

Marginal sea ecosystems, such as the Baltic Sea, are severely affected by anthropogenic pressures, such as climate warming, pollution, and eutrophication, which increased in the course of the past century. Biodiversity monitoring data and assessment of environmental status in such systems have typically been carried out only for the past few decades, if at all, and knowledge on pre-impact stability and good ecological status is limited. An extension of monitoring time series can potentially be achieved through analyses of paleoecological records, e.g. for phytoplankton, which form the base of the food web and are highly susceptible to environmental changes. Within the phytoplankton community, dinoflagellates and diatoms play a significant role as primary producers, and their relative dominance in the spring bloom, calculated as Dia/Dino index, is used as an indicator for the environmental status of the Baltic Sea. To extend time series on the dominance patterns and include non-fossilized dinoflagellates, we here establish a simple droplet digital PCR (ddPCR) reaction on ancient DNA from sediment cores that decodes phytoplankton dynamics. We focus on two common spring bloom species, the diatom *Skeletonema marinoi* and the dinoflagellate *Apocalathium malmogiense*, for which we evaluate a DNA based dominance index. It performs very well in comparison to DNA metabarcoding and modern monitoring and can elucidate past species dominance across the past century in three basins of the Baltic and across millennia in two of these basins. For the past century, we see a dominance shift already starting before the mid-20th century in two of the Baltic Sea basins, thus substantially predating current monitoring programs. Shifts are only partly coeval among the cores and the index shows different degrees of stability. This pattern is confirmed across millennia, where a long-term stable relationship between the diatom and the dinoflagellate is observed in the Eastern Gotland Basin, while data from the Gulf of Finland bear testimony to a much more unstable relationship. This confirms that good ecological status based on the dominance pattern of diatoms and dinoflagellates must be established locally and exemplifies how sediment core DNA can be employed to extend monitoring data.

## Introduction

Marine ecosystems, currently providing a main source of livelihood for three billion people worldwide (“Ocean economy and developing countries - OECD,” 2023), are under extreme pressure due to a variety of stressors (Pandion et al., 2022), such as current temperature rise (Cheng et al., 2022), biodiversity loss through overexploitation (Ward et al., 2022; Worm et al., 2006), pollution (Datta, 2023; Yu and Singh, 2023) and increased nutrient loads (Howarth et al., 2021; Malone and Newton, 2020). In particular, coastal marine systems and their inhabitants are increasingly affected by climate change and anthropogenic pressures, such as elevated nutrient loads (Breitburg et al., 2018). Over the past century, significant changes have been observed in these ecosystems, which are largely ongoing. However, systematic monitoring of biological communities typically only began a few decades ago. This has resulted in a lack of data on communities prior to the onset of these alteration, i.e. on pre-impact reference states, on the exact timing and potential historic causes of the anthropogenic community changes (Costello et al., 2017; Obst et al., 2020; Van De Putte et al., 2021) and on the long-term stability of the ecosystem prior to the current rise in anthropogenic pressure.

The few existing biodiversity records that extend beyond the monitoring time series, such as historical plankton counts (Wasmund, 2017), cannot fully address these issues. They do not extend beyond the realm of historical memory, are not continuous in time and exist only for a few locations. The latter aspect limits the possibility to investigate dynamics and establish pre-impact reference states and good ecological status (GES) on local scales, i.e. for separate basins in marine systems. This shortcoming can potentially be addressed through the use of paleoecological archives, from which the history of ecosystems and their constituent species can be studied through the fossil record (Nguyen et al., 2023; Wingard et al., 2017).

Paleoecological archives, such as sediment cores, can deliver data on past aquatic assemblages from a multitude of sites and thus provide relevant local time series. They can elucidate the range of natural variation and the speed and trajectory of change, and thus provide a long-term perspective to evaluate present day observations (Saunders and Taffs, 2009). However, the direct incorporation of paleoecological datasets and the subsequent extension of monitoring data into the past is complicated by several factors.

In particular, records display differences in temporal resolution of sampling and differences in taxonomic resolution and taxonomic identification. Current monitoring datasets are typically based on taking discrete samples or making discrete observations at specific timepoints across the year and can take seasonal patterns into account. In contrast, samples from paleoecological archives, such as slices of sediment cores, integrate information from longer periods, such as a complete year or several years. Furthermore, living organisms from current ecosystems can generally be identified to species level visually, while information from paleoecological archives is based on remains of organisms, which are often not identifiable to species level. These can be pollen or microfossils of organisms that contain hard parts or shells (McQUOID et al., 2002), or also cysts and spores of otherwise soft-bodied plankton (Rengefors et al., 1998; Sundqvist et al., 2018). Many of these remains lack species-specific features, allowing identification only to a higher taxonomic level such as genus or family. In addition, they can become damaged or degraded in the course of sediment formation and over time. Finally, a large part of aquatic organisms are soft-bodied in all life stages and are thus absent from the visual fossil record.

These problems can be diminished or overcome by using not visible, but molecular remains, in particular ancient DNA preserved in sediments (sedimentary ancient DNA or sedaDNA). This delivers a record of aquatic communities with a high degree of spatial fidelity (Wang et al., 2023), making it suitable for investigations of local aquatic ecosystem history. Focusing on highly variable genomic markers, sedaDNA offers the potential for high taxonomic resolution, discriminating between sister species and reaching population level (Epp et al., 2018; Lammers et al., 2021). It also includes taxa that are not well-preserved in the fossil record (Nguyen et al., 2023), such as a number of phytoplankton taxa lacking rigid structures.

Among these are dinoflagellates, which, together with diatoms, represent a dominant group of phytoplankton world-wide (Bi et al., 2021; Kang et al., 2021; Zhang et al., 2019). In temperate waters, these groups are often typical components of the spring bloom. Due to differences in nutritional value, biochemical composition, and phenology of diatoms and dinoflagellates, their relative dominance and the timing of their temporal succession has consequences for the complete ecosystem (Wasmund et al., 2017). Current global changes have induced shifts in their relative dominance and successions (Bi et al., 2021; Zhang et al., 2019).

For example in the Baltic Sea, where the two groups occur simultaneously in the spring bloom, a shift in dominance from diatoms to dinoflagellates has lately been observed, with far reaching ecological consequences (Spilling et al., 2018). In particular, it influences the function of the food web: the increased deposition of pelagic diatoms to the sea floor provides an abundant food supply for zoobenthos, while the pelagic dinoflagellates provide a food source for zooplankton or encyst. A shift from diatoms to dinoflagellates thus signifies a shift from a nutrient pathway favoring benthic organisms to more nutrients for the pelagic components of the food web (Wasmund et al., 2017). Furthermore, a shift in dominance from diatoms to dinoflagellates is also linked to the trophic status of the Baltic Sea. It may indicate silicate limitation caused by eutrophication (Conley et al., 2008), given that diatoms rely on silicate. Thus, a regime shift towards dinoflagellate dominance is considered an indicator for eutrophication (Wasmund et al., 2017).

The proportions of diatoms and dinoflagellates in the spring bloom have shifted during the past decades towards higher dinoflagellate abundance. This development is attributed to both eutrophication, mainly caused by anthropogenic discharge of nitrogen and phosphorus, and to climate warming (Klais et al., 2011). The degree of this change in proportions of the two taxa has been utilized to develop an index for the environmental status of the Baltic Sea, used in the assessments of the Marine Strategy Framework Directive (MSFD) (Wasmund et al., 2017). It is currently calculated based on microscopically produced phytoplankton biomass data collected in regular biomonitoring surveys of the HELCOM monitoring and assessment program (Wasmund et al., 2017), using seasonal averages for diatom and dinoflagellate biomass.

These regular biomonitoring surveys in the Baltic Sea were initiated in the 1980s due to the tremendous anthropogenic pressures on its unique marine and coastal environment (Storie et al., 2021). Within this timeframe, the Dia/Dino-index identified a regime shift at the end of the 1980s in the Baltic Proper, which has been attributed to climate warming (Wasmund et al., 2017). However, at the start of the biomonitoring, changes in trophic state had already exerted stress on the complete Baltic ecosystem and its communities (Mack et al., 2020; Storie et al., 2021). Eutrophication predates the 1980s, based on available nutrient data extending back to 1900 (HELCOM, 2021; Wasmund, 2017). Thus, the ecosystem’s responses were putatively already underway, rendering it impossible to accurately determine the onset of shifts in response to multiple impacts of the past century, or to define values for the index that would characterize a potential pre-impact reference state, or GES. The initial shift, and the conditions preceding it, can only be investigated by turning to historical data or paleoenvironmental archives.

Historical records of plankton counts, although few and temporally patchy, confirm a clear dominance of diatoms over dinoflagellates at the beginning of the 20th century, with higher index values than in later years (Wasmund, 2017). Across this timescale, there seems to be a clear trend related to human impact, and no fluctuation of the index. However, records are not continuous, missing large parts of the past century. In addition, the existing data - historic and recent - also show a clear heterogeneity in the index values, prompting the development of different values for the GES for different basins (Wasmund, 2017). As historic data are not available for all basins, the question of pre-impact stability cannot be conclusively answered for the complete Baltic Sea, also considering the relatively short period covered in relation to the history of the ecosystem. Analyzing the diatom/dinoflagellate ratio on longer, millennial time scales, where they were governed by natural environmental changes, can elucidate the relevance of recent, anthropogenic impact on the index.

In this study, we develop and test a simple sedaDNA approach relying on species-specific PCR reactions to investigate this stability across the 20th century and across the millennial history of the Baltic Sea. The approach exclusively targets key members of the spring-bloom in the diatoms and the dinoflagellates, i.e., the diatom *Skeletonema marinoi* (Sarno et al., 2005) and the dinoflagellate *Apocalathium malmogiense* (Craveiro et al., 2017). Both provide stable records through the Baltic history and can therefore be used as models, representing the two major taxonomic groups at large. We evaluate quantitative results obtained by digital droplet PCR (ddPCR). This method provides precise, sensitive, and robust quantification of nucleic acids without the need for external standards, while being able to perform massive sample partitioning, leading to reliable and sensitive measurements (NRCM and Bio-Rad, 2020). We derive an index from these results, which is equivalent to the existing Dia/Dino index, and compare resulting values to data obtained from metabarcoding of sedaDNA and to recent Baltic Sea biomonitoring data. We thus introduce a cost-effective and efficient molecular tool, designed to reconstruct the historical dynamics of these key taxa with the potential to be used also in current monitoring applications on modern environmental DNA (eDNA). We investigate 1) how this approach compares to other employed methods, 2) the initial timing of the recent, anthropogenically induced dominance shift from diatoms to dinoflagellates and 3) the degree of stability of the pre-impact reference state across millennia in relation to natural environmental changes in two open Baltic Sea locations.

## 1. Material & methods

### 1.1. Study area

The Baltic Sea is a brackish, shallow, and semi-enclosed sea with limited water exchange to the North Sea and lower biodiversity than fully marine systems (Snoeijs-Leijonmalm et al., 2017) and is especially at risk due to a multitude of reasons. It is particularly vulnerable due to its low biodiversity and its unique position as a sea enclosed by land, densely populated human settlements, and high industrial activity. It suffers from decreasing oxygen levels caused by a number of intertwined processes. Higher oxygen levels are, for one, induced by the higher water temperatures, which diminish the amount of dissolved oxygen and lead to an increase in oxygen-deficient areas (Ito et al., 2017). In addition, eutrophication, caused among other factors by the input of agricultural waste, such as fertilizers, exacerbates the formation of anoxic areas (Carstensen et al., 2014). Discharge of nutrients, such as nitrogen and phosphorus, furthermore promotes cyanobacteria and algal blooms. These primary producers consume nutrients and fix carbon, but they also lead to an increase of decomposing biomatter in the benthic zone. Bacterial decomposition again consumes oxygen (Conley et al., 2009). Taken together, increasing hypoxia is a major challenge facing the Baltic Sea. The Baltic Marine Environment Protection Commission (HELCOM), the governing body for the Baltic Sea, is actively working to reverse the impacts of anthropogenic pressures, including efforts to reduce nutrient levels (HELCOM, 2021).

These recent pressures are not the first challenges the Baltic Sea has encountered (Borzenkova et al., 2015; Emeis et al., 2002). Since the latest glaciation, the Baltic Sea has undergone profound changes (Snoeijs-Leijonmalm and Andrén, 2017). Following the retreat of the Scandinavian ice sheet, the Baltic Sea underwent a number of distinct phases. The initial glacial retreat left a giant meltwater lake, the Baltic Ice Lake (12050-9750 yr BCE, years Before Present) in the Baltic basin, which finally drained as a connection to the ocean opened over middle Sweden and became the Yoldia Sea (9750-8750 yr BCE). Further meltwater and isostatic rebound of the Scandinavian land masses led to the closure of the oceanic connection, and the formation of the freshwater Ancylus Lake (8750-7850 yr BCE). After a transitional, almost freshwater phase, the Initial Littorina Sea (7850-5550 yr BCE), the salinity increased, related to a rising sea level, and the brackish Littorina Sea phase started around 5550 BCE (Andrén et al., 2011). Between 5550-2050 yr BCE, during the Holocene Thermal Maximum, the temperature in the Baltic Sea region was 1-2 °C above modern values (Seppä et al., 2009) and the Baltic Sea salinity reached a maximum between 4050-2050 yr BCE (Gustafsson and Westman, 2002). Both temperature and salinity decreased between 2050 and 50 yr BCE (Seppä et al., 2009; Snoeijs-Leijonmalm and Andrén, 2017). The Baltic Sea temperature increased again during the Medieval Climate Anomaly (950-1250 yr CE; years Common Era; Mann et al., 2009) and the Modern Warm Period (since 1850 yr CE) (Kabel et al., 2012).

### 1.2. Study organisms

Diatoms and dinoflagellates are two major taxonomic groups of the Baltic Sea phytoplankton (Andersson et al., 2017). These unicellular microalgae are important primary producers and thus play a crucial role in carbon fixation. While diatoms generally predominate during the cold season, dinoflagellates often thrive in warm water. However, in the Baltic Sea, cold adapted dinoflagellates grow to high biomasses in spring, co-occurring and even dominating over diatoms under certain conditions (Spilling et al., 2018). Within these groups, the cold-water diatom *Skeletonema marinoi* and the cold-adapted dinoflagellate *Apocalathium malmogiense* are particularly abundant species of the spring bloom (Hällfors, 2004) and characterize the phytoplankton communities. Both form resting stages, spores and cysts, respectively, to overcome unfavorable conditions such as during the warm-season (Hinners et al., 2017; Stenow et al., 2020). For both species, continuous decadal and centennial records of living resting stages have been established from Baltic coastal sediments as well as deposits of the deep basins (Härnström et al., 2011; Kremp et al., 2018).

Here we investigated how DNA based time series of the two groups (Diatoms and Dinoflagellates), and records of the two representative species of the spring bloom, *A. malmogiense* and *S. marinoi*, specifically, compare to existing monitoring data. We evaluate information on millennial scale changes of phytoplankton in relation to environmental status of the Baltic Sea and long-term fluctuations.

### 1.3. Sampling & Data acquisition

Our study is based on two main sources of data: biomonitoring data and data extracted from sediment archives. By combining and comparing these two datasets, we aim to establish a comprehensive time series of key phytoplankton species and extend existing monitoring approaches. This approach allows us to understand not only the current state of these communities but also their historical changes and potential future trends.

Five sediment cores were retrieved (Fig. 1) in April 2021, during expedition EMB262 in April 2021 onboard the research vessel Elisabeth Mann Borgese from three different locations (Fig. 1): 1) Eastern Gotland Basin (EGB, 57°17.004’N, 020°07.244’E, Water Depth: 241 m water depth, cores EMB262/6-28MUC and EMB262/6-30GC); 2) Gulf of Finland (GOF, 59°34.443’N, 023°36.461’E, Water Depth: 81 m bsl, cores EMB262/12-2MUC and EMB262/12-3GC). 3) Landsort Deep (LD, 58°38.391’N, 018°15.997’E, Water Depth: 436m bsl, core EMB262/13-8MUC) (see supplementary Table S1). Short cores (ca. 50 cm) were retrieved at all locations using a multicorer (MUC) preserving the water-sediment interface undisturbed. Long cores (ca. 500 cm) were retrieved from the EGB and GOF using a gravity corer (GC). The EGB long core sampling began at a depth of 32 cm, corresponding to the 1980s. Thus, it misses the final decades, which are only captured in the short core. The cores were sampled onboard using sterile syringes according to Epp et al. (2019) and immediately frozen for storage.

**Fig. 1.**
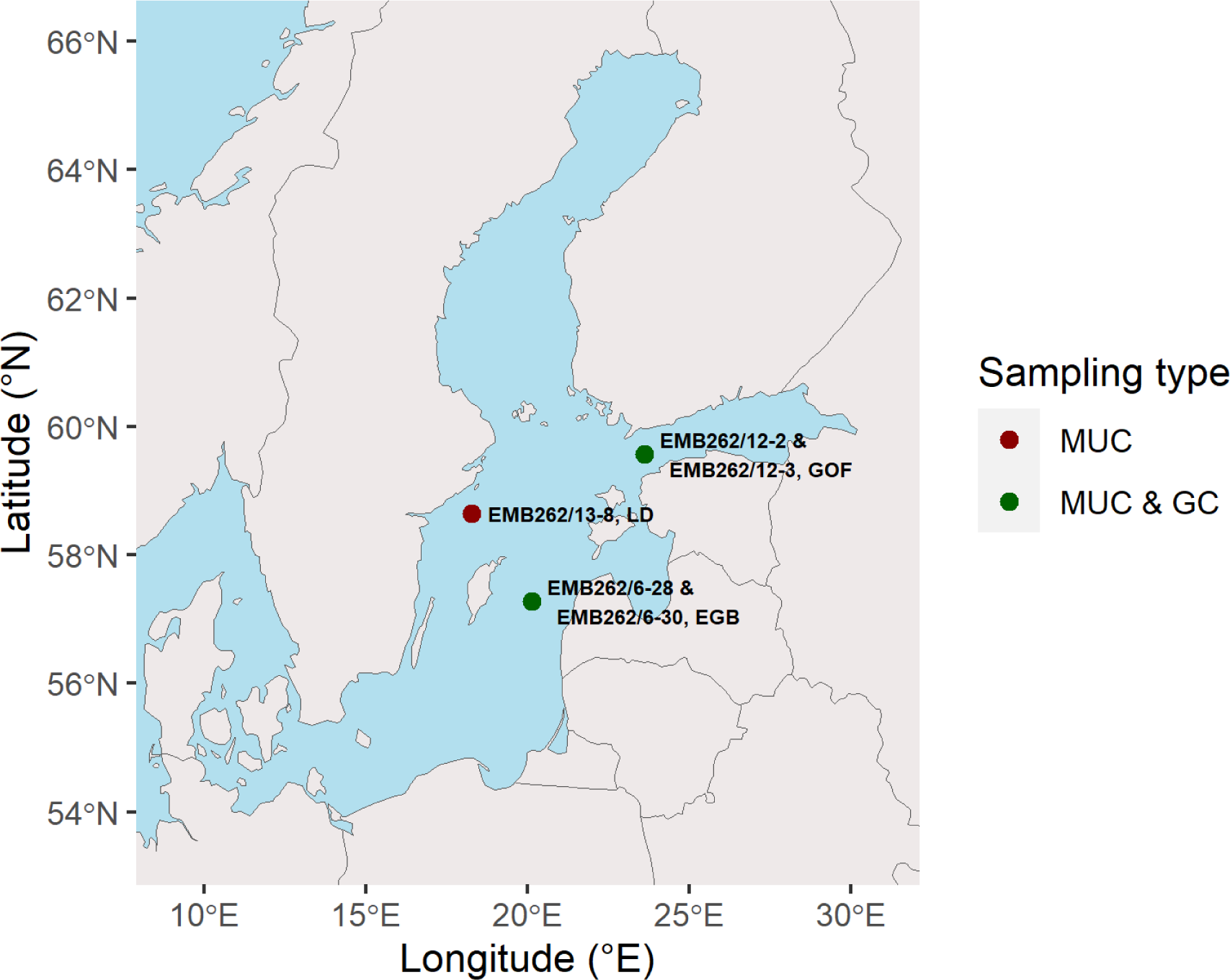
Location of the coring sites in the central Baltic Sea. Gulf of Finland (GOF; cores EMB262/12-2MUC and EMB262/12-3GC), Landsort Deep (LD; core EMB262/13-8MUC) and Eastern Gotland Basin (EGB; cores EMB262/6-28MUC and EMB262/6-30GC)

In addition to the sediment cores, biomonitoring data, specifically phytoplankton monitoring data, were obtained from the ICES database and filtered based on proximity to the sediment core locations. We focused on data from the spring bloom period (March to May) and excluded years with incomplete monitoring data. The data were processed to join size classes and averaged between stations per month. Yearly biomass proxies were calculated for general diatom and dinoflagellate data as well as for specific genera.

#### Core Dating

##### 1.3.1. Short cores (MUCs)

The short sediment cores from the EGB (EMB262/6-28MUC) and LD (EMB262/13-8MUC) were dated by transferring published age models of short sediment cores from the same or nearby locations (Fig. S1; Häusler et al., 2018; Kaiser et al., 2023). For the GOF short core (EMB262/12-2MUC), an event stratigraphy approach has been applied, which is a current method used to date recent sediments of the Baltic Sea (Dellwig et al., 2018; Häusler et al., 2018; Kaiser et al., 2023; Moros et al., 2017). Three parameters were considered to build the short core stratigraphy: the activity of ^241^Am and ^137^Cs radionuclides, and the Hg content. Altogether, five stratigraphical events, or time markers, were used to date the sediment (Figure S1). The uncertainties of the time markers were estimated as in Kaiser et al. (2023). The early increase in Pb content reflects the early increase in coal combustion around 1870 CE related to the beginning of the Second Industrial Revolution. A sharpening in Pb increase starting around 1950 CE is related to a further increase in coal combustion after the Second World War. The early increase and peak in ^241^Am activity reflect the beginning and maximum in the global fallout related to the atmospheric nuclear bomb tests in 1953 and 1963 CE, respectively. The synchronous increase in ^241^Am and ^137^Cs has been attributed to the Chernobyl accident in 1986 CE. The year of the core recovery (2021 CE) was attributed to the core top layer. Linear sedimentation rates were assumed between the time markers.

##### 1.3.2. Long cores (GCs)

Organic carbon records from post-Littorina-transgression sediments in the central Baltic Sea exhibit patterns ideal for detailed inter-core correlations (Moros et al., 2020; Warden et al., 2017). The chronology of core EMB262/6-30GC from the Eastern Gotland basin is based on such a correlation using the relative content of S and the Br/K ratio (XRF scanner elemental data), which reflect changes in the bulk organic carbon content of the sediments (REFS). The data were visually tuned with XRF scanner and organic carbon data of dated sediment cores from the central Baltic Sea (Figure S2). For the mid-Holocene part (∼5000-3000 BCE), the chronology of sediment core P435/2-1 (Warden et al., 2017), retrieved from the same location as EMB262/6-30 in the EGB, was used as a reference. For the late Holocene part (the last ∼3500 years), sediment cores M86-1a/36-4GC (Häusler et al., 2018) and MSM62-60GC (Moros et al., 2020) from the western Gotland Basin were used as reference chronologies. Precise age assignment of pre-Littorina-Stage sediments was not possible, but the oldest sediments of the core likely belong to the late Ancylus Lake phase (< 6500 BCE) of the Baltic Sea.

For sediment core EMB262/12-3GC from the Gulf of Finland, an independent Bayesian age model using BACON 2.5.5 (Blaauw and Christen, 2011) was established based on seven radiocarbon dates of the bulk organic matter in the sediment (Figure S3, Table S4). With average sedimentation rates of ∼100 cm/1000 years, the core dates back to ∼2300 BCE.

### 1.4. Laboratory work

#### 1.4.1. DNA Extraction

DNA was extracted from all sediment samples (n = 226, see supplementary Table S2 & S3) using the PowerSoil Pro Kit from Qiagen (Hilden Germany) and a starting amount of ∼0.5 g, resulting in 226 DNA extracts and 22 extraction blanks. From 15 horizons more than one extract was generated due to additional pre-tests. The extraction followed the manufacturer’s protocol with an additional overnight incubation at 56°C and 20µL of proteinase K (20mg/ml) to optimize lysis. The washing steps were performed using a Qiagen Vacuum Pump (Hilden, Germany) following the manufacturer’s instructions. Afterwards, we centrifuged for 3 minutes instead of the recommended 2 minutes before conducting the elution. The spin columns were incubated twice with 75 μl of elution buffer for 5 minutes and eluted into the same tube.

#### 1.4.2. Metabarcoding

Comprehensive changes in phytoplankton communities were investigated across all sediment cores using DNA metabarcoding, targeting all eukaryotic organisms with the Euka02, or Allshorts, primer pair (forward: 5’-TTTGTCTGSTTAATTSCG-3’, reverse: 5’-CACAGACCTGTTATTGC-3’) (Guardiola et al., 2015). The PCR setup was automated using the workstation Biomek i7 (Beckman Coulter GmbH) and included 2 PCR positive controls, 4 PCR negative controls, and 8 multiplexing controls, including the extraction controls for 4 PCR replicates. Extracts were 1:2 diluted and amplified using the AmpliTaq Gold ™ Mastermix (Thermo Fisher). The PCR amplification started with an initial denaturation step at 94°C for 5 min, followed by 40 cycles of 94°C for 30s, 45°C for 30s, 68°C for 45s and a final elongation step at 72 °C for 10 min. Unique tag combinations were used to separate replicates and samples (Taberlet et al., 2018). All PCR products were equally pooled and 1200 µL of each pool was purified using the MinElute PCR Purification Kit (Qiagen) according to manufacturer’s instructions. PCR-free library preparation and paired-end sequencing (2 × 150 bp NovaSeq 525M) were performed at Fasteris SA (Geneva, Switzerland).

#### 1.4.3. Droplet digital PCR (ddPCR) on the target species

A duplex droplet digital PCR was performed to quantitatively investigate the DNA concentration of the two key target species. With this duplex reaction the concentrations of both species could be simultaneously determined and thus directly compared. To this end, two species-specific primers were designed to target the study organisms *S. marinoi* and *A. malmogiense*. ITS1, 5.8S rRNA and ITS2 sequences from GenBank (Benson et al., 2015) were downloaded and aligned separately for both species using the MUSCLE alignment function in Geneious Prime (v. 21.2.2). Primer pairs were manually selected and compared by eye to the respective species alignment and outgroups (*Homo sapiens*, *Nannochloropsis limnetica*) based on criteria such as short fragment length and specificity to the target species. The primers were then tested and optimized in four steps: 1) primer specificity and taxonomic resolution was determined by running *in silico* PCR with the program ecoPCR (OBITools ecoPCR, version 1.0.0, Ficetola et al., 2010); 2) the optimal annealing temperature *in vitro* was determined by gradient PCR on DNA extracts of tissue and upper sediment layers; 3) specificity of the reactions was tested *in vitro* by running PCRs on DNA of both target species (tissue from stock cell cultures and resurrected cultures) and non-target environmental samples from Lake Constance Site S23 (Wang et al., 2023); and 4) PCR with sedaDNA. From these tests, an optimal primer pair for *A. malmogiense*(forward: 5’-GATACCCTTGTGCAGAAACTC-3’, reverse: 5’-ATTAGACAAGAAGCAAGAAGTAG-3’) and *S. marinoi* was selected (forward: 5’-CTTGTGAGTTGCCGAGGC-3’, reverse: 5’-TCCATAGATGAGGTACATTCAT-3’).

The ddPCRs were run in a probe-based assay, for which ddPCR probes specific to the amplicon were designed using Geneious Prime (v. 22.1.1) according to the Bio-Rad Laboratories (Hercules CA, USA) application guide. The *A. malmogiense* probe was 19 base pairs in length (GCAGGATGGGTGCTTGTCA), while the *S. marinoi* probe was 20 base pairs long (TCTCCAGCGAATTGGGCTAC). Both probes were double quenched in a duplex ddPCR reaction, with *A. malmogiense* marked by FAM (blue) and *S. marinoi* by HEX (green) fluorescent dyes.

Each 22 µL ddPCR reaction contained the following five components: 1) 11 µL ddPCR supermix for probes (Bio-Rad); 2) 6.8 µL DEPC treated H_2_O; 3) 1.1 µL 20x Target-Primers/Probes FAM; 4) 1.1 v 20x Target-Primers/Probes HEX; 5) 2 µL DNA sample. The ddPCR was performed according to the Bio-Rad manufacturer’s protocol. 21 µL of reaction mix was transferred to the Bio-Rad QX200 droplet generator, avoiding bubble formation by only pushing the pipette to the first pressure point. The resulting droplets were transferred to a 96 well PCR plate for amplification on a Bio-Rad C1000 Touch^TM^ thermal cycler using the following program: 10 min at 95°C, 40 cycles of 30s at 94°C, 30s at 50°C and 30s at 60°C, followed by a final enzyme deactivation at 98°C for 10 min. The Bio-Rad QX200 droplet reader and the corresponding QuantaSoft^TM^ Analysis Pro software (v. 1.0.596) were used to analyze the plate and determine the number of positive droplets, measured in copies per µL. Three replicates of the 226 DNA extracts and 22 extraction blanks were run, with each run consisting of 56 PCR reactions, two master mixes, and two NTCs.

### 1.5. Preparation of datasets

Four different datasets were prepared for use in common analyses: 1) ddPCR concentrations from sediment core DNA specific to the two species abundant in the spring bloom, 2) read numbers of the two phyla from DNA metabarcoding on sediment core DNA, 3) relative abundance of the two phyla from biomonitoring counts and 4) relative abundance of the two species from biomonitoring counts. To compare the four datasets, both the overall relative abundance as well as 5-year intervals (sum of 5 years) were used. The latter can account for the accumulation of molecular compounds in the sediment samples, which likely result from several years.

To ensure that it is valid to use the four datasets quantitatively, we implemented a consistency check. This check is based on the five-year intervals of all four datasets described above. We restricted our analyses of correlation between datasets to data derived from the short cores due to their temporal resolution, particularly in the years when biomonitoring was initiated (∼1980s). Importantly, only the years where data was available across all four datasets were included in the comparison. This approach ensured a comprehensive comparison.

#### 1.5.1 ddPCR

The concentrations of the three replicates per sample were correlated for consistency, with each dye concentration (FAM and HEX) representing a separate reaction (supplementary Fig S4). Afterwards, the average DNA concentration of *S. marinoi* and *A. malmogiense* per sample was determined by ddPCR. Afterwards, the results were normalized based on the weight of the sediment used in the DNA extraction process. This allowed for a more accurate comparison of the data across samples.

#### 1.5.2 DNA metabarcoding

Raw metabarcoding paired-end reads were processed using ObiTools3 (v.3.01b13). Read pairs were merged, and sequences with an average alignment score less than 0.8 were discarded. Afterwards reads were demultiplexed, dereplicated and pre-filtered based on count and length. Reads with low counts (<10 counts) were removed and only those within a specified length range (80-250 bp) were retained. Reads were assigned using mothur (v.1.40.4) (Schloss et al., 2009) against the pr2 database (v. 4.14.0) (Vaulot et al., 2022) with a consensus confidence threshold of 80%.

The data were further filtered in R (v 4.1.3). ASVs were defined as rare based on frequency plots, and ASVs with low read counts (<90 reads) or present in few replicates (<10 replicates) were discarded. Additionally, filtering according to ObiTools3 (Flück et al., 2022) was performed to remove ASVs with higher “internal” counts than head or singleton counts. The maximum read count of each ASV in the negative controls was subtracted from the dataset. Read counts were normalized based on wet sample weight. ASVs belonging to dinoflagellates or diatoms were summed for each replicate (Table S5). Dinoflagellate and diatom community dissimilarities were assessed across all PCR replicates using nonmetric multidimensional scaling (NMDS) via the metaDMS function (seed number 12345) from the R-package vegan (Dixon, 2003). Ordination plots were generated for each short core location.

#### 1.5.3 Phytoplankton monitoring data

Phytoplankton monitoring data were obtained from the ICES database (ICES Data Portal, Dome Phytoplankton. (2022), October 18^th^) and filtered based on proximity to the sediment core locations (in a circle 5% around the latitudes and longitudes, if considered as decimal numeric value). Species names were verified and taxonomy obtained from WoRMS (https://www.marinespecies.org/) using the wm_record function of the “worrms” package v0.4.3 (Chamberlain and Vanhoorne, 2023). The data were filtered for dinoflagellates and diatoms. Only the spring bloom monitoring data was investigated and thus three months from March to May were used. Years with incomplete monitoring data were excluded. Data count was processed to join size classes and averaged between stations per month. Yearly biomass proxies were calculated for general diatom and dinoflagellate data as well as for specific genera. For the correlation with the molecular data, biomonitoring count data were retained only for the genera *Skeletonema*, *Apocalathium*, *Scrippsiella* and *Peridinium*. Due to changes in systematics and species concepts, *Scrippsiella* and *Peridinium* were treated as *Apocalathium*. The average count data per month and species were calculated and summed over the three months as yearly biomass proxy.

#### 1.5.4 Paleoenvironmental conditions

Data on the history of the Baltic Sea and proxies for paleoenvironmental conditions was visualized using results from the core and published records.The annual mean air temperature was based on a core from Lake Flarken in southern Sweden (Seppä et al., 2005), providing a representative depiction of the climate in the Baltic Sea area. The Mn/Ti value from our EGB core was utilized as a proxy for oxygen levels (Supplementary Data, Core dating).

### 1.6 Analyses

#### 1.6.1 Correlations among datasets

All four datasets were pairwise correlated against each other, calculating Pearson’s correlations. Rows that did not show measurements in all datasets (presence of NAs) and outliers were removed according to the interquartile range method, calculating the first (0.25) and third (0.75) quartile of the input vector and the interquartile range (IQR) of the input vector. Then the IQR was multiplied by 1.5 to get H, which determines the threshold for which data points are considered outliers. Any variable that is less than the first quartile minus H, or greater than the third quartile plus H, was treated as outlier and was removed. No outliers were found using the interquartile range method, so no data points were removed.

For the analyses and visualizations, the following R packages were utilized: corrplot v0.92 (Taiyun, 2023), scales v1.2.1 (Wickham et al., 2022), ggpubr v0.6.0 (Kassambara, 2023), ggimage v0.3.3 (Yu, 2023), gridExtra v2.3 (Auguie and Antonov, 2017) and cowplot v1.1.1 (Wilke, 2020). Additionally, the biomonitoring Dia/Dino-index was correlated against the biomonitoring ske/apo-index (extracted from the biomonitoring data) to evaluate if the ske/apo-index represents the Dia/Dino-index. Prior to performing further analysis with the molecular ddPCR ske/apo-index, we controlled for any correlation between ddPCR concentrations of *A. malmogiense* and *S. marinoi*. This was done to ensure that any observed trends in the concentration of each species were independent of the overall DNA concentration fluctuations in the sediment cores.

#### 1.6.2 Ecological indices derived from the data

The ratio between ddPCR concentrations of *Skeletonema marinoi* and *Apocalathium malmogiense*, defined as the ske/apo-index, for each horizon, 5-year interval and the biomonitoring data was calculated using the following formula:

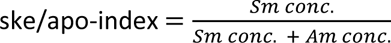

The Dia/Dino-index of the biomonitoring data is based on Wasmund 2017. It was calculated with the following formula (Wasmund et al., 2017):

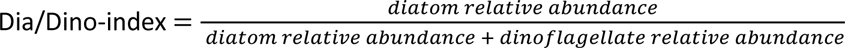

The relative abundance of a group or species at a sampling site was determined by calculating its proportion of the overall biomass at the site. The ddPCR concentrations, read numbers and abundances were formatted and combined with sample metadata using the R packages “tidyverse”, “readxl” and “writexl” (Ooms et al., 2023; Wickham et al., 2023, 2019).

#### 1.6.3 Changes over time

The R package “ggplot2” v0.4.3 (Wickham, 2016) was used to visualize the relative abundance of the two species or their phyla over time at the three locations using all four datasets. The ske/apo-index, the Dia/Dino index of the biomonitoring data, and ddPCR data were visualized over time. A linear trend analysis was performed by calculating the correlation between the ddPCR concentrations and time for both species, utilizing all data points from the three short cores. This was done using Pearson’s correlation.

## 2. Results

### 2.1. Data consistency

The ddPCR concentration measurement of the replicates resulted in consistent concentration per sample, justifying the use of average concentrations. Full data of replicate concentrations are available in the supplements (See Supplementary Tab. S6). The dinoflagellate and diatom communities of the metabarcoding replicates resulted in similar composition except for a few single replicates (see Supplementary Figure S5, Table S5). Biomonitoring data has only been available since the 1980s, and the relative abundance of taxa in the datasets varies dynamically (Figure S6). We found no statistically significant correlation between *A. malmogiense* and *S. marinoi* (R=-0.004, p=0.96) (See Fig. S7).

### 2.2. Correlation analyses

The correlation analysis of four datasets, 1) ddPCR concentration, 2) biomonitoring genus, 3) biomonitoring phylum, and 4) metabarcoding phylum, revealed significant positive correlations, but also distinct differences between the two species, *A. malmogiense* and *S. marinoi*, and their respective datasets targeting different taxonomic levels (Fig. 2).

**Fig. 2:**
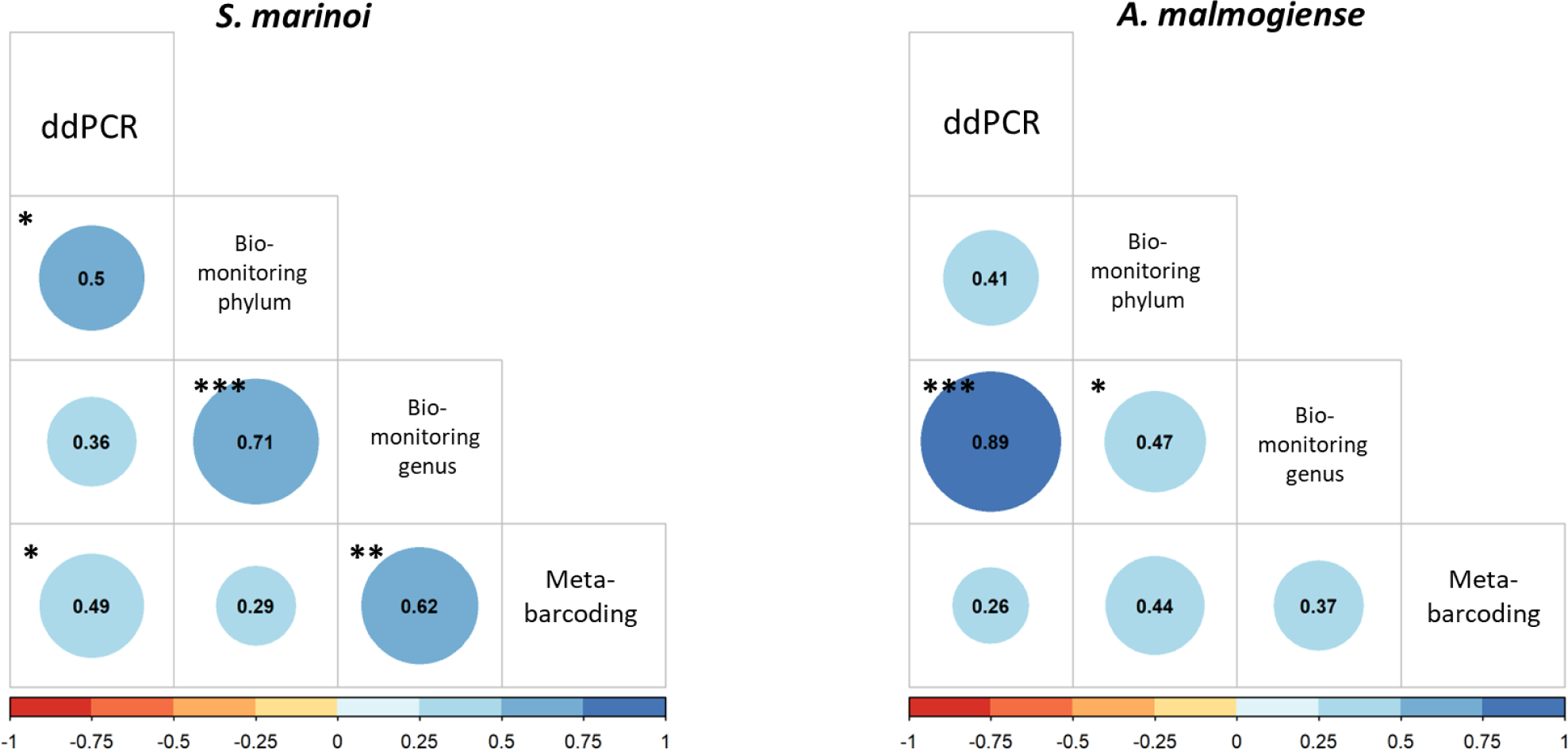
Pearson’s correlation between the four different datasets for *A. malmogiense* and *S. marinoi*: 1) ddPCR concentration, 2) Biomonitoring phylum, 3) Biomonitoring species, 4) Metabarcoding phylum. Correlation coefficients from −1 (red) to 1 (blue) and their significance. *: p<0.05, **: p<0.01, ***: p<0.001. Detailed information in supplementary Table S7.

For the dinoflagellate *A. malmogiense*, we found a significant positive correlation between the measured ddPCR concentration and the biomonitoring data of the genus (R=0.891, p=3.216E-07). However, when considering the correlation between the ddPCR concentration and the biomonitoring at the phylum level, the correlation was weaker (R=0.414, p=0.078). Regarding the metabarcoding read count, dinoflagellates had a weaker correlation between read number and ddPCR concentration (R=0.262, p=0.277). The correlation of metabarcoding read count with biomonitoring relative abundance at the phylum level was stronger (R=0.441, p=0.058).

For the diatom *S. marinoi*, the correlation between the measured ddPCR concentration and the biomonitoring data of the genus was weaker (R=0.363, p=0.127) than for *A. malmogiense*. However, *S. marinoi* has a stronger and significant correlation between the ddPCR concentration and the biomonitoring at the phylum level (R=0.501, p=0.028). Regarding the metabarcoding read count, diatoms had a stronger correlation between read numbers and ddPCR concentrations (R=0.492, p=0.032). Metabarcoding diatom read numbers also had a strong and significant correlation with *Skeletonema* biomonitoring relative abundance at a genus level (R=0.617, p=0.005). However, the correlation of metabarcoding read count with biomonitoring relative abundance at the phylum level was weak (R=0.287, p=0.233).

In summary, there are notable strong correlations between the ddPCR concentrations of both species and the biomonitoring data.

### 2.3 Trends over the past decades and centuries

A linear trend analysis of ddPCR concentration over time reveals a significant positive correlation between the sediment sample age (Year) and the ddPCR concentration of *A. malmogiense*, indicating a substantial increase in its concentration in recent years (R=0.21, p<0.05). Conversely, *S. marinoi* shows a weaker correlation (R=0.09) that is not statistically significant (p=0.366). These findings support the interpretation of a recent shift in dominance from *S. marinoi* to *A. malmogiense*, a trend further corroborated by the segmented regression analysis (Fig. 3, A).

**Fig. 3:**
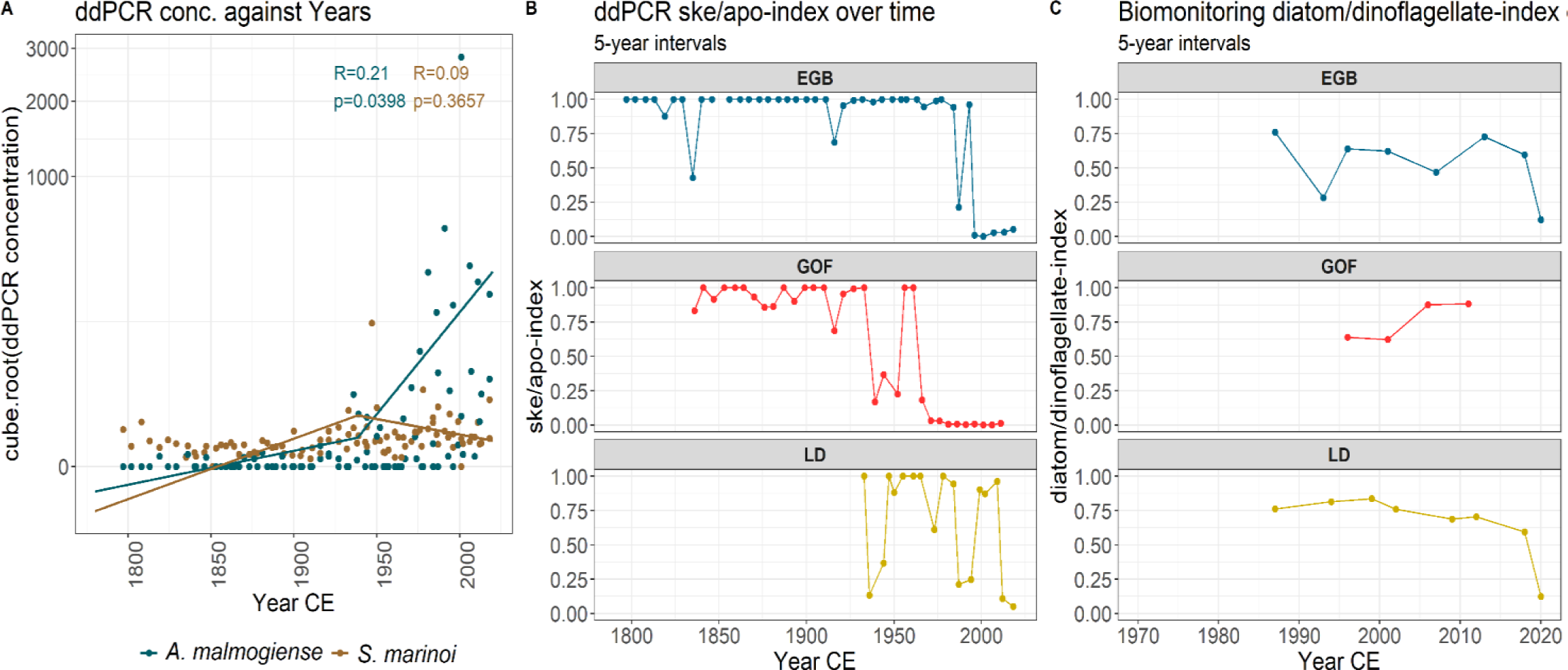
A) Pearson’s correlation between time and ddPCR concentration (cube root transformed) for *A. malmogiense* (blue) and *S. marinoi* (brown); segmented regression. B) ddPCR ske/apo-index over time in five year intervals at all the locations (1798-2018). C) Biomonitoring Dia/Dino-index over time in five year intervals at all three locations (1984-2021).

This shift is further supported by the trends in the ske/apo-index based on the ddPCR data and the Dia/Dino-index based on the biomonitoring data (Fig. 3 B-C), which shows drops in recent years. Notably, however, the biomonitoring based Dia/Dino-index starts at values below 1 during the timespan of monitoring, while the ske/apo-index reaches values of 1 in the early years of the core. This record suggests that the initial shift in dominance - and hence drop in ske/apo-index - occurs earlier than the timespan covered by the biomonitoring data. In the EGB, a first drop can be observed in the ddPCR data around 1980, which then advances into the late 1990s. The Dia/Dino-index shows that the index was around 0.6 in the 1980s, and decreased to 0.12 in recent years. In the GOF, the ske/apo-index and Dia/Dino-index show contradicting dynamics. While the ddPCR results show a first decrease in the 1930s and a second in the 1960s-1970s, the biomonitoring results show an increase since the first data point in the 1980s, though never reaching 1. In the LD, a first decrease in the ske/apo-index can be observed in the 1930s as well, followed by a second decrease in the 1980s and a third decrease around the 2000s. The Dia/Dino-index also shows an ongoing decrease since the 1980s, becoming more rapid in the 2010s.

Overall, the two biomonitoring indices display a positive significant correlation (R=0.39, p=0.012). This supports the use of the molecular ske/apo-index as the ske/apo-index shows a similar trend than the Dia/Dino-index.

### 2.4 Trends over millennia

To evaluate long-term variability and stability of the target species ratio, we extended our investigation into the past. Figure 4 provides an overview of the ddPCR concentrations and the resulting ske/apo-index in the Eastern Gotland Basin (EGB) and the Gulf of Finland (GOF), extending from approximately 8000 years BCE to 2000 years CE. In the EGB, the relationship was highly stable, except for one significant shift in the concentration of *A. malmogiense* around 4500 years BCE and one minor shift around 1500 years BCE. Interestingly, *A. malmogiense* occurred quite frequently in more recent years, but only very seldom before. The GOF, on the other hand, was much less stable. Multiple dominance shifts are observed starting at 2000 years BCE, followed by less pronounced shifts over the next 3000 years, and then again from 1000 years CE to 2000 years CE. Notably, in the GOF, *A. malmogiense* is found more frequently and at higher relative concentrations throughout the record. We visually compare these results to the historical development of the Baltic Sea, focussing on detailed changes in oxygen - through changing XFR Mn/Ti proportions - and atmospheric temperature, and considering overall changes in salinity in the different stages of the Baltic (Fig. 4). The blue line, indicative of air temperature, portrays an overall upward trend despite intermittent fluctuations. The black line shows the dynamics of XRF Mn/Ti proportion at the EGB, and the red bars indicate the main anoxic phases in the central Baltic Sea at approximately 5500 to 4500 BCE, 4000 to 1500 BCE, 500 to 1200 CE and around 2000 CE.

**Fig. 4:**
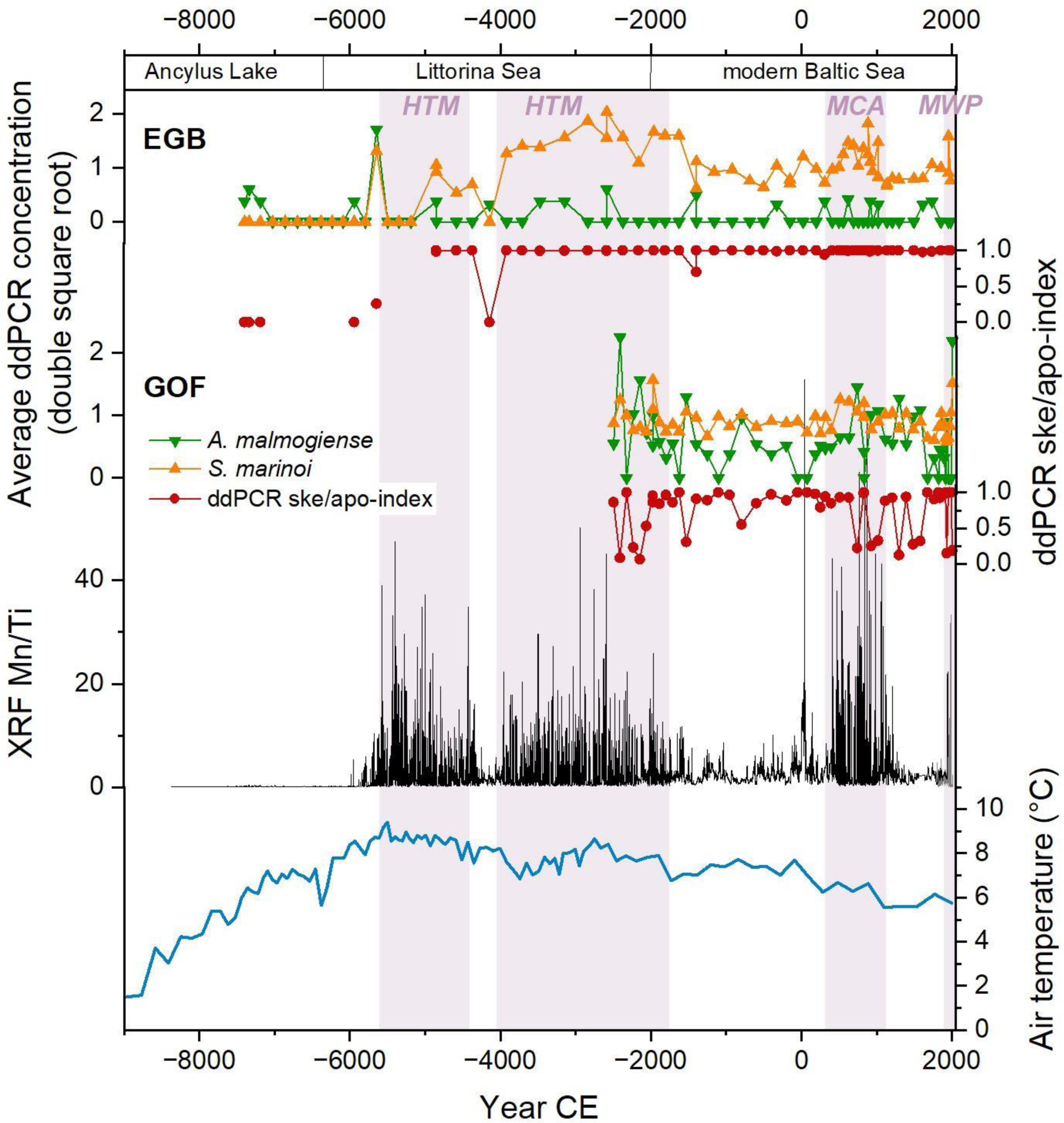
Temporal development of the double square root transformed ddPCR concentration (expressed in copies/µL) at two locations: the Eastern Gotland Basin (EGB) and the Gulf of Finland (GOF). The ddPCR ske/apo-index over time is highlighted in red. The air temperature is shown in blue and based on data from Lake Flarken, Sweden (Seppä et al., 2005). The XRF Mn/Ti data, proxy for oxygen levels, from core EMB262/6-30GC is pin black. Red bars are used to indicate the primary anoxic phases in the central Baltic Sea (Andrén et al., 2011; Rosentau et al., 2017). Main phases of the Baltic Sea are used from Andrén et al. (2011) and Rosentau et al. (2017).

## 3. Discussion

### 3.1. Validation of methods and extension of monitoring records into the past - ske/apo-index

We compared three different methods to investigate diatom and dinoflagellate dominance: available biomonitoring data from ICES (2022), which dates back to the 1980s, DNA metabarcoding of eukaryotes, and species-specific ddPCR reactions. The molecular data showed consistent results across replicates, both in terms of community composition and in quantity of single species reads or ddPCR concentrations. The observed ddPCR concentrations are independent from each other, as evidenced by the lack of observed correlation between *A. malmogiense* and *S. marinoi* concentrations. This independence is crucial as it allows us to avoid attributing changes in species-specific concentrations to broader fluctuations in sedaDNA concentrations. By checking for this, we ensure a more accurate and species-specific trend analysis, thereby enhancing the reliability of our subsequent ske/apo-index analysis.

Metabarcoding demonstrated its capability to detect changes in both phyla and species, although it was not as sensitive as ddPCR to changes in the biomonitoring data when it came to *A. malmogiense* or the diatom phylum. While metabarcoding is effective in reflecting biomonitoring-based dynamics, it is of particular importance to note that it not only focuses on spring bloom organisms crucial for the Dia/Dino index. A broad metabarcoding primer is needed for investigating different phyla, typically resulting in a loss of taxonomic resolution, such that retrieved sequences can often not be identified to species level (Bunse et al., 2016; Haraguchi et al., 2023; Jaanus et al., 2006). In combination with the limited temporal resolution of sediment core samples, which typically include more than a year, this prohibits the recovery of season-specific dynamics. In contrast, targeting specific seasonally distinct species with ddPCR, i.e. spring dinoflagellates and diatoms, such as *A. malmogiense* and *S. marinoi*, restores this seasonal resolution. The retrieved signal from each sample will likely stem from multiple spring blooms, but ignores the other seasons. Quantitative approaches for single species from eDNA are by now well established, and both ddPCR and real time quantitative PCR (qPCR) have been utilized for many years (e.g. Doi et al., 2015; Takahara et al., 2012). In some qPCR studies (Knudsen et al., 2019; Spear et al., 2015), no clear links between DNA concentrations and biomass estimates were established, whereas ddPCR is suggested to be more accurate, especially at low DNA concentrations (Doi et al., 2015), where it shows higher detection rates compared to qPCR, possibly due to its resistance to inhibition (Dingle et al., 2013; Rački et al., 2014).

The two species were found across many segments of the core, but the patterns of appearance differed: In the case of *A*. *malmogiense*, changes in concentration were particularly pronounced, as many samples were negative. In *S. marinoi* we observed more gradual changes between samples. An explanation for this could be different taphonomy of the sedaDNA of the two species, as the fate of the two target species differs substantially once the bloom ceases. While diatoms such as *S. marinoi* primarily sink to the sea floor, where many of them disintegrate and some transform into a stage of physiological resting, cells of *A. malmogiense* partly demineralize in the euphotic zone (Wasmund et al., 2017) or encyst. When deposited in anoxic sediment after the bloom, their DNA is retained in a resistant resting propagule (Spilling et al., 2018). *S. marinoi* not only occurs in the spring bloom, but throughout the year, with a second smaller bloom in September, as well. Considering this, the probability of finding *S. marinoi* DNA in the sediment core is generally higher than of encountering *A. malmogiense* (Saravanan and Godhe, 2010). The findings of *A. malmogiense* DNA in the sediment thus point to a significant increase in dinoflagellates in the study area.

Our study underscores the potential of molecular tools in decoding historical events and long-term environmental trend dynamics through sedaDNA, and including organismal signals that are not present in the visual fossil record. Here, we use simple species-specific reactions, including a quantitative aspect, i.e. highly sensitive ddPCR. This can simplify data in comparison to metabarcoding or metagenomics (Gielings et al., 2021; Nguyen et al., 2023) and our analyses suggest that it can deliver reliable information on ecological changes by narrowing the focus on ecologically relevant key species that are used as proxies and in indices for ecosystem status. Such simple approaches could be implemented in monitoring and management relatively easily. While our current research has primarily centered around duplex ddPCR, it is crucial to acknowledge the emerging potential of multiplex ddPCR as a promising approach for the future. By extending the available fluorescent channels to accommodate up to six distinct targets, researchers can now detect and quantify multiple targets simultaneously within a single assay (Wainman et al., 2024). For instance, in the context of the molecular Dia/Dino-index, it becomes feasible to select three target spring bloom species per taxa, thereby creating a more complex index that does not lose temporal resolution. In the case of the molecular Dia/Dino-index we test here, such assays would allow to include more than the two model species used in this study. These additional assays would enhance our understanding of functional diversity across different taxa. Specifically, during the Baltic spring bloom, both diatoms and dinoflagellates play a crucial role. They consist of several key species, each possessing distinct traits that contribute to the index (Haraguchi et al., 2023). In the central and northern parts of the Baltic Sea, the dinoflagellate community is dominated by three cold-water species: *Apocalathium malmogiense* (Peridiniales), *Biecheleria baltica* (Suessiales), and *Gymnodinium corollarium* (Gymnodiniales) (Kremp et al., 2009). These species, although similar in habit, exhibit significant phylogenetic and ecological differences. By incorporating these diverse species through appropriate assays, we can enhance the complexity and informative value of the index.

The possibility to retrieve paleoecological records from a number of locations, i.e., from different regions of the Baltic Sea, as done here, enables a more precise validation of local ecological indices and the establishment of region specific values for good ecological status (GES). For the original Dia/Dino index, Wasmund (2017) investigated historical count data from the first half of the 20th century, to verify that the index was stable prior to eutrophication and to establish regional values for GES. The data indeed indicated a pre-eutrophication stability, and demonstrated that values for GES are not uniform across the Baltic Sea. They suggested regionally different values for GES, which are currently only established for the southern and central Baltic Sea (HELCOM, 2023), not for e.g., the GOF.

Our sediment core analyses indicate that the GOF was much less stable regarding the dominance of diatoms and dinoflagellates on millennial scales (Fig. 4). However, Ske/Apo-index values for the past century demonstrate that the current anthropogenic eutrophication is captured very well by changes in this index (Fig. 3). The instability of the index in the GOF, as well as the historical drops in the relatively stable EGB record (Fig. 4) rather points to the spring bloom composition being sensitive to both eutrophication and warming. Overall, the data highlights the importance of defining a GES for each region, as the optimal index value is significantly influenced by the unique environmental conditions in each region.

### 3.2. Trends over the past decades and centuries

Monitoring records suggest that diatoms were more abundant than dinoflagellates in the past, making diatom dominance the preferred status, reflecting a GES of the Baltic Sea (Wasmund, 2017; Wasmund et al., 2017). According to Wasmund et al. (2017), the Dia/Dino-index points to a regime shift at the end of the 1980s in the central Baltic Sea. This is also recorded in our data using the ddPCR-based ske/apo-index, but in particular the GOF and the LD show earlier drops in the dominance reversal during the 20th century (Fig. 3). In the GOF, Ske/Apo-Index values have remained low since the 1960s, and in both the LD and the GOF, an initial massive drop is recorded in the first half of the century, slightly predating the Second World War, which aligns with the introduction of artificial fertilizers in the early 20th century (Treitel, 2015). Results from the EGB and the LD show shifts around the 1980s when eutrophication was nearing its peak in the Baltic Sea.

The massive eutrophication in the open Baltic Sea started after the Second World War and reached its peak in nutrient concentrations (phosphorus and nitrogen) in the 1980s and 1990s (Gustafsson et al., 2012; HELCOM, 2021; Murray et al., 2019). Murray et al. (2019) conducted a comprehensive study, modeling the past, present, and future eutrophication status of the Baltic Sea. They identified a shift from a healthy state to a state with eutrophication issues occurring in the late 1950s and early 1960s. Our results align with this timeline, showing a noticeable drop in the ske/apo-index during the 1980s and 1990s. Interestingly, we also observed a similar drop in the 1950s and 1960s specifically in the GOF. Consequently, our data suggests that changes in the trophic status may trigger species-level responses, particularly evident in the increased concentrations of *A. malmogiense*. Hence, the ske/apo-index built with sedaDNA ddPCR concentration is a promising tool to estimate changes in the trophic state of the past beyond the instrumental period.

### 3.3 Trends over millennia

Our study furthermore elucidates millennial dynamics of the key phytoplankton species dominance by examining long cores from the Eastern Gotland Basin (EGB) and the Gulf of Finland (GOF). These cover time periods of ca. 7000 years and can be compared to environmental data covering ca. 11,000 years. Looking at the long timeline allows us to better explore the influence of various factors on the phytoplankton species on long time scales, including eutrophication, large-scale salinity change, and temperature. A previous study indicates that an increase in temperature can boost the abundance of dinoflagellates, but not of diatoms (Bi et al., 2021). Elevated water temperatures have been observed to trigger earlier blooms of diatoms and dinoflagellates, while also reducing the magnitude of diatom blooms (Hjerne et al., 2019). This suggests that a gradual temporal increase in temperature could potentially foster dinoflagellate dominance. A study examining the impact of salinity on dinoflagellates and diatoms using *Alexandrium ostenfeldii* (another common Baltic Sea dinoflagellate) and *S. marinoi* found that *A. ostenfeldii* exhibited significantly higher trait plasticity than *S. marinoi* (Orizar and Lewandowska, 2022). Our two target organisms, *S. marinoi* and *A. malmogiense*, are both considered to be well adapted to brackish salinities and can grow well across the entire Baltic Sea salinity gradient (Logares et al., 2007; Sjöqvist et al., 2015). Nonetheless, salinity shifts may alter the competitive dynamics between the two to some extent.

Overall, temporal dynamics of the two species are not fully congruent across the examined cores. In the EGB core, *Skeletonema* dominance prior to the 20th century was mostly stable, with only two major shifts in the ske/apo-index observed, at about 4500 years BCE and 1500 years BCE respectively. Conversely, the GOF exhibits a more dynamic history with several shifts among the two species across millennia. We find that both species are present in many samples of the GOF, while the EGB experiences several explicit drops of *Skeletonema*, coupled with sporadic occurrences of *A. malmogiense*. The presence of *A. malmogiense* in the recent sediment layers of the EGB, in comparison to its lower detection likelihood across millennia, suggests that the recent trends are indeed specific to the anthropogenic changes of the past century, but similar shifts were induced by previous ecosystem perturbations. Taken together, the records confirm the trends of high stability of the ske/apo-index in the EGB and a higher variability in the GOF that was found across the past century

The few shifts that do occur in the EGB record are associated with periods of ecosystem change: We note a brief dominance of *A. malmogiense* around 4500 BCE, within the Holocene Thermal Maximum (HTM; 6000-4000 BCE), but coinciding with a break from anoxic phases. At 1500 BCE, a decrease in the index again aligns with more oxygen in the deep water column. A coeval drop of the index occurs in the GOF core after a more stable time period. This initiates a series of index drops in the GOF in the period of the Medieval Climate Anomaly (MCA; 950-1250 CE). During this time, water was warmer at all depths, accompanied by an extensive hypoxia (Andrén et al., 2020; Funkey et al., 2014; Zillén et al., 2008). In the EGB, *S. marinoi* was highly abundant during this period, both in our data and in previously published count data from another core (Andrén et al., 2020), and *A. malmogiense* also appeared more frequently in the ddPCR. In the GOF, this increase of *A. malmogiense* was more pronounced, while *S. marinoi* ddPCR concentrations remained similar-resulting in a relative increase of the dinoflagellate and drops in the ske/apo-index.

These prominent index shifts, as well as smaller variations in the concentrations of *S. marinoi* and *A. malmogiense* DNA are not linked to a particular condition among the abiotic factors oxygen level, temperature and salinity. Instead, our observations indicate that any major change in these parameters can elicit a response in the ske/apo-index, the magnitude of which can vary. The long-term data of the two species suggests that *A. malmogiense* is competitively superior in times of ecosystem instability, while *S. marinoi* populations drastically decline. This decline, however, is not long-lasting, and the initial dominance relation is re-established, apparently without a return to previous conditions. *S. marinoi* populations have apparently adapted to new ecological conditions, but with a slight delay in comparison to *A. malmogiense*.

At the same time, the observed difference in stability of the ske/apo-index between the two sites is striking. The stability across millennia in the EGB in conjunction with the late drop of the ske/apo-index compared to the GOF and LD in the past century, could be considered an indication of a highly resilient ecosystem in the Eastern Gotland Basin and a high tolerance to environmental change. However, the variability in the GOF is mostly caused by a high prevalence of *A. malmogiense* throughout the core, and the less striking dominance of *S. marinoi*. This could also be a reflection of the specific adaptation of *A. malmogiense* to cold-water habitats (Logares et al., 2007). The species is often associated with sea ice in coastal waters, and the conditions in the GOF are generally more conducive to its establishment. This underscores the complexity of the dynamics of the Baltic Sea marine ecosystem and highlights the need for further research.

## 4. Conclusion

The application of ddPCR allowed us to identify species-level responses to ecosystem perturbations caused by anthropogenic pressures, such as eutrophication. Our analysis resulted in a new index, the ske/apo-index, providing an attainable historic perspective on shifts in species dominance over time. This methodology enabled us to develop a time series extending back multiple millennia that is directly comparable to results from modern biomonitoring, which only began in the 1980s. Our findings indicate that certain areas of the Baltic Sea, but not all, showed an extremely stable relationship of diatoms and dinoflagellate across millennia, that was altered only very seldomly, as in the 20th century. This confirms that an index based on changes in diatom to dinoflagellate dominance can be used in these areas to infer GES, and demonstrates the utility of molecular paleoecological analyses to validate simple indices for modern biomonitoring. The results from the analyses across the past century furthermore pinpoint the timing of initial 20th century change in the Baltic Sea, which substantially predates regular biomonitoring programs. This shift occurred around the Second World War era or slightly earlier, possibly aligning with the introduction of artificial fertilizers in the early 20th century (Treitel et al. 2015).

## Author contributions

**Alexandra Schmidt:** conceptualization (lead), data generation (lead), analyses (lead); data visualization (lead), data interpretation (lead), writing - original draft (lead); **Juliane Romahn:** data generation, analyses (supported); **Elinor Andrén:** conceptualization (supported), writing - review and editing (supported). **Anke Kremp:** conceptualization (supported), sampling (lead), writing - review and editing (supported), funding acquisition; **Jérôme Kaiser:** data generation, analyses, data visualization, data interpretation (supported), writing - review and editing (supported); **Helge W. Arz:** data generation, analyses, data visualization, data interpretation (supported), writing - review and editing (supported); **Olaf Dellwig:** data generation, analyses, data interpretation (supported), writing - review and editing (supported); **Miklós Bálint:** conceptualization (supported), analyses (supported), funding acquisition, writing - review and editing (supported); **Laura S. Epp:** conceptualization (lead), sampling (lead), writing - review and editing, supervision, project administration, funding acquisition

## Acknowledgements

This study was funded by the K314/2020 grant of the Collaborative Excellence Programme of the Leibniz Association. MB is supported by the Hessian Ministry of Higher Education, Research and the Arts through the LOEWE Centre for Translational Biodiversity Genomics. We thank Matthias Moros for performing radionuclide measurements of the GOF MUC. For help with the initial primer design we thank Ilayda Ayyuce Kocak and Kim Isabelle Mayer. For setting up the ancient DNA clean laboratory in Frankfurt where all DNA extractions were done by AS and JR, we thank Leonie Schardt. EA was financially supported by a grant provided by The Foundation for Baltic and East European Studies (42/19).

## Data availability statement

Unix shell scripts, R scripts and data are available at GitHub (https://github.com/Alex-132/EI_paper_rep).

