## Supplement_file for "Decoding the Baltic Sea’s Past and Present: A simple Molecular Index for Ecosystem Assessment"

#### Supplementary Data

### Sampling data

#### Samples

Table S1 provides information about the core samples taken during expedition EGB262 and their locations in the Baltic Sea. Two samples were taken from the Eastern Gotland Basin (EGB) at a depth of 241 meters below sea level (m bsl), and two from the Gulf of Finland (GOF) at a depth of 81.1 m bsl. Another sample was taken from Landsort Deep (LD) at a depth of 436 m bsl. The core identifiers, exact latitudes, and longitudes are also provided for each sample.

**Table S1: Core sample location and details of EMB262. The core identifiers, exact locations (latitude and longitude), and water depths are provided. The locations include the Eastern Gotland Basin (EGB), the Gulf of Finland (GOF), and Landsort Deep (LD). The water depth is measured in meters below sea level (m bsl).**

| Core | Location | Abbreviation | Latitude [°N] | Longditude [°E] | water depth [m bsl] |
|---|---|---|---|---|---|
| EMB262.6.28.MUC | Eastern Gotland Basin | EGB | 57°17.004' | 020°07.244' | 241 |
| EM262B.12.2.MUC | Gulf of Finland | GOF | 59°34.443' | 023°36.461' | 81,1 |
| EMB262.12.3.GC | Gulf of Finland | GOF | 59°34.450' | 023°36.455' | 81,1 |
| EMB262.6.30.GC | Eastern Gotland Basin | EGB | 57°17.022' | 020°07.285' | 241 |
| EMB262.13.8.MUC | Landsort Deep | LD | 58°38.391' | 018°15.997' | 436 |

A detailed table (Table S2) that provides comprehensive information about the multi corer samples collected. Each sample is uniquely identified and includes data such as the trip identifier, station number, event number, gear type, sample number, tag, depth, age, latitude, longitude, and water depth at the collection site.

**Table S2: Detailed information of the multi corer (MUC) samples collected during expedition EMB262 (Kremp et al., 2021) in April 2021 onboard the research vessel Elisabeth Mann Borgese.**

| trip | station | event | gear | sample | tag | depth | age | latitude [°N] | longitude [°E] | water depth [m bsl] |
|---|---|---|---|---|---|---|---|---|---|---|
| EMB262 | 3 | 10 | MUC | 1 | EMB262_3_10_MUC_1 | 1 | 0 | 55°15.044' | 015°58.926' | 89,9 |
| EMB262 | 3 | 10 | MUC | 2 | EMB262_3_10_MUC_2 | 2 | 0 | 55°15.044' | 015°58.926' | 89,9 |
| EMB262 | 3 | 10 | MUC | 3 | EMB262_3_10_MUC_3 | 3 | 0 | 55°15.044' | 015°58.926' | 89,9 |
| EMB262 | 3 | 10 | MUC | 4 | EMB262_3_10_MUC_4 | 4 | 0 | 55°15.044' | 015°58.926' | 89,9 |
| EMB262 | 3 | 10 | MUC | 5 | EMB262_3_10_MUC_5 | 5 | 0 | 55°15.044' | 015°58.926' | 89,9 |
| EMB262 | 3 | 10 | MUC | 6 | EMB262_3_10_MUC_6 | 6 | 0 | 55°15.044' | 015°58.926' | 89,9 |
| EMB262 | 3 | 10 | MUC | 7 | EMB262_3_10_MUC_7 | 7 | 0 | 55°15.044' | 015°58.926' | 89,9 |
| EMB262 | 3 | 10 | MUC | 8 | EMB262_3_10_MUC_8 | 8 | 0 | 55°15.044' | 015°58.926' | 89,9 |
| EMB262 | 3 | 10 | MUC | 9 | EMB262_3_10_MUC_9 | 9 | 0 | 55°15.044' | 015°58.926' | 89,9 |
| EMB262 | 3 | 10 | MUC | 10 | EMB262_3_10_MUC_10 | 10 | 0 | 55°15.044' | 015°58.926' | 89,9 |
| EMB262 | 3 | 10 | MUC | 11 | EMB262_3_10_MUC_11 | 11 | 0 | 55°15.044' | 015°58.926' | 89,9 |
| EMB262 | 3 | 10 | MUC | 12 | EMB262_3_10_MUC_12 | 12 | 0 | 55°15.044' | 015°58.926' | 89,9 |
| EMB262 | 3 | 10 | MUC | 13 | EMB262_3_10_MUC_13 | 13 | 0 | 55°15.044' | 015°58.926' | 89,9 |
| EMB262 | 3 | 10 | MUC | 14 | EMB262_3_10_MUC_14 | 14 | 0 | 55°15.044' | 015°58.926' | 89,9 |
| EMB262 | 3 | 10 | MUC | 15 | EMB262_3_10_MUC_15 | 15 | 0 | 55°15.044' | 015°58.926' | 89,9 |
| EMB262 | 3 | 10 | MUC | 16 | EMB262_3_10_MUC_16 | 16 | 0 | 55°15.044' | 015°58.926' | 89,9 |
| EMB262 | 3 | 10 | MUC | 17 | EMB262_3_10_MUC_17 | 17 | 0 | 55°15.044' | 015°58.926' | 89,9 |
| EMB262 | 3 | 10 | MUC | 18 | EMB262_3_10_MUC_18 | 18 | 0 | 55°15.044' | 015°58.926' | 89,9 |
| EMB262 | 3 | 10 | MUC | 19 | EMB262_3_10_MUC_19 | 19 | 0 | 55°15.044' | 015°58.926' | 89,9 |
| EMB262 | 3 | 10 | MUC | 20 | EMB262_3_10_MUC_20 | 20 | 0 | 55°15.044' | 015°58.926' | 89,9 |
| EMB262 | 3 | 10 | MUC | 21 | EMB262_3_10_MUC_21 | 21 | 0 | 55°15.044' | 015°58.926' | 89,9 |
| EMB262 | 3 | 10 | MUC | 22 | EMB262_3_10_MUC_22 | 22 | 0 | 55°15.044' | 015°58.926' | 89,9 |
| EMB262 | 3 | 10 | MUC | 23 | EMB262_3_10_MUC_23 | 23 | 0 | 55°15.044' | 015°58.926' | 89,9 |
| EMB262 | 3 | 10 | MUC | 24 | EMB262_3_10_MUC_24 | 24 | 0 | 55°15.044' | 015°58.926' | 89,9 |
| EMB262 | 3 | 10 | MUC | 25 | EMB262_3_10_MUC_25 | 25 | 0 | 55°15.044' | 015°58.926' | 89,9 |
| EMB262 | 3 | 10 | MUC | 26 | EMB262_3_10_MUC_26 | 26 | 0 | 55°15.044' | 015°58.926' | 89,9 |
| EMB262 | 3 | 10 | MUC | 27 | EMB262_3_10_MUC_27 | 27 | 0 | 55°15.044' | 015°58.926' | 89,9 |
| EMB262 | 3 | 10 | MUC | 28 | EMB262_3_10_MUC_28 | 28 | 0 | 55°15.044' | 015°58.926' | 89,9 |
| EMB262 | 3 | 10 | MUC | 29 | EMB262_3_10_MUC_29 | 29 | 0 | 55°15.044' | 015°58.926' | 89,9 |
| EMB262 | 3 | 10 | MUC | 30 | EMB262_3_10_MUC_30 | 31 | 0 | 55°15.044' | 015°58.926' | 89,9 |
| EMB262 | 3 | 10 | MUC | 31 | EMB262_3_10_MUC_31 | 31 | 0 | 55°15.044' | 015°58.926' | 89,9 |
| EMB262 | 3 | 10 | MUC | 32 | EMB262_3_10_MUC_32 | 32 | 0 | 55°15.044' | 015°58.926' | 89,9 |
| EMB262 | 3 | 10 | MUC | 33 | EMB262_3_10_MUC_33 | 33 | 0 | 55°15.044' | 015°58.926' | 89,9 |
| EMB262 | 3 | 10 | MUC | 34 | EMB262_3_10_MUC_34 | 34 | 0 | 55°15.044' | 015°58.926' | 89,9 |
| EMB262 | 3 | 10 | MUC | 35 | EMB262_3_10_MUC_35 | 35 | 0 | 55°15.044' | 015°58.926' | 89,9 |
| EMB262 | 3 | 10 | MUC | 36 | EMB262_3_10_MUC_36 | 36 | 0 | 55°15.044' | 015°58.926' | 89,9 |
| EMB262 | 3 | 10 | MUC | 37 | EMB262_3_10_MUC_37 | 37 | 0 | 55°15.044' | 015°58.926' | 89,9 |
| EMB262 | 3 | 10 | MUC | 38 | EMB262_3_10_MUC_38 | 38 | 0 | 55°15.044' | 015°58.926' | 89,9 |
| EMB262 | 3 | 10 | MUC | 39 | EMB262_3_10_MUC_39 | 39 | 0 | 55°15.044' | 015°58.926' | 89,9 |
| EMB262 | 6 | 28 | MUC | 1 | EMB262_6_28_MUC_1 | 1 | 2018 | 57°17.004' | 020°07.244' | 241 |
| EMB262 | 6 | 28 | MUC | 3 | EMB262_6_28_MUC_3 | 3 | 2013 | 57°17.004' | 020°07.244' | 241 |
| EMB262 | 6 | 28 | MUC | 5 | EMB262_6_28_MUC_5 | 5 | 2007 | 57°17.004' | 020°07.244' | 241 |

| | | | | | | | | | | |
|---|---|---|---|---|---|---|---|---|---|---|
| EMB262 | 6 | 28 | MUC | 7 | EMB262_6_28_MUC_7 | 7 | 2001 | 57°17.004' | 020°07.244' | 241 |
| EMB262 | 6 | 28 | MUC | 9 | EMB262_6_28_MUC_9 | 9 | 1996 | 57°17.004' | 020°07.244' | 241 |
| EMB262 | 6 | 28 | MUC | 11 | EMB262_6_28_MUC_11 | 11 | 1993 | 57°17.004' | 020°07.244' | 241 |
| EMB262 | 6 | 28 | MUC | 12 | EMB262_6_28_MUC_12 | 12 | 1990 | 57°17.004' | 020°07.244' | 241 |
| EMB262 | 6 | 28 | MUC | 13 | EMB262_6_28_MUC_13 | 13 | 1987 | 57°17.004' | 020°07.244' | 241 |
| EMB262 | 6 | 28 | MUC | 14 | EMB262_6_28_MUC_14 | 14 | 1984 | 57°17.004' | 020°07.244' | 241 |
| EMB262 | 6 | 28 | MUC | 15 | EMB262_6_28_MUC_15 | 15 | 1981 | 57°17.004' | 020°07.244' | 241 |
| EMB262 | 6 | 28 | MUC | 16 | EMB262_6_28_MUC_16 | 16 | 1977 | 57°17.004' | 020°07.244' | 241 |
| EMB262 | 6 | 28 | MUC | 17 | EMB262_6_28_MUC_17 | 17 | 1974 | 57°17.004' | 020°07.244' | 241 |
| EMB262 | 6 | 28 | MUC | 18 | EMB262_6_28_MUC_18 | 18 | 1970 | 57°17.004' | 020°07.244' | 241 |
| EMB262 | 6 | 28 | MUC | 19 | EMB262_6_28_MUC_19 | 19 | 1967 | 57°17.004' | 020°07.244' | 241 |
| EMB262 | 6 | 28 | MUC | 20 | EMB262_6_28_MUC_20 | 20 | 1963 | 57°17.004' | 020°07.244' | 241 |
| EMB262 | 6 | 28 | MUC | 21 | EMB262_6_28_MUC_21 | 21 | 1960 | 57°17.004' | 020°07.244' | 241 |
| EMB262 | 6 | 28 | MUC | 22 | EMB262_6_28_MUC_22 | 22 | 1957 | 57°17.004' | 020°07.244' | 241 |
| EMB262 | 6 | 28 | MUC | 23 | EMB262_6_28_MUC_23 | 23 | 1954 | 57°17.004' | 020°07.244' | 241 |
| EMB262 | 6 | 28 | MUC | 24 | EMB262_6_28_MUC_24 | 24 | 1949 | 57°17.004' | 020°07.244' | 241 |
| EMB262 | 6 | 28 | MUC | 25 | EMB262_6_28_MUC_25 | 25 | 1943 | 57°17.004' | 020°07.244' | 241 |
| EMB262 | 6 | 28 | MUC | 26 | EMB262_6_28_MUC_26 | 26 | 1938 | 57°17.004' | 020°07.244' | 241 |
| EMB262 | 6 | 28 | MUC | 27 | EMB262_6_28_MUC_27 | 27 | 1932 | 57°17.004' | 020°07.244' | 241 |
| EMB262 | 6 | 28 | MUC | 28 | EMB262_6_28_MUC_28 | 28 | 1927 | 57°17.004' | 020°07.244' | 241 |
| EMB262 | 6 | 28 | MUC | 29 | EMB262_6_28_MUC_29 | 29 | 1921 | 57°17.004' | 020°07.244' | 241 |
| EMB262 | 6 | 28 | MUC | 30 | EMB262_6_28_MUC_30 | 30 | 1916 | 57°17.004' | 020°07.244' | 241 |
| EMB262 | 6 | 28 | MUC | 31 | EMB262_6_28_MUC_31 | 31 | 1911 | 57°17.004' | 020°07.244' | 241 |
| EMB262 | 6 | 28 | MUC | 32 | EMB262_6_28_MUC_32 | 32 | 1905 | 57°17.004' | 020°07.244' | 241 |
| EMB262 | 6 | 28 | MUC | 33 | EMB262_6_28_MUC_33 | 33 | 1900 | 57°17.004' | 020°07.244' | 241 |
| EMB262 | 6 | 28 | MUC | 34 | EMB262_6_28_MUC_34 | 34 | 1894 | 57°17.004' | 020°07.244' | 241 |
| EMB262 | 6 | 28 | MUC | 35 | EMB262_6_28_MUC_35 | 35 | 1889 | 57°17.004' | 020°07.244' | 241 |
| EMB262 | 6 | 28 | MUC | 36 | EMB262_6_28_MUC_36 | 36 | 1884 | 57°17.004' | 020°07.244' | 241 |
| EMB262 | 6 | 28 | MUC | 37 | EMB262_6_28_MUC_37 | 37 | 1878 | 57°17.004' | 020°07.244' | 241 |
| EMB262 | 6 | 28 | MUC | 38 | EMB262_6_28_MUC_38 | 38 | 1873 | 57°17.004' | 020°07.244' | 241 |
| EMB262 | 6 | 28 | MUC | 39 | EMB262_6_28_MUC_39 | 39 | 1867 | 57°17.004' | 020°07.244' | 241 |
| EMB262 | 6 | 28 | MUC | 40 | EMB262_6_28_MUC_40 | 40 | 1862 | 57°17.004' | 020°07.244' | 241 |
| EMB262 | 6 | 28 | MUC | 41 | EMB262_6_28_MUC_41 | 41 | 1856 | 57°17.004' | 020°07.244' | 241 |
| EMB262 | 6 | 28 | MUC | 42 | EMB262_6_28_MUC_42 | 42 | 1851 | 57°17.004' | 020°07.244' | 241 |
| EMB262 | 6 | 28 | MUC | 43 | EMB262_6_28_MUC_43 | 43 | 1846 | 57°17.004' | 020°07.244' | 241 |
| EMB262 | 6 | 28 | MUC | 44 | EMB262_6_28_MUC_44 | 44 | 1840 | 57°17.004' | 020°07.244' | 241 |
| EMB262 | 6 | 28 | MUC | 45 | EMB262_6_28_MUC_45 | 45 | 1835 | 57°17.004' | 020°07.244' | 241 |
| EMB262 | 6 | 28 | MUC | 46 | EMB262_6_28_MUC_46 | 46 | 1829 | 57°17.004' | 020°07.244' | 241 |
| EMB262 | 6 | 28 | MUC | 47 | EMB262_6_28_MUC_47 | 47 | 1824 | 57°17.004' | 020°07.244' | 241 |
| EMB262 | 6 | 28 | MUC | 48 | EMB262_6_28_MUC_48 | 48 | 1819 | 57°17.004' | 020°07.244' | 241 |
| EMB262 | 6 | 28 | MUC | 49 | EMB262_6_28_MUC_49 | 49 | 1813 | 57°17.004' | 020°07.244' | 241 |
| EMB262 | 6 | 28 | MUC | 50 | EMB262_6_28_MUC_50 | 50 | 1808 | 57°17.004' | 020°07.244' | 241 |
| EMB262 | 6 | 28 | MUC | 51 | EMB262_6_28_MUC_51 | 51 | 1802 | 57°17.004' | 020°07.244' | 241 |
| EMB262 | 6 | 28 | MUC | 52 | EMB262_6_28_MUC_52 | 52 | 1797 | 57°17.004' | 020°07.244' | 241 |
| EMB261 | 12 | 2 | MUC | 2 | EMB261_12_2_MUC_2 | 1 | 2016 | 59°34.443’ | 023°36.461’ | 81,1 |
| EMB262 | 12 | 2 | MUC | 2 | EMB262_12_2_MUC_2 | 2 | 2011 | 59°34.443’ | 023°36.461’ | 81,1 |
| EMB262 | 12 | 2 | MUC | 3 | EMB262_12_2_MUC_3 | 3 | 2006 | 59°34.443’ | 023°36.461’ | 81,1 |

| | | | | | | | | | | |
|---|---|---|---|---|---|---|---|---|---|---|
| EMB262 | 12 | 2 | MUC | 4 | EMB262_12_2_MUC_4 | 4 | 2001 | 59°34.443’ | 023°36.461’ | 81,1 |
| EMB262 | 12 | 2 | MUC | 5 | EMB262_12_2_MUC_5 | 5 | 1996 | 59°34.443’ | 023°36.461’ | 81,1 |
| EMB262 | 12 | 2 | MUC | 6 | EMB262_12_2_MUC_6 | 6 | 1991 | 59°34.443’ | 023°36.461’ | 81,1 |
| EMB262 | 12 | 2 | MUC | 7 | EMB262_12_2_MUC_7 | 7 | 1986 | 59°34.443’ | 023°36.461’ | 81,1 |
| EMB262 | 12 | 2 | MUC | 8 | EMB262_12_2_MUC_8 | 8 | 1981 | 59°34.443’ | 023°36.461’ | 81,1 |
| EMB262 | 12 | 2 | MUC | 9 | EMB262_12_2_MUC_9 | 9 | 1976 | 59°34.443’ | 023°36.461’ | 81,1 |
| EMB262 | 12 | 2 | MUC | 10 | EMB262_12_2_MUC_10 | 10 | 1971 | 59°34.443’ | 023°36.461’ | 81,1 |
| EMB262 | 12 | 2 | MUC | 11 | EMB262_12_2_MUC_11 | 11 | 1966 | 59°34.443’ | 023°36.461’ | 81,1 |
| EMB262 | 12 | 2 | MUC | 12 | EMB262_12_2_MUC_12 | 12 | 1961 | 59°34.443’ | 023°36.461’ | 81,1 |
| EMB262 | 12 | 2 | MUC | 13 | EMB262_12_2_MUC_13 | 13 | 1956 | 59°34.443’ | 023°36.461’ | 81,1 |
| EMB262 | 12 | 2 | MUC | 14 | EMB262_12_2_MUC_14 | 14 | 1952 | 59°34.443’ | 023°36.461’ | 81,1 |
| EMB262 | 12 | 2 | MUC | 15 | EMB262_12_2_MUC_15 | 15 | 1950 | 59°34.443’ | 023°36.461’ | 81,1 |
| EMB262 | 12 | 2 | MUC | 16 | EMB262_12_2_MUC_16 | 16 | 1944 | 59°34.443’ | 023°36.461’ | 81,1 |
| EMB262 | 12 | 2 | MUC | 17 | EMB262_12_2_MUC_17 | 17 | 1939 | 59°34.443’ | 023°36.461’ | 81,1 |
| EMB262 | 12 | 2 | MUC | 18 | EMB262_12_2_MUC_18 | 18 | 1933 | 59°34.443’ | 023°36.461’ | 81,1 |
| EMB262 | 12 | 2 | MUC | 19 | EMB262_12_2_MUC_19 | 19 | 1927 | 59°34.443’ | 023°36.461’ | 81,1 |
| EMB262 | 12 | 2 | MUC | 20 | EMB262_12_2_MUC_20 | 20 | 1921 | 59°34.443’ | 023°36.461’ | 81,1 |
| EMB262 | 12 | 2 | MUC | 21 | EMB262_12_2_MUC_21 | 21 | 1916 | 59°34.443’ | 023°36.461’ | 81,1 |
| EMB262 | 12 | 2 | MUC | 22 | EMB262_12_2_MUC_22 | 22 | 1910 | 59°34.443’ | 023°36.461’ | 81,1 |
| EMB262 | 12 | 2 | MUC | 23 | EMB262_12_2_MUC_23 | 23 | 1904 | 59°34.443’ | 023°36.461’ | 81,1 |
| EMB262 | 12 | 2 | MUC | 24 | EMB262_12_2_MUC_24 | 24 | 1899 | 59°34.443’ | 023°36.461’ | 81,1 |
| EMB262 | 12 | 2 | MUC | 25 | EMB262_12_2_MUC_25 | 25 | 1893 | 59°34.443’ | 023°36.461’ | 81,1 |
| EMB262 | 12 | 2 | MUC | 26 | EMB262_12_2_MUC_26 | 26 | 1887 | 59°34.443’ | 023°36.461’ | 81,1 |
| EMB262 | 12 | 2 | MUC | 27 | EMB262_12_2_MUC_27 | 27 | 1881 | 59°34.443’ | 023°36.461’ | 81,1 |
| EMB262 | 12 | 2 | MUC | 28 | EMB262_12_2_MUC_28 | 28 | 1876 | 59°34.443’ | 023°36.461’ | 81,1 |
| EMB262 | 12 | 2 | MUC | 29 | EMB262_12_2_MUC_29 | 29 | 1870 | 59°34.443’ | 023°36.461’ | 81,1 |
| EMB262 | 12 | 2 | MUC | 30 | EMB262_12_2_MUC_30 | 30 | 1864 | 59°34.443’ | 023°36.461’ | 81,1 |
| EMB262 | 12 | 2 | MUC | 31 | EMB262_12_2_MUC_31 | 31 | 1859 | 59°34.443’ | 023°36.461’ | 81,1 |
| EMB262 | 12 | 2 | MUC | 32 | EMB262_12_2_MUC_32 | 32 | 1853 | 59°34.443’ | 023°36.461’ | 81,1 |
| EMB262 | 12 | 2 | MUC | 33 | EMB262_12_2_MUC_33 | 33 | 1847 | 59°34.443’ | 023°36.461’ | 81,1 |
| EMB262 | 12 | 2 | MUC | 34 | EMB262_12_2_MUC_34 | 34 | 1841 | 59°34.443’ | 023°36.461’ | 81,1 |
| EMB262 | 12 | 2 | MUC | 35 | EMB262_12_2_MUC_35 | 35 | 1836 | 59°34.443’ | 023°36.461’ | 81,1 |
| EMB262 | 13 | 8 | MUC | 27 | EMB262_13_8_MUC_27 | 2 | 2018 | 58°38.391’ | 018°15.997’ | 436 |
| EMB262 | 13 | 8 | MUC | 26 | EMB262_13_8_MUC_26 | 4 | 2015 | 58°38.391’ | 018°15.997’ | 436 |
| EMB262 | 13 | 8 | MUC | 25 | EMB262_13_8_MUC_25 | 6 | 2012 | 58°38.391’ | 018°15.997’ | 436 |
| EMB262 | 13 | 8 | MUC | 24 | EMB262_13_8_MUC_24 | 8 | 2009 | 58°38.391’ | 018°15.997’ | 436 |
| EMB262 | 13 | 8 | MUC | 23 | EMB262_13_8_MUC_23 | 10 | 2005 | 58°38.391’ | 018°15.997’ | 436 |
| EMB262 | 13 | 8 | MUC | 22 | EMB262_13_8_MUC_22 | 12 | 2002 | 58°38.391’ | 018°15.997’ | 436 |
| EMB262 | 13 | 8 | MUC | 21 | EMB262_13_8_MUC_21 | 14 | 1999 | 58°38.391’ | 018°15.997’ | 436 |
| EMB262 | 13 | 8 | MUC | 20 | EMB262_13_8_MUC_20 | 16 | 1996 | 58°38.391’ | 018°15.997’ | 436 |
| EMB262 | 13 | 8 | MUC | 19 | EMB262_13_8_MUC_19 | 18 | 1994 | 58°38.391’ | 018°15.997’ | 436 |
| EMB262 | 13 | 8 | MUC | 18 | EMB262_13_8_MUC_18 | 20 | 1991 | 58°38.391’ | 018°15.997’ | 436 |
| EMB262 | 13 | 8 | MUC | 17 | EMB262_13_8_MUC_17 | 22 | 1987 | 58°38.391’ | 018°15.997’ | 436 |
| EMB262 | 13 | 8 | MUC | 16 | EMB262_13_8_MUC_16 | 24 | 1985 | 58°38.391’ | 018°15.997’ | 436 |
| EMB262 | 13 | 8 | MUC | 15 | EMB262_13_8_MUC_15 | 26 | 1984 | 58°38.391’ | 018°15.997’ | 436 |
| EMB262 | 13 | 8 | MUC | 14 | EMB262_13_8_MUC_14 | 28 | 1982 | 58°38.391’ | 018°15.997’ | 436 |
| EMB262 | 13 | 8 | MUC | 13 | EMB262_13_8_MUC_13 | 30 | 1980 | 58°38.391’ | 018°15.997’ | 436 |

| | | | | | | | | | | |
|---|---|---|---|---|---|---|---|---|---|---|
| EMB262 | 13 | 8 | MUC | 12 | EMB262_13_8_MUC_12 | 32 | 1978 | 58°38.391’ | 018°15.997’ | 436 |
| EMB262 | 13 | 8 | MUC | 11 | EMB262_13_8_MUC_11 | 34 | 1973 | 58°38.391’ | 018°15.997’ | 436 |
| EMB262 | 13 | 8 | MUC | 10 | EMB262_13_8_MUC_10 | 36 | 1970 | 58°38.391’ | 018°15.997’ | 436 |
| EMB262 | 13 | 8 | MUC | 9 | EMB262_13_8_MUC_9 | 38 | 1965 | 58°38.391’ | 018°15.997’ | 436 |
| EMB262 | 13 | 8 | MUC | 8 | EMB262_13_8_MUC_8 | 40 | 1961 | 58°38.391’ | 018°15.997’ | 436 |
| EMB262 | 13 | 8 | MUC | 7 | EMB262_13_8_MUC_7 | 42 | 1955 | 58°38.391’ | 018°15.997’ | 436 |
| EMB262 | 13 | 8 | MUC | 6 | EMB262_13_8_MUC_6 | 44 | 1950 | 58°38.391’ | 018°15.997’ | 436 |
| EMB262 | 13 | 8 | MUC | 5 | EMB262_13_8_MUC_5 | 46 | 1947 | 58°38.391’ | 018°15.997’ | 436 |
| EMB262 | 13 | 8 | MUC | 4 | EMB262_13_8_MUC_4 | 48 | 1944 | 58°38.391’ | 018°15.997’ | 436 |
| EMB262 | 13 | 8 | MUC | 3 | EMB262_13_8_MUC_3 | 50 | 1940 | 58°38.391’ | 018°15.997’ | 436 |
| EMB262 | 13 | 8 | MUC | 2 | EMB262_13_8_MUC_2 | 52 | 1936 | 58°38.391’ | 018°15.997’ | 436 |
| EMB262 | 13 | 8 | MUC | 1 | EMB262_13_8_MUC_1 | 54 | 1933 | 58°38.391’ | 018°15.997’ | 436 |

A detailed table (Table S3) that provides comprehensive information about the gravity corer samples collected. Each sample is uniquely identified and includes data such as the trip identifier, station number, event number, gear type, sample number, tag, depth, age, latitude, longitude, and water depth at the collection site.

**Table S3: Detailed information of the gravity corer (GC) samples collected during expedition EMB262 (Kremp et al., 2021) in April 2021 onboard the research vessel Elisabeth Mann Borgese.**

| trip | station | event | gear | sample | tag | depth | age | latitude [°N] | longitude [°E] | water depth [m bsl] |
|---|---|---|---|---|---|---|---|---|---|---|
| EMB262 | 6 | 30 | GC | 16 | EMB262_6_30_GC_16 | 32 | 1982 | 57°17.022' | 020°07.285' | 241 |
| EMB262 | 6 | 30 | GC | 15 | EMB262_6_30_GC_15 | 37 | 1955 | 57°17.022' | 020°07.285' | 241 |
| EMB262 | 6 | 30 | GC | 14 | EMB262_6_30_GC_14 | 42 | 1854 | 57°17.022' | 020°07.285' | 241 |
| EMB262 | 6 | 30 | GC | 13 | EMB262_6_30_GC_13 | 47 | 1735 | 57°17.022' | 020°07.285' | 241 |
| EMB262 | 6 | 30 | GC | 12 | EMB262_6_30_GC_12 | 52 | 1615 | 57°17.022' | 020°07.285' | 241 |
| EMB262 | 6 | 30 | GC | 35 | EMB262_6_30_GC_35 | 57 | 1496 | 57°17.022' | 020°07.285' | 241 |
| EMB262 | 6 | 30 | GC | 34 | EMB262_6_30_GC_34 | 62 | 1374 | 57°17.022' | 020°07.285' | 241 |
| EMB262 | 6 | 30 | GC | 33 | EMB262_6_30_GC_33 | 67 | 1297 | 57°17.022' | 020°07.285' | 241 |
| EMB262 | 6 | 30 | GC | 32 | EMB262_6_30_GC_32 | 72 | 1208 | 57°17.022' | 020°07.285' | 241 |
| EMB262 | 6 | 30 | GC | 31 | EMB262_6_30_GC_31 | 77 | 1138 | 57°17.022' | 020°07.285' | 241 |
| EMB262 | 6 | 30 | GC | 30 | EMB262_6_30_GC_30 | 82 | 1075 | 57°17.022' | 020°07.285' | 241 |
| EMB262 | 6 | 30 | GC | 29 | EMB262_6_30_GC_29 | 87 | 1020 | 57°17.022' | 020°07.285' | 241 |
| EMB262 | 6 | 30 | GC | 28 | EMB262_6_30_GC_28 | 92 | 973 | 57°17.022' | 020°07.285' | 241 |
| EMB262 | 6 | 30 | GC | 27 | EMB262_6_30_GC_27 | 97 | 945 | 57°17.022' | 020°07.285' | 241 |
| EMB262 | 6 | 30 | GC | 26 | EMB262_6_30_GC_26 | 102 | 916 | 57°17.022' | 020°07.285' | 241 |
| EMB262 | 6 | 30 | GC | 25 | EMB262_6_30_GC_25 | 107 | 886 | 57°17.022' | 020°07.285' | 241 |
| EMB262 | 6 | 30 | GC | 24 | EMB262_6_30_GC_24 | 112 | 822 | 57°17.022' | 020°07.285' | 241 |
| EMB262 | 6 | 30 | GC | 23 | EMB262_6_30_GC_23 | 117 | 758 | 57°17.022' | 020°07.285' | 241 |
| EMB262 | 6 | 30 | GC | 22 | EMB262_6_30_GC_22 | 122 | 693 | 57°17.022' | 020°07.285' | 241 |
| EMB262 | 6 | 30 | GC | 21 | EMB262_6_30_GC_21 | 127 | 620 | 57°17.022' | 020°07.285' | 241 |
| EMB262 | 6 | 30 | GC | 20 | EMB262_6_30_GC_20 | 132 | 555 | 57°17.022' | 020°07.285' | 241 |
| EMB262 | 6 | 30 | GC | 19 | EMB262_6_30_GC_19 | 137 | 501 | 57°17.022' | 020°07.285' | 241 |
| EMB262 | 6 | 30 | GC | 18 | EMB262_6_30_GC_18 | 142 | 412 | 57°17.022' | 020°07.285' | 241 |
| EMB262 | 6 | 30 | GC | 17 | EMB262_6_30_GC_17 | 146 | 309 | 57°17.022' | 020°07.285' | 241 |
| EMB262 | 6 | 30 | GC | 45 | EMB262_6_30_GC_45 | 152 | 197 | 57°17.022' | 020°07.285' | 241 |
| EMB262 | 6 | 30 | GC | 44 | EMB262_6_30_GC_44 | 162 | 23 | 57°17.022' | 020°07.285' | 241 |
| EMB262 | 6 | 30 | GC | 43 | EMB262_6_30_GC_43 | 172 | -152 | 57°17.022' | 020°07.285' | 241 |
| EMB262 | 6 | 30 | GC | 42 | EMB262_6_30_GC_42 | 182 | -326 | 57°17.022' | 020°07.285' | 241 |
| EMB262 | 6 | 30 | GC | 41 | EMB262_6_30_GC_41 | 192 | -501 | 57°17.022' | 020°07.285' | 241 |
| EMB262 | 6 | 30 | GC | 40 | EMB262_6_30_GC_40 | 202 | -685 | 57°17.022' | 020°07.285' | 241 |
| EMB262 | 6 | 30 | GC | 39 | EMB262_6_30_GC_39 | 212 | -921 | 57°17.022' | 020°07.285' | 241 |
| EMB262 | 6 | 30 | GC | 38 | EMB262_6_30_GC_38 | 222 | -1157 | 57°17.022' | 020°07.285' | 241 |
| EMB262 | 6 | 30 | GC | 37 | EMB262_6_30_GC_37 | 232 | -1393 | 57°17.022' | 020°07.285' | 241 |
| EMB262 | 6 | 30 | GC | 36 | EMB262_6_30_GC_36 | 242 | -1626 | 57°17.022' | 020°07.285' | 241 |
| EMB262 | 6 | 30 | GC | 55 | EMB262_6_30_GC_55 | 252 | -1811 | 57°17.022' | 020°07.285' | 241 |
| EMB262 | 6 | 30 | GC | 54 | EMB262_6_30_GC_54 | 262 | -1960 | 57°17.022' | 020°07.285' | 241 |
| EMB262 | 6 | 30 | GC | 53 | EMB262_6_30_GC_53 | 272 | -2161 | 57°17.022' | 020°07.285' | 241 |
| EMB262 | 6 | 30 | GC | 52 | EMB262_6_30_GC_52 | 282 | -2373 | 57°17.022' | 020°07.285' | 241 |
| EMB262 | 6 | 30 | GC | 51 | EMB262_6_30_GC_51 | 292 | -2587 | 57°17.022' | 020°07.285' | 241 |
| EMB262 | 6 | 30 | GC | 50 | EMB262_6_30_GC_50 | 302 | -2838 | 57°17.022' | 020°07.285' | 241 |
| EMB262 | 6 | 30 | GC | 49 | EMB262_6_30_GC_49 | 312 | -3145 | 57°17.022' | 020°07.285' | 241 |
| EMB262 | 6 | 30 | GC | 48 | EMB262_6_30_GC_48 | 322 | -3471 | 57°17.022' | 020°07.285' | 241 |

| | | | | | | | | | | |
|---|---|---|---|---|---|---|---|---|---|---|
| EMB262 | 6 | 30 | GC | 47 | EMB262_6_30_GC_47 | 332 | -3709 | 57°17.022' | 020°07.285' | 241 |
| EMB262 | 6 | 30 | GC | 46 | EMB262_6_30_GC_46 | 342 | -3918 | 57°17.022' | 020°07.285' | 241 |
| EMB262 | 6 | 30 | GC | 65 | EMB262_6_30_GC_65 | 352 | -4142 | 57°17.022' | 020°07.285' | 241 |
| EMB262 | 6 | 30 | GC | 64 | EMB262_6_30_GC_64 | 362 | -4372 | 57°17.022' | 020°07.285' | 241 |
| EMB262 | 6 | 30 | GC | 63 | EMB262_6_30_GC_63 | 372 | -4584 | 57°17.022' | 020°07.285' | 241 |
| EMB262 | 6 | 30 | GC | 62 | EMB262_6_30_GC_62 | 382 | -4851 | 57°17.022' | 020°07.285' | 241 |
| EMB263 | 6 | 30 | GC | 66 | EMB263_6_30_GC_66 | 387 | -5019 | 57°17.022' | 020°07.285' | 241 |
| EMB262 | 6 | 30 | GC | 61 | EMB262_6_30_GC_61 | 392 | -5188 | 57°17.022' | 020°07.285' | 241 |
| EMB262 | 6 | 30 | GC | 60 | EMB262_6_30_GC_60 | 402 | Ancylus lake | 57°17.022' | 020°07.285' | 241 |
| EMB262 | 6 | 30 | GC | 59 | EMB262_6_30_GC_59 | 412 | Ancylus lake | 57°17.022' | 020°07.285' | 241 |
| EMB262 | 6 | 30 | GC | 58 | EMB262_6_30_GC_58 | 422 | Ancylus lake | 57°17.022' | 020°07.285' | 241 |
| EMB262 | 6 | 30 | GC | 57 | EMB262_6_30_GC_57 | 432 | Ancylus lake | 57°17.022' | 020°07.285' | 241 |
| EMB262 | 6 | 30 | GC | 56 | EMB262_6_30_GC_56 | 442 | Ancylus lake | 57°17.022' | 020°07.285' | 241 |
| EMB262 | 6 | 30 | GC | 10 | EMB262_6_30_GC_10 | 452 | Ancylus lake | 57°17.022' | 020°07.285' | 241 |
| EMB262 | 6 | 30 | GC | 9 | EMB262_6_30_GC_9 | 462 | Ancylus lake | 57°17.022' | 020°07.285' | 241 |
| EMB262 | 6 | 30 | GC | 8 | EMB262_6_30_GC_8 | 472 | Ancylus lake | 57°17.022' | 020°07.285' | 241 |
| EMB262 | 6 | 30 | GC | 7 | EMB262_6_30_GC_7 | 482 | Ancylus lake | 57°17.022' | 020°07.285' | 241 |
| EMB262 | 6 | 30 | GC | 6 | EMB262_6_30_GC_6 | 492 | Ancylus lake | 57°17.022' | 020°07.285' | 241 |
| EMB262 | 6 | 30 | GC | 5 | EMB262_6_30_GC_5 | 502 | Ancylus lake | 57°17.022' | 020°07.285' | 241 |
| EMB262 | 6 | 30 | GC | 4 | EMB262_6_30_GC_4 | 512 | Ancylus lake | 57°17.022' | 020°07.285' | 241 |
| EMB262 | 6 | 30 | GC | 3 | EMB262_6_30_GC_3 | 522 | Ancylus lake | 57°17.022' | 020°07.285' | 241 |
| EMB262 | 6 | 30 | GC | 2 | EMB262_6_30_GC_2 | 532 | Ancylus lake | 57°17.022' | 020°07.285' | 241 |
| EMB262 | 6 | 30 | GC | 1 | EMB262_6_30_GC_1 | 542 | Ancylus lake | 57°17.022' | 020°07.285' | 241 |
| EMB262 | 12 | 3 | GC | 41 | EMB262_12_3_GC_41 | 2 | 2002 | 59°34.450’ | 023°36.455’ | 81,1 |
| EMB262 | 12 | 3 | GC | 40 | EMB262_12_3_GC_40 | 4 | 1984 | 59°34.450’ | 023°36.455’ | 81,1 |
| EMB262 | 12 | 3 | GC | 39 | EMB262_12_3_GC_39 | 6 | 1966 | 59°34.450’ | 023°36.455’ | 81,1 |
| EMB262 | 12 | 3 | GC | 38 | EMB262_12_3_GC_38 | 8 | 1948 | 59°34.450’ | 023°36.455’ | 81,1 |
| EMB262 | 12 | 3 | GC | 37 | EMB262_12_3_GC_37 | 10 | 1930 | 59°34.450’ | 023°36.455’ | 81,1 |
| EMB262 | 12 | 3 | GC | 36 | EMB262_12_3_GC_36 | 12 | 1911 | 59°34.450’ | 023°36.455’ | 81,1 |
| EMB262 | 12 | 3 | GC | 35 | EMB262_12_3_GC_35 | 14 | 1893 | 59°34.450’ | 023°36.455’ | 81,1 |
| EMB262 | 12 | 3 | GC | 34 | EMB262_12_3_GC_34 | 16 | 1875 | 59°34.450’ | 023°36.455’ | 81,1 |
| EMB262 | 12 | 3 | GC | 33 | EMB262_12_3_GC_33 | 18 | 1856 | 59°34.450’ | 023°36.455’ | 81,1 |
| EMB262 | 12 | 3 | GC | 32 | EMB262_12_3_GC_32 | 20 | 1838 | 59°34.450’ | 023°36.455’ | 81,1 |
| EMB262 | 12 | 3 | GC | 31 | EMB262_12_3_GC_31 | 22 | 1818 | 59°34.450’ | 023°36.455’ | 81,1 |
| EMB262 | 12 | 3 | GC | 20 | EMB262_12_3_GC_20 | 28 | 1761 | 59°34.450’ | 023°36.455’ | 81,1 |
| EMB262 | 12 | 3 | GC | 19 | EMB262_12_3_GC_19 | 38 | 1670 | 59°34.450’ | 023°36.455’ | 81,1 |
| EMB262 | 12 | 3 | GC | 18 | EMB262_12_3_GC_18 | 48 | 1576 | 59°34.450’ | 023°36.455’ | 81,1 |
| EMB262 | 12 | 3 | GC | 17 | EMB262_12_3_GC_17 | 58 | 1482 | 59°34.450’ | 023°36.455’ | 81,1 |
| EMB262 | 12 | 3 | GC | 16 | EMB262_12_3_GC_16 | 68 | 1389 | 59°34.450’ | 023°36.455’ | 81,1 |
| EMB262 | 12 | 3 | GC | 15 | EMB262_12_3_GC_15 | 78 | 1296 | 59°34.450’ | 023°36.455’ | 81,1 |
| EMB262 | 12 | 3 | GC | 14 | EMB262_12_3_GC_14 | 88 | 1202 | 59°34.450’ | 023°36.455’ | 81,1 |
| EMB262 | 12 | 3 | GC | 13 | EMB262_12_3_GC_13 | 98 | 1110 | 59°34.450’ | 023°36.455’ | 81,1 |

| | | | | | | | | | | |
|---|---|---|---|---|---|---|---|---|---|---|
| EMB262 | 12 | 3 | GC | 12 | EMB262_12_3_GC_12 | 108 | 1017 | 59°34.450' | 023°36.455' | 81,1 |
| EMB262 | 12 | 3 | GC | 11 | EMB262_12_3_GC_11 | 118 | 924 | 59°34.450' | 023°36.455' | 81,1 |
| EMB262 | 12 | 3 | GC | 30 | EMB262_12_3_GC_30 | 128 | 828 | 59°34.450' | 023°36.455' | 81,1 |
| EMB262 | 12 | 3 | GC | 29 | EMB262_12_3_GC_29 | 138 | 737 | 59°34.450' | 023°36.455' | 81,1 |
| EMB262 | 12 | 3 | GC | 28 | EMB262_12_3_GC_28 | 148 | 628 | 59°34.450' | 023°36.455' | 81,1 |
| EMB262 | 12 | 3 | GC | 27 | EMB262_12_3_GC_27 | 158 | 509 | 59°34.450' | 023°36.455' | 81,1 |
| EMB262 | 12 | 3 | GC | 26 | EMB262_12_3_GC_26 | 168 | 391 | 59°34.450' | 023°36.455' | 81,1 |
| EMB262 | 12 | 3 | GC | 25 | EMB262_12_3_GC_25 | 178 | 315 | 59°34.450' | 023°36.455' | 81,1 |
| EMB262 | 12 | 3 | GC | 24 | EMB262_12_3_GC_24 | 188 | 250 | 59°34.450' | 023°36.455' | 81,1 |
| EMB262 | 12 | 3 | GC | 23 | EMB262_12_3_GC_23 | 198 | 182 | 59°34.450' | 023°36.455' | 81,1 |
| EMB262 | 12 | 3 | GC | 22 | EMB262_12_3_GC_22 | 208 | 71 | 59°34.450' | 023°36.455' | 81,1 |
| EMB262 | 12 | 3 | GC | 21 | EMB262_12_3_GC_21 | 218 | -52 | 59°34.450' | 023°36.455' | 81,1 |
| EMB262 | 12 | 3 | GC | 51 | EMB262_12_3_GC_51 | 228 | -202 | 59°34.450' | 023°36.455' | 81,1 |
| EMB262 | 12 | 3 | GC | 50 | EMB262_12_3_GC_50 | 238 | -402 | 59°34.450' | 023°36.455' | 81,1 |
| EMB262 | 12 | 3 | GC | 49 | EMB262_12_3_GC_49 | 248 | -600 | 59°34.450' | 023°36.455' | 81,1 |
| EMB262 | 12 | 3 | GC | 48 | EMB262_12_3_GC_48 | 258 | -796 | 59°34.450' | 023°36.455' | 81,1 |
| EMB262 | 12 | 3 | GC | 47 | EMB262_12_3_GC_47 | 268 | -955 | 59°34.450' | 023°36.455' | 81,1 |
| EMB262 | 12 | 3 | GC | 46 | EMB262_12_3_GC_46 | 278 | -1103 | 59°34.450' | 023°36.455' | 81,1 |
| EMB262 | 12 | 3 | GC | 45 | EMB262_12_3_GC_45 | 288 | -1251 | 59°34.450' | 023°36.455' | 81,1 |
| EMB262 | 12 | 3 | GC | 44 | EMB262_12_3_GC_44 | 298 | -1398 | 59°34.450' | 023°36.455' | 81,1 |
| EMB262 | 12 | 3 | GC | 43 | EMB262_12_3_GC_43 | 308 | -1532 | 59°34.450' | 023°36.455' | 81,1 |
| EMB262 | 12 | 3 | GC | 42 | EMB262_12_3_GC_42 | 318 | -1623 | 59°34.450' | 023°36.455' | 81,1 |
| EMB262 | 12 | 3 | GC | 10 | EMB262_12_3_GC_10 | 328 | -1711 | 59°34.450' | 023°36.455' | 81,1 |
| EMB262 | 12 | 3 | GC | 9 | EMB262_12_3_GC_9 | 338 | -1800 | 59°34.450' | 023°36.455' | 81,1 |
| EMB262 | 12 | 3 | GC | 8 | EMB262_12_3_GC_8 | 348 | -1887 | 59°34.450' | 023°36.455' | 81,1 |
| EMB262 | 12 | 3 | GC | 7 | EMB262_12_3_GC_7 | 358 | -1974 | 59°34.450' | 023°36.455' | 81,1 |
| EMB262 | 12 | 3 | GC | 6 | EMB262_12_3_GC_6 | 368 | -2061 | 59°34.450' | 023°36.455' | 81,1 |
| EMB262 | 12 | 3 | GC | 5 | EMB262_12_3_GC_5 | 378 | -2147 | 59°34.450' | 023°36.455' | 81,1 |
| EMB262 | 12 | 3 | GC | 4 | EMB262_12_3_GC_4 | 388 | -2234 | 59°34.450' | 023°36.455' | 81,1 |
| EMB262 | 12 | 3 | GC | 3 | EMB262_12_3_GC_3 | 398 | -2322 | 59°34.450' | 023°36.455' | 81,1 |
| EMB262 | 12 | 3 | GC | 2 | EMB262_12_3_GC_2 | 408 | -2409 | 59°34.450' | 023°36.455' | 81,1 |
| EMB262 | 12 | 3 | GC | 1 | EMB262_12_3_GC_1 | 418 | -2496 | 59°34.450' | 023°36.455' | 81,1 |

#### Core dating

Figure S1 illustrates the construction of age models for the short cores (MUCs). It includes three graphs (A, B, C) that demonstrate the correlation of Hg and Mn contents with previously dated cores, and an age model based on event stratigraphy. These correlations and models provide more information of the core samples and their respective ages.

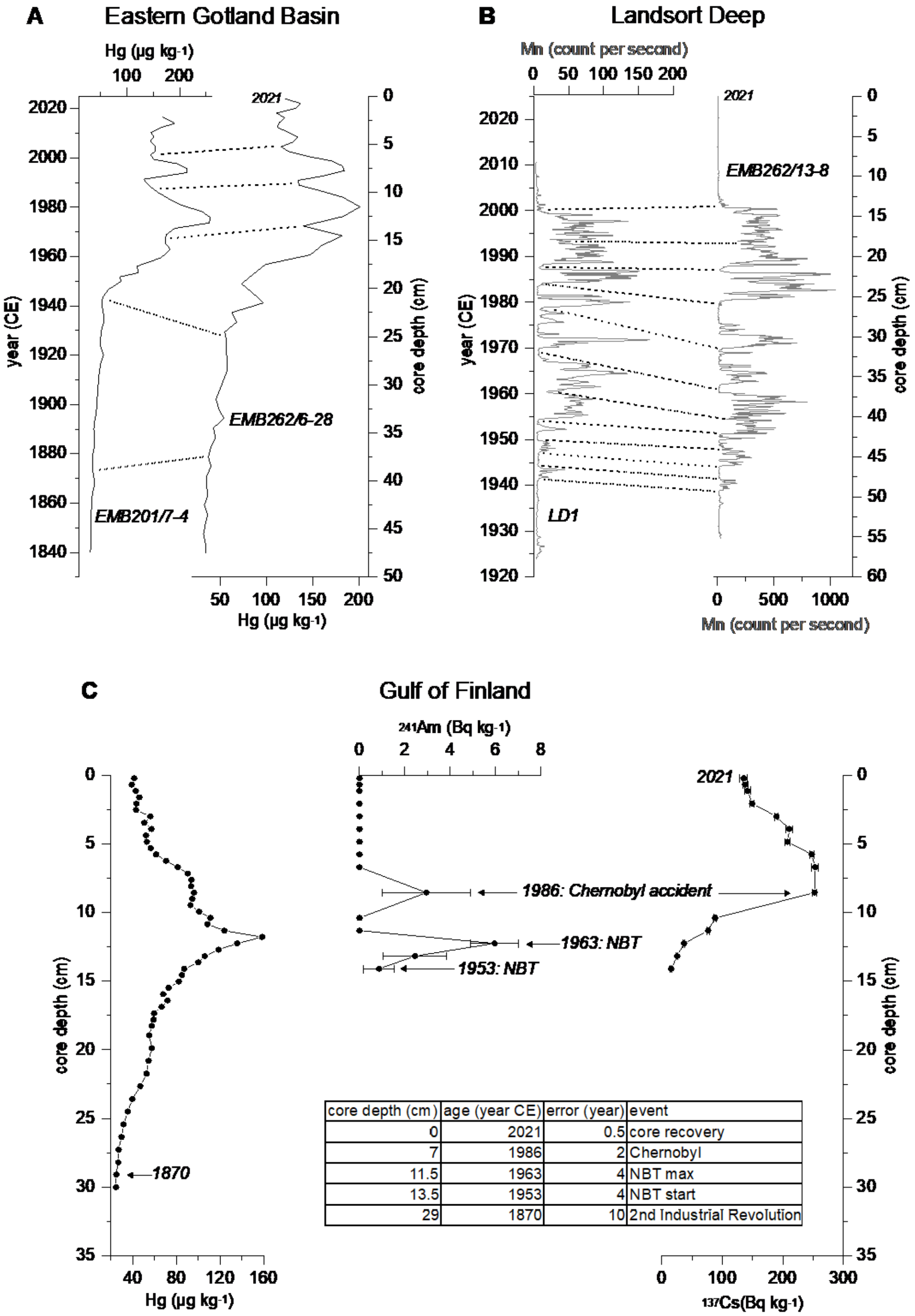

| core depth (cm) | age (year CE) | error (year) | event |
|---|---|---|---|
| 0 | 2021 | 0.5 | core recovery |
| 7 | 1986 | 2 | Chernobyl |
| 11.5 | 1963 | 4 | NBT max |
| 13.5 | 1953 | 4 | NBT start |
| 29 | 1870 | 10 | 2nd Industrial Revolution |

**Fig. S1: Construction of the age models for the short cores. (A) Pointers (dashed lines) used to correlate the Hg contents of core EMB262/6-28 with the previously dated core EMB201/7-4 retrieved at virtually the same location (Kaiser et al., 2022); (B) Pointers (dashed lines) used to correlate the Mn relative contents of core EMB262/13-8 with the previously dated core LD1 retrieved at virtually the same location (Häulser et al., 2018); (C) age model of core EMB262/12-2 based on an event stratigraphy (see main text for details).**

The age assignment of sediment core EMB262-6-30-GC (EGB long core) is depicted in Figure S2. This is achieved through a detailed visual correlation of high-resolution XRF scanner elemental Br/K ratios. These ratios are compared with loss-on-ignition (LOI) and total organic carbon (TOC) data from recently published central Baltic Sea sediment cores owning exceptionally well-dated radiocarbon chronologies. For the older, mid-Holocene part, the chronology of sediment core P435-2-1 cored at about the same location as EMB262-6-30 in the EGB was used as a reference (Warden et al. 2017). For the late Holocene part (last ~3000 yr) sediment cores M86-1a/36 (Häusler et al. 2018) and MSM62-60 (Moros et al. 2020) from the western Gotland Basin served as reference chronologies. A precise age assignment of pre-Littorina-Stage sediments was not possible, but the lowermost about 50 cm of the core most likely belong to the late Ancylus Lake phase < 9500 cal yr BP of the Baltic Sea.

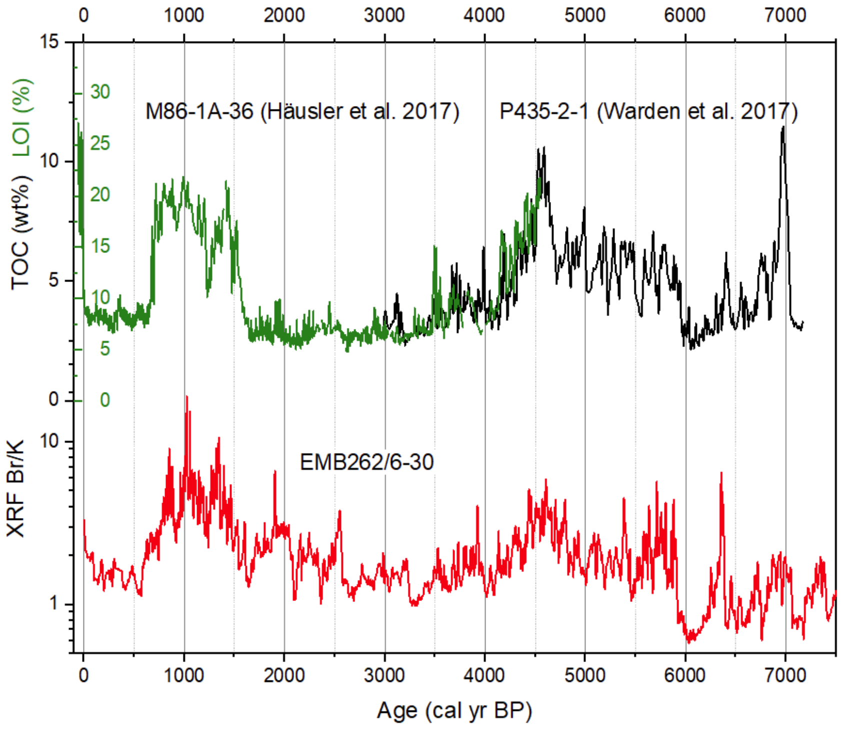

**Figure S2: Age assignment of sediment core EMB262-6-30 through detailed visual correlation of high-resolution XRF scanner elemental Br/K ratios with LOI and total organic carbon data from published central Baltic Sea sediment cores (Häusler et al. 2017; Moros et al. 2020).**

The age-depth model of the EMB262-12-3-GC (GOF long core) sediment core is illustrated in Figure S3. It shows a comprehensive analysis using BACON 2.5.5 software and Marine20 calibration curve (Blaauw and Christen, 2011). These findings were obtained from seven radiocarbon dates derived from the humic acid fraction of bulk organic carbon samples (Table S4).

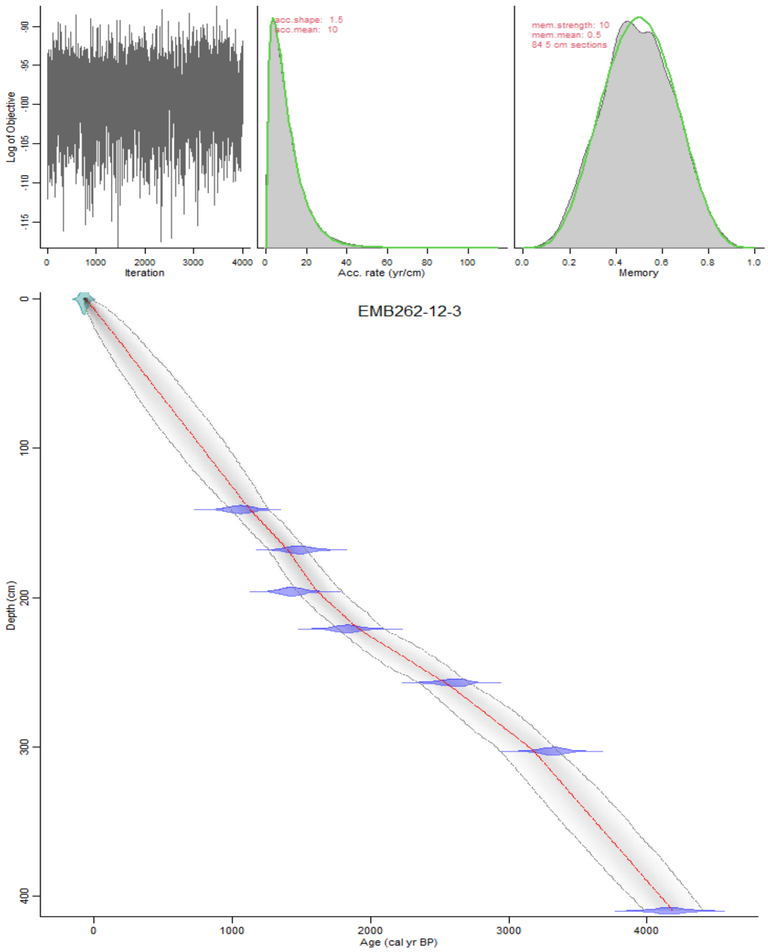

**Figure S3: Age-depth model of the EMB262-12-3 sediment core produced with the BACON 2.5.5 software (Blaauw and Christen, 2011) and Marine20 calibration curve (Heaton et al., 2020), using seven radiocarbon dates obtained from dating of the humic acid fraction of bulk organic carbon samples (Table S4). The upper panels depict the Markov Chain Monte Carlo iterations (left panel), and the prior (green curves) and posterior (gray histograms) distributions of the sedimentation rate (middle panel) and memory (right panel). The lower panel shows the calibrated 14C dates and the age-depth model. The probability distribution of the calibrated 14C ages is indicated in blue. The 95% confidence interval of the age-depth model is indicated by the gray dashed lines. The mean age of the model is shown with the red line.**

**Table S4: Radiocarbon dates obtained from AMS dating of the humic acid fraction of bulk organic carbon samples from gravity core EMB262-12-3.**

| Lab.number | Core depth (cm) | Conventional radiocarbon age (yr BP) |
|---|---|---|
| Beta-625102 | 141 | 1670 ±30 |
| Beta-625103 | 168 | 2090 ±30 |
| Beta-625104 | 196 | 2040 ±30 |
| Beta-625105 | 221 | 2380 ±30 |
| Beta-625106 | 257 | 2990 ±30 |
| Beta-625107 | 303 | 3590 ±30 |
| Beta-625108 | 410 | 4250 ±30 |

### Metabarcoding

Metabarcoding results of the DNA extracts are detailed in Table S5. Each sample is uniquely identified by a tag. The table delineates the taxonomic division and genus of the organisms found in each sample, along with the unique sequence variant identifier (ASV), the total number of sequence reads for the ASV, and the number of replicates in which the ASV was found.

**Table S5: Metabarcoding of Dinoflagellates and Diatoms from DNA extracts of EGB262 samples. Each sample from each location is uniquely tagged (tag). Displayed are the corresponding division, sequence variant identifier (ASV), total reads (sum_reads), and the count of replicates where the ASV was present (repl).**

| tag | division | ASV | sum_reads | repl |
|---|---|---|---|---|
| EMB262_12_2_GC_1 | Dinoflagellates | 0 | 0 | 0 |
| EMB262_12_2_GC_1 | Diatoms | 0 | 0 | 0 |
| EMB262_12_2_GC_10 | Dinoflagellates | 1 | 1 | 1 |
| EMB262_12_2_GC_10 | Diatoms | 1 | 125 | 1 |
| EMB262_12_2_GC_11 | Dinoflagellates | 1 | 2325 | 4 |
| EMB262_12_2_GC_11 | Diatoms | 1 | 1 | 2 |
| EMB262_12_2_GC_12 | Dinoflagellates | 1 | 16 | 4 |
| EMB262_12_2_GC_12 | Diatoms | 0 | 0 | 0 |
| EMB262_12_2_GC_13 | Dinoflagellates | 1 | 5 | 1 |
| EMB262_12_2_GC_13 | Diatoms | 2 | 225 | 2 |
| EMB262_12_2_GC_14 | Dinoflagellates | 0 | 0 | 0 |
| EMB262_12_2_GC_14 | Diatoms | 1 | 5 | 1 |
| EMB262_12_2_GC_15 | Dinoflagellates | 1 | 51 | 4 |
| EMB262_12_2_GC_15 | Diatoms | 1 | 1 | 3 |
| EMB262_12_2_GC_16 | Dinoflagellates | 1 | 1 | 2 |
| EMB262_12_2_GC_16 | Diatoms | 0 | 0 | 0 |
| EMB262_12_2_GC_17 | Dinoflagellates | 1 | 5575 | 4 |
| EMB262_12_2_GC_17 | Diatoms | 1 | 75 | 1 |
| EMB262_12_2_GC_18 | Dinoflagellates | 1 | 875 | 4 |
| EMB262_12_2_GC_18 | Diatoms | 1 | 2 | 2 |
| EMB262_12_2_GC_19 | Dinoflagellates | 1 | 25 | 1 |
| EMB262_12_2_GC_19 | Diatoms | 1 | 425 | 3 |
| EMB262_12_2_GC_2 | Dinoflagellates | 1 | 26225 | 2 |
| EMB262_12_2_GC_2 | Diatoms | 2 | 225 | 3 |
| EMB262_12_2_GC_20 | Dinoflagellates | 1 | 175 | 1 |
| EMB262_12_2_GC_20 | Diatoms | 0 | 0 | 0 |
| EMB262_12_2_GC_21 | Dinoflagellates | 1 | 15 | 1 |
| EMB262_12_2_GC_21 | Diatoms | 1 | 125 | 1 |
| EMB262_12_2_GC_22 | Dinoflagellates | 0 | 0 | 0 |
| EMB262_12_2_GC_22 | Diatoms | 0 | 0 | 0 |
| EMB262_12_2_GC_23 | Dinoflagellates | 0 | 0 | 0 |
| EMB262_12_2_GC_23 | Diatoms | 1 | 65 | 3 |
| EMB262_12_2_GC_24 | Dinoflagellates | 0 | 0 | 0 |
| EMB262_12_2_GC_24 | Diatoms | 1 | 25 | 1 |
| EMB262_12_2_GC_25 | Dinoflagellates | 0 | 0 | 0 |
| EMB262_12_2_GC_25 | Diatoms | 0 | 0 | 0 |

| | | | | |
|---|---|---|---|---|
| EMB262_12_2_GC_26 | Dinoflagellates | 0 | 0 | 0 |
| EMB262_12_2_GC_26 | Diatoms | 0 | 0 | 0 |
| EMB262_12_2_GC_28 | Dinoflagellates | 0 | 0 | 0 |
| EMB262_12_2_GC_28 | Diatoms | 1 | 125 | 1 |
| EMB262_12_2_GC_29 | Dinoflagellates | 1 | 2325 | 4 |
| EMB262_12_2_GC_29 | Diatoms | 0 | 0 | 0 |
| EMB262_12_2_GC_3 | Dinoflagellates | 0 | 0 | 0 |
| EMB262_12_2_GC_3 | Diatoms | 1 | 225 | 2 |
| EMB262_12_2_GC_30 | Dinoflagellates | 0 | 0 | 0 |
| EMB262_12_2_GC_30 | Diatoms | 0 | 0 | 0 |
| EMB262_12_2_GC_30 | Dinoflagellates | 0 | 0 | 0 |
| EMB262_12_2_GC_30 | Diatoms | 2 | 25 | 3 |
| EMB262_12_2_GC_31 | Dinoflagellates | 1 | 45 | 2 |
| EMB262_12_2_GC_31 | Diatoms | 1 | 26 | 2 |
| EMB262_12_2_GC_32 | Dinoflagellates | 0 | 0 | 0 |
| EMB262_12_2_GC_32 | Diatoms | 2 | 525 | 2 |
| EMB262_12_2_GC_33 | Dinoflagellates | 0 | 0 | 0 |
| EMB262_12_2_GC_33 | Diatoms | 2 | 25 | 3 |
| EMB262_12_2_GC_35 | Dinoflagellates | 0 | 0 | 0 |
| EMB262_12_2_GC_35 | Diatoms | 2 | 425 | 2 |
| EMB262_12_2_GC_36 | Dinoflagellates | 1 | 325 | 1 |
| EMB262_12_2_GC_36 | Diatoms | 1 | 225 | 1 |
| EMB262_12_2_GC_37 | Dinoflagellates | 1 | 6 | 3 |
| EMB262_12_2_GC_37 | Diatoms | 0 | 0 | 0 |
| EMB262_12_2_GC_38 | Dinoflagellates | 0 | 0 | 0 |
| EMB262_12_2_GC_38 | Diatoms | 1 | 25 | 1 |
| EMB262_12_2_GC_39 | Dinoflagellates | 0 | 0 | 0 |
| EMB262_12_2_GC_39 | Diatoms | 1 | 175 | 1 |
| EMB262_12_2_GC_4 | Dinoflagellates | 1 | 218 | 3 |
| EMB262_12_2_GC_4 | Diatoms | 1 | 6 | 2 |
| EMB262_12_2_GC_40 | Dinoflagellates | 0 | 0 | 0 |
| EMB262_12_2_GC_40 | Diatoms | 1 | 1 | 1 |
| EMB262_12_2_GC_42 | Dinoflagellates | 1 | 5 | 1 |
| EMB262_12_2_GC_42 | Diatoms | 1 | 25 | 1 |
| EMB262_12_2_GC_43 | Dinoflagellates | 0 | 0 | 0 |
| EMB262_12_2_GC_43 | Diatoms | 2 | 45 | 2 |
| EMB262_12_2_GC_44 | Dinoflagellates | 0 | 0 | 0 |
| EMB262_12_2_GC_44 | Diatoms | 1 | 4 | 1 |
| EMB262_12_2_GC_45 | Dinoflagellates | 1 | 175 | 1 |
| EMB262_12_2_GC_45 | Diatoms | 1 | 1 | 2 |
| EMB262_12_2_GC_46 | Dinoflagellates | 0 | 0 | 0 |
| EMB262_12_2_GC_46 | Diatoms | 1 | 525 | 2 |
| EMB262_12_2_GC_47 | Dinoflagellates | 1 | 25 | 1 |
| EMB262_12_2_GC_47 | Diatoms | 1 | 25 | 1 |
| EMB262_12_2_GC_48 | Dinoflagellates | 1 | 2 | 2 |
| EMB262_12_2_GC_48 | Diatoms | 1 | 5 | 1 |
| EMB262_12_2_GC_49 | Dinoflagellates | 0 | 0 | 0 |

| | | | | |
|---|---|---|---|---|
| EMB262_12_2_GC_49 | Diatoms | 1 | 1 | 1 |
| EMB262_12_2_GC_5 | Dinoflagellates | 1 | 28825 | 4 |
| EMB262_12_2_GC_5 | Diatoms | 1 | 25 | 1 |
| EMB262_12_2_GC_50 | Dinoflagellates | 0 | 0 | 0 |
| EMB262_12_2_GC_50 | Diatoms | 1 | 25 | 1 |
| EMB262_12_2_GC_51 | Dinoflagellates | 0 | 0 | 0 |
| EMB262_12_2_GC_51 | Diatoms | 2 | 225 | 2 |
| EMB262_12_2_GC_6 | Dinoflagellates | 0 | 0 | 0 |
| EMB262_12_2_GC_6 | Diatoms | 0 | 0 | 0 |
| EMB262_12_2_GC_7 | Dinoflagellates | 0 | 0 | 0 |
| EMB262_12_2_GC_7 | Diatoms | 2 | 5 | 2 |
| EMB262_12_2_GC_8 | Dinoflagellates | 1 | 35 | 1 |
| EMB262_12_2_GC_8 | Diatoms | 3 | 1425 | 5 |
| EMB262_12_2_GC_9 | Dinoflagellates | 1 | 125 | 1 |
| EMB262_12_2_GC_9 | Diatoms | 2 | 105 | 4 |
| EMB262_12_2_MUC_1 | Dinoflagellates | 1 | 825 | 4 |
| EMB262_12_2_MUC_1 | Diatoms | 0 | 0 | 0 |
| EMB262_12_2_MUC_10 | Dinoflagellates | 1 | 1095 | 4 |
| EMB262_12_2_MUC_10 | Diatoms | 0 | 0 | 0 |
| EMB262_12_2_MUC_11 | Dinoflagellates | 1 | 3525 | 4 |
| EMB262_12_2_MUC_11 | Diatoms | 0 | 0 | 0 |
| EMB262_12_2_MUC_12 | Dinoflagellates | 0 | 0 | 0 |
| EMB262_12_2_MUC_12 | Diatoms | 0 | 0 | 0 |
| EMB262_12_2_MUC_13 | Dinoflagellates | 0 | 0 | 0 |
| EMB262_12_2_MUC_13 | Diatoms | 0 | 0 | 0 |
| EMB262_12_2_MUC_14 | Dinoflagellates | 1 | 10225 | 4 |
| EMB262_12_2_MUC_14 | Diatoms | 0 | 0 | 0 |
| EMB262_12_2_MUC_15 | Dinoflagellates | 0 | 0 | 0 |
| EMB262_12_2_MUC_15 | Diatoms | 1 | 75 | 2 |
| EMB262_12_2_MUC_16 | Dinoflagellates | 1 | 475 | 2 |
| EMB262_12_2_MUC_16 | Diatoms | 1 | 125 | 1 |
| EMB262_12_2_MUC_17 | Dinoflagellates | 1 | 43 | 4 |
| EMB262_12_2_MUC_17 | Diatoms | 1 | 25 | 1 |
| EMB262_12_2_MUC_18 | Dinoflagellates | 0 | 0 | 0 |
| EMB262_12_2_MUC_18 | Diatoms | 1 | 125 | 2 |
| EMB262_12_2_MUC_19 | Dinoflagellates | 0 | 0 | 0 |
| EMB262_12_2_MUC_19 | Diatoms | 1 | 25 | 1 |
| EMB262_12_2_MUC_2 | Dinoflagellates | 1 | 185 | 3 |
| EMB262_12_2_MUC_2 | Diatoms | 0 | 0 | 0 |
| EMB262_12_2_MUC_20 | Dinoflagellates | 0 | 0 | 0 |
| EMB262_12_2_MUC_20 | Diatoms | 1 | 1 | 1 |
| EMB262_12_2_MUC_21 | Dinoflagellates | 0 | 0 | 0 |
| EMB262_12_2_MUC_21 | Diatoms | 2 | 1225 | 2 |
| EMB262_12_2_MUC_22 | Dinoflagellates | 1 | 25 | 2 |
| EMB262_12_2_MUC_22 | Diatoms | 0 | 0 | 0 |
| EMB262_12_2_MUC_23 | Dinoflagellates | 0 | 0 | 0 |
| EMB262_12_2_MUC_23 | Diatoms | 1 | 375 | 1 |

| | | | | |
|---|---|---|---|---|
| EMB262_12_2_MUC_24 | Dinoflagellates | 0 | 0 | 0 |
| EMB262_12_2_MUC_24 | Diatoms | 0 | 0 | 0 |
| EMB262_12_2_MUC_25 | Dinoflagellates | 0 | 0 | 0 |
| EMB262_12_2_MUC_25 | Diatoms | 1 | 3 | 3 |
| EMB262_12_2_MUC_26 | Dinoflagellates | 0 | 0 | 0 |
| EMB262_12_2_MUC_26 | Diatoms | 0 | 0 | 0 |
| EMB262_12_2_MUC_27 | Dinoflagellates | 1 | 4 | 1 |
| EMB262_12_2_MUC_27 | Diatoms | 2 | 75 | 2 |
| EMB262_12_2_MUC_28 | Dinoflagellates | 1 | 575 | 2 |
| EMB262_12_2_MUC_28 | Diatoms | 2 | 1225 | 4 |
| EMB262_12_2_MUC_29 | Dinoflagellates | 0 | 0 | 0 |
| EMB262_12_2_MUC_29 | Diatoms | 0 | 0 | 0 |
| EMB262_12_2_MUC_3 | Dinoflagellates | 1 | 49425 | 4 |
| EMB262_12_2_MUC_3 | Diatoms | 0 | 0 | 0 |
| EMB262_12_2_MUC_30 | Dinoflagellates | 0 | 0 | 0 |
| EMB262_12_2_MUC_30 | Diatoms | 1 | 1125 | 2 |
| EMB262_12_2_MUC_31 | Dinoflagellates | 0 | 0 | 0 |
| EMB262_12_2_MUC_31 | Diatoms | 1 | 175 | 1 |
| EMB262_12_2_MUC_32 | Dinoflagellates | 0 | 0 | 0 |
| EMB262_12_2_MUC_32 | Diatoms | 3 | 7 | 3 |
| EMB262_12_2_MUC_33 | Dinoflagellates | 0 | 0 | 0 |
| EMB262_12_2_MUC_33 | Diatoms | 0 | 0 | 0 |
| EMB262_12_2_MUC_34 | Dinoflagellates | 0 | 0 | 0 |
| EMB262_12_2_MUC_34 | Diatoms | 2 | 165 | 2 |
| EMB262_12_2_MUC_35 | Dinoflagellates | 1 | 1325 | 1 |
| EMB262_12_2_MUC_35 | Diatoms | 2 | 20 | 2 |
| EMB262_12_2_MUC_4 | Dinoflagellates | 1 | 36025 | 4 |
| EMB262_12_2_MUC_4 | Diatoms | 0 | 0 | 0 |
| EMB262_12_2_MUC_5 | Dinoflagellates | 1 | 32 | 3 |
| EMB262_12_2_MUC_5 | Diatoms | 1 | 25 | 1 |
| EMB262_12_2_MUC_6 | Dinoflagellates | 1 | 465 | 4 |
| EMB262_12_2_MUC_6 | Diatoms | 0 | 0 | 0 |
| EMB262_12_2_MUC_6 | Dinoflagellates | 1 | 605 | 4 |
| EMB262_12_2_MUC_6 | Diatoms | 0 | 0 | 0 |
| EMB262_12_2_MUC_7 | Dinoflagellates | 1 | 362 | 4 |
| EMB262_12_2_MUC_7 | Diatoms | 0 | 0 | 0 |
| EMB262_12_2_MUC_8 | Dinoflagellates | 1 | 12335 | 4 |
| EMB262_12_2_MUC_8 | Diatoms | 0 | 0 | 0 |
| EMB262_12_2_MUC_9 | Dinoflagellates | 1 | 7815 | 4 |
| EMB262_12_2_MUC_9 | Diatoms | 1 | 5 | 1 |
| EMB262_13_8_MUC_1 | Dinoflagellates | 0 | 0 | 0 |
| EMB262_13_8_MUC_1 | Diatoms | 2 | 125 | 4 |
| EMB262_13_8_MUC_10 | Dinoflagellates | 0 | 0 | 0 |
| EMB262_13_8_MUC_10 | Diatoms | 1 | 2465 | 1 |
| EMB262_13_8_MUC_11 | Dinoflagellates | 0 | 0 | 0 |
| EMB262_13_8_MUC_11 | Diatoms | 0 | 0 | 0 |
| EMB262_13_8_MUC_12 | Dinoflagellates | 0 | 0 | 0 |

| | | | | |
|---|---|---|---|---|
| EMB262_13_8_MUC_12 | Diatoms | 0 | 0 | 0 |
| EMB262_13_8_MUC_13 | Dinoflagellates | 0 | 0 | 0 |
| EMB262_13_8_MUC_13 | Diatoms | 0 | 0 | 0 |
| EMB262_13_8_MUC_14 | Dinoflagellates | 0 | 0 | 0 |
| EMB262_13_8_MUC_14 | Diatoms | 0 | 0 | 0 |
| EMB262_13_8_MUC_15 | Dinoflagellates | 0 | 0 | 0 |
| EMB262_13_8_MUC_15 | Diatoms | 0 | 0 | 0 |
| EMB262_13_8_MUC_16 | Dinoflagellates | 0 | 0 | 0 |
| EMB262_13_8_MUC_16 | Diatoms | 0 | 0 | 0 |
| EMB262_13_8_MUC_17 | Dinoflagellates | 1 | 4075 | 4 |
| EMB262_13_8_MUC_17 | Diatoms | 0 | 0 | 0 |
| EMB262_13_8_MUC_18 | Dinoflagellates | 1 | 205 | 4 |
| EMB262_13_8_MUC_18 | Diatoms | 0 | 0 | 0 |
| EMB262_13_8_MUC_19 | Dinoflagellates | 0 | 0 | 0 |
| EMB262_13_8_MUC_19 | Diatoms | 0 | 0 | 0 |
| EMB262_13_8_MUC_2 | Dinoflagellates | 1 | 14325 | 4 |
| EMB262_13_8_MUC_2 | Diatoms | 1 | 25 | 1 |
| EMB262_13_8_MUC_20 | Dinoflagellates | 0 | 0 | 0 |
| EMB262_13_8_MUC_20 | Diatoms | 1 | 5 | 1 |
| EMB262_13_8_MUC_21 | Dinoflagellates | 0 | 0 | 0 |
| EMB262_13_8_MUC_21 | Diatoms | 2 | 5 | 2 |
| EMB262_13_8_MUC_22 | Dinoflagellates | 0 | 0 | 0 |
| EMB262_13_8_MUC_22 | Diatoms | 0 | 0 | 0 |
| EMB262_13_8_MUC_23 | Dinoflagellates | 0 | 0 | 0 |
| EMB262_13_8_MUC_23 | Diatoms | 1 | 75 | 2 |
| EMB262_13_8_MUC_24 | Dinoflagellates | 0 | 0 | 0 |
| EMB262_13_8_MUC_24 | Diatoms | 0 | 0 | 0 |
| EMB262_13_8_MUC_25 | Dinoflagellates | 1 | 675 | 3 |
| EMB262_13_8_MUC_25 | Diatoms | 0 | 0 | 0 |
| EMB262_13_8_MUC_26 | Dinoflagellates | 1 | 825 | 3 |
| EMB262_13_8_MUC_26 | Diatoms | 0 | 0 | 0 |
| EMB262_13_8_MUC_27 | Dinoflagellates | 1 | 1575 | 4 |
| EMB262_13_8_MUC_27 | Diatoms | 0 | 0 | 0 |
| EMB262_13_8_MUC_3 | Dinoflagellates | 1 | 3425 | 4 |
| EMB262_13_8_MUC_3 | Diatoms | 1 | 25 | 1 |
| EMB262_13_8_MUC_4 | Dinoflagellates | 1 | 3825 | 4 |
| EMB262_13_8_MUC_4 | Diatoms | 2 | 75 | 3 |
| EMB262_13_8_MUC_5 | Dinoflagellates | 0 | 0 | 0 |
| EMB262_13_8_MUC_5 | Diatoms | 4 | 385125 | 8 |
| EMB262_13_8_MUC_6 | Dinoflagellates | 1 | 4275 | 4 |
| EMB262_13_8_MUC_6 | Diatoms | 3 | 175 | 5 |
| EMB262_13_8_MUC_7 | Dinoflagellates | 0 | 0 | 0 |
| EMB262_13_8_MUC_7 | Diatoms | 0 | 0 | 0 |
| EMB262_13_8_MUC_8 | Dinoflagellates | 0 | 0 | 0 |
| EMB262_13_8_MUC_8 | Diatoms | 0 | 0 | 0 |
| EMB262_13_8_MUC_9 | Dinoflagellates | 0 | 0 | 0 |
| EMB262_13_8_MUC_9 | Diatoms | 0 | 0 | 0 |

| | | | | |
|---|---|---|---|---|
| EMB262_6_28_MUC_1 | Dinoflagellates | 0 | 0 | 0 |
| EMB262_6_28_MUC_1 | Diatoms | 0 | 0 | 0 |
| EMB262_6_28_MUC_11 | Dinoflagellates | 0 | 0 | 0 |
| EMB262_6_28_MUC_11 | Diatoms | 0 | 0 | 0 |
| EMB262_6_28_MUC_12 | Dinoflagellates | 0 | 0 | 0 |
| EMB262_6_28_MUC_12 | Diatoms | 1 | 1 | 3 |
| EMB262_6_28_MUC_13 | Dinoflagellates | 0 | 0 | 0 |
| EMB262_6_28_MUC_13 | Diatoms | 1 | 5 | 1 |
| EMB262_6_28_MUC_14 | Dinoflagellates | 1 | 25 | 1 |
| EMB262_6_28_MUC_14 | Diatoms | 0 | 0 | 0 |
| EMB262_6_28_MUC_15 | Dinoflagellates | 0 | 0 | 0 |
| EMB262_6_28_MUC_15 | Diatoms | 2 | 1 | 2 |
| EMB262_6_28_MUC_16 | Dinoflagellates | 0 | 0 | 0 |
| EMB262_6_28_MUC_16 | Diatoms | 0 | 0 | 0 |
| EMB262_6_28_MUC_17 | Dinoflagellates | 0 | 0 | 0 |
| EMB262_6_28_MUC_17 | Diatoms | 3 | 125 | 4 |
| EMB262_6_28_MUC_18 | Dinoflagellates | 0 | 0 | 0 |
| EMB262_6_28_MUC_18 | Diatoms | 0 | 0 | 0 |
| EMB262_6_28_MUC_19 | Dinoflagellates | 0 | 0 | 0 |
| EMB262_6_28_MUC_19 | Diatoms | 0 | 0 | 0 |
| EMB262_6_28_MUC_20 | Dinoflagellates | 1 | 145 | 1 |
| EMB262_6_28_MUC_20 | Diatoms | 1 | 125 | 1 |
| EMB262_6_28_MUC_21 | Dinoflagellates | 0 | 0 | 0 |
| EMB262_6_28_MUC_21 | Diatoms | 2 | 75 | 2 |
| EMB262_6_28_MUC_22 | Dinoflagellates | 0 | 0 | 0 |
| EMB262_6_28_MUC_22 | Diatoms | 1 | 5 | 1 |
| EMB262_6_28_MUC_23 | Dinoflagellates | 0 | 0 | 0 |
| EMB262_6_28_MUC_23 | Diatoms | 1 | 75 | 2 |
| EMB262_6_28_MUC_24 | Dinoflagellates | 0 | 0 | 0 |
| EMB262_6_28_MUC_24 | Diatoms | 1 | 575 | 2 |
| EMB262_6_28_MUC_26 | Dinoflagellates | 0 | 0 | 0 |
| EMB262_6_28_MUC_26 | Diatoms | 2 | 575 | 4 |
| EMB262_6_28_MUC_27 | Dinoflagellates | 0 | 0 | 0 |
| EMB262_6_28_MUC_27 | Diatoms | 2 | 1925 | 3 |
| EMB262_6_28_MUC_28 | Dinoflagellates | 0 | 0 | 0 |
| EMB262_6_28_MUC_28 | Diatoms | 1 | 1 | 2 |
| EMB262_6_28_MUC_28 | Dinoflagellates | 0 | 0 | 0 |
| EMB262_6_28_MUC_28 | Diatoms | 1 | 125 | 1 |
| EMB262_6_28_MUC_29 | Dinoflagellates | 0 | 0 | 0 |
| EMB262_6_28_MUC_29 | Diatoms | 1 | 25 | 1 |
| EMB262_6_28_MUC_3 | Dinoflagellates | 1 | 325 | 4 |
| EMB262_6_28_MUC_3 | Diatoms | 0 | 0 | 0 |
| EMB262_6_28_MUC_30 | Dinoflagellates | 1 | 175 | 2 |
| EMB262_6_28_MUC_30 | Diatoms | 1 | 75 | 1 |
| EMB262_6_28_MUC_31 | Dinoflagellates | 0 | 0 | 0 |
| EMB262_6_28_MUC_31 | Diatoms | 0 | 0 | 0 |
| EMB262_6_28_MUC_32 | Dinoflagellates | 0 | 0 | 0 |

| | | | | |
|---|---|---|---|---|
| EMB262_6_28_MUC_32 | Diatoms | 1 | 15 | 1 |
| EMB262_6_28_MUC_33 | Dinoflagellates | 0 | 0 | 0 |
| EMB262_6_28_MUC_33 | Diatoms | 1 | 1 | 1 |
| EMB262_6_28_MUC_34 | Dinoflagellates | 0 | 0 | 0 |
| EMB262_6_28_MUC_34 | Diatoms | 1 | 25 | 1 |
| EMB262_6_28_MUC_35 | Dinoflagellates | 0 | 0 | 0 |
| EMB262_6_28_MUC_35 | Diatoms | 0 | 0 | 0 |
| EMB262_6_28_MUC_36 | Dinoflagellates | 0 | 0 | 0 |
| EMB262_6_28_MUC_36 | Diatoms | 0 | 0 | 0 |
| EMB262_6_28_MUC_37 | Dinoflagellates | 0 | 0 | 0 |
| EMB262_6_28_MUC_37 | Diatoms | 1 | 2 | 1 |
| EMB262_6_28_MUC_38 | Dinoflagellates | 0 | 0 | 0 |
| EMB262_6_28_MUC_38 | Diatoms | 2 | 725 | 2 |
| EMB262_6_28_MUC_39 | Dinoflagellates | 0 | 0 | 0 |
| EMB262_6_28_MUC_39 | Diatoms | 3 | 325 | 3 |
| EMB262_6_28_MUC_40 | Dinoflagellates | 0 | 0 | 0 |
| EMB262_6_28_MUC_40 | Diatoms | 1 | 15 | 2 |
| EMB262_6_28_MUC_41 | Dinoflagellates | 0 | 0 | 0 |
| EMB262_6_28_MUC_41 | Diatoms | 0 | 0 | 0 |
| EMB262_6_28_MUC_42 | Dinoflagellates | 0 | 0 | 0 |
| EMB262_6_28_MUC_42 | Diatoms | 1 | 575 | 1 |
| EMB262_6_28_MUC_43 | Dinoflagellates | 0 | 0 | 0 |
| EMB262_6_28_MUC_43 | Diatoms | 1 | 275 | 1 |
| EMB262_6_28_MUC_44 | Dinoflagellates | 0 | 0 | 0 |
| EMB262_6_28_MUC_44 | Diatoms | 1 | 1 | 1 |
| EMB262_6_28_MUC_45 | Dinoflagellates | 0 | 0 | 0 |
| EMB262_6_28_MUC_45 | Diatoms | 1 | 35 | 1 |
| EMB262_6_28_MUC_46 | Dinoflagellates | 0 | 0 | 0 |
| EMB262_6_28_MUC_46 | Diatoms | 1 | 25 | 2 |
| EMB262_6_28_MUC_47 | Dinoflagellates | 1 | 25 | 1 |
| EMB262_6_28_MUC_47 | Diatoms | 1 | 25 | 1 |
| EMB262_6_28_MUC_48 | Dinoflagellates | 0 | 0 | 0 |
| EMB262_6_28_MUC_48 | Diatoms | 1 | 775 | 2 |
| EMB262_6_28_MUC_49 | Dinoflagellates | 0 | 0 | 0 |
| EMB262_6_28_MUC_49 | Diatoms | 1 | 175 | 1 |
| EMB262_6_28_MUC_5 | Dinoflagellates | 1 | 265 | 4 |
| EMB262_6_28_MUC_5 | Diatoms | 0 | 0 | 0 |
| EMB262_6_28_MUC_50 | Dinoflagellates | 0 | 0 | 0 |
| EMB262_6_28_MUC_50 | Diatoms | 0 | 0 | 0 |
| EMB262_6_28_MUC_51 | Dinoflagellates | 0 | 0 | 0 |
| EMB262_6_28_MUC_51 | Diatoms | 0 | 0 | 0 |
| EMB262_6_28_MUC_7 | Dinoflagellates | 1 | 91 | 2 |
| EMB262_6_28_MUC_7 | Diatoms | 0 | 0 | 0 |
| EMB262_6_28_MUC_9 | Dinoflagellates | 0 | 0 | 0 |
| EMB262_6_28_MUC_9 | Diatoms | 0 | 0 | 0 |
| EMB262_6_30_GC_1 | Dinoflagellates | 0 | 0 | 0 |
| EMB262_6_30_GC_1 | Diatoms | 0 | 0 | 0 |

| | | | | |
|---|---|---|---|---|
| EMB262_6_30_GC_10 | Dinoflagellates | 0 | 0 | 0 |
| EMB262_6_30_GC_10 | Diatoms | 0 | 0 | 0 |
| EMB262_6_30_GC_12 | Dinoflagellates | 1 | 75 | 1 |
| EMB262_6_30_GC_12 | Diatoms | 2 | 3 | 2 |
| EMB262_6_30_GC_13 | Dinoflagellates | 0 | 0 | 0 |
| EMB262_6_30_GC_13 | Diatoms | 2 | 2 | 2 |
| EMB262_6_30_GC_14 | Dinoflagellates | 0 | 0 | 0 |
| EMB262_6_30_GC_14 | Diatoms | 2 | 3 | 2 |
| EMB262_6_30_GC_15 | Dinoflagellates | 0 | 0 | 0 |
| EMB262_6_30_GC_15 | Diatoms | 1 | 25 | 1 |
| EMB262_6_30_GC_16 | Dinoflagellates | 0 | 0 | 0 |
| EMB262_6_30_GC_16 | Diatoms | 1 | 5 | 2 |
| EMB262_6_30_GC_17 | Dinoflagellates | 0 | 0 | 0 |
| EMB262_6_30_GC_17 | Diatoms | 0 | 0 | 0 |
| EMB262_6_30_GC_18 | Dinoflagellates | 0 | 0 | 0 |
| EMB262_6_30_GC_18 | Diatoms | 0 | 0 | 0 |
| EMB262_6_30_GC_19 | Dinoflagellates | 0 | 0 | 0 |
| EMB262_6_30_GC_19 | Diatoms | 2 | 3 | 2 |
| EMB262_6_30_GC_2 | Dinoflagellates | 0 | 0 | 0 |
| EMB262_6_30_GC_2 | Diatoms | 1 | 12 | 1 |
| EMB262_6_30_GC_20 | Dinoflagellates | 0 | 0 | 0 |
| EMB262_6_30_GC_20 | Diatoms | 2 | 875 | 4 |
| EMB262_6_30_GC_21 | Dinoflagellates | 0 | 0 | 0 |
| EMB262_6_30_GC_21 | Diatoms | 3 | 975 | 4 |
| EMB262_6_30_GC_22 | Dinoflagellates | 0 | 0 | 0 |
| EMB262_6_30_GC_22 | Diatoms | 0 | 0 | 0 |
| EMB262_6_30_GC_23 | Dinoflagellates | 0 | 0 | 0 |
| EMB262_6_30_GC_23 | Diatoms | 1 | 75 | 1 |
| EMB262_6_30_GC_24 | Dinoflagellates | 0 | 0 | 0 |
| EMB262_6_30_GC_24 | Diatoms | 4 | 325 | 5 |
| EMB262_6_30_GC_25 | Dinoflagellates | 0 | 0 | 0 |
| EMB262_6_30_GC_25 | Diatoms | 1 | 25 | 1 |
| EMB262_6_30_GC_26 | Dinoflagellates | 0 | 0 | 0 |
| EMB262_6_30_GC_26 | Diatoms | 0 | 0 | 0 |
| EMB262_6_30_GC_27 | Dinoflagellates | 0 | 0 | 0 |
| EMB262_6_30_GC_27 | Diatoms | 0 | 0 | 0 |
| EMB262_6_30_GC_29 | Dinoflagellates | 0 | 0 | 0 |
| EMB262_6_30_GC_29 | Diatoms | 0 | 0 | 0 |
| EMB262_6_30_GC_29 | Dinoflagellates | 1 | 5 | 1 |
| EMB262_6_30_GC_29 | Diatoms | 2 | 1825 | 8 |
| EMB262_6_30_GC_3 | Dinoflagellates | 0 | 0 | 0 |
| EMB262_6_30_GC_3 | Diatoms | 0 | 0 | 0 |
| EMB262_6_30_GC_31 | Dinoflagellates | 0 | 0 | 0 |
| EMB262_6_30_GC_31 | Diatoms | 1 | 475 | 2 |
| EMB262_6_30_GC_32 | Dinoflagellates | 0 | 0 | 0 |
| EMB262_6_30_GC_32 | Diatoms | 1 | 125 | 2 |
| EMB262_6_30_GC_33 | Dinoflagellates | 0 | 0 | 0 |

| | | | | |
|---|---|---|---|---|
| EMB262_6_30_GC_33 | Diatoms | 0 | 0 | 0 |
| EMB262_6_30_GC_35 | Dinoflagellates | 0 | 0 | 0 |
| EMB262_6_30_GC_35 | Diatoms | 0 | 0 | 0 |
| EMB262_6_30_GC_36 | Dinoflagellates | 0 | 0 | 0 |
| EMB262_6_30_GC_36 | Diatoms | 3 | 15975 | 7 |
| EMB262_6_30_GC_37 | Dinoflagellates | 0 | 0 | 0 |
| EMB262_6_30_GC_37 | Diatoms | 1 | 425 | 3 |
| EMB262_6_30_GC_38 | Dinoflagellates | 0 | 0 | 0 |
| EMB262_6_30_GC_38 | Diatoms | 2 | 4 | 2 |
| EMB262_6_30_GC_39 | Dinoflagellates | 0 | 0 | 0 |
| EMB262_6_30_GC_39 | Diatoms | 1 | 475 | 1 |
| EMB262_6_30_GC_4 | Dinoflagellates | 0 | 0 | 0 |
| EMB262_6_30_GC_4 | Diatoms | 0 | 0 | 0 |
| EMB262_6_30_GC_40 | Dinoflagellates | 0 | 0 | 0 |
| EMB262_6_30_GC_40 | Diatoms | 1 | 2 | 1 |
| EMB262_6_30_GC_41 | Dinoflagellates | 0 | 0 | 0 |
| EMB262_6_30_GC_41 | Diatoms | 1 | 225 | 1 |
| EMB262_6_30_GC_42 | Dinoflagellates | 0 | 0 | 0 |
| EMB262_6_30_GC_42 | Diatoms | 0 | 0 | 0 |
| EMB262_6_30_GC_43 | Dinoflagellates | 0 | 0 | 0 |
| EMB262_6_30_GC_43 | Diatoms | 2 | 3625 | 2 |
| EMB262_6_30_GC_43 | Dinoflagellates | 0 | 0 | 0 |
| EMB262_6_30_GC_43 | Diatoms | 0 | 0 | 0 |
| EMB262_6_30_GC_44 | Dinoflagellates | 0 | 0 | 0 |
| EMB262_6_30_GC_44 | Diatoms | 1 | 125 | 1 |
| EMB262_6_30_GC_45 | Dinoflagellates | 0 | 0 | 0 |
| EMB262_6_30_GC_45 | Diatoms | 0 | 0 | 0 |
| EMB262_6_30_GC_46 | Dinoflagellates | 0 | 0 | 0 |
| EMB262_6_30_GC_46 | Diatoms | 1 | 1025 | 3 |
| EMB262_6_30_GC_47 | Dinoflagellates | 0 | 0 | 0 |
| EMB262_6_30_GC_47 | Diatoms | 1 | 1225 | 4 |
| EMB262_6_30_GC_48 | Dinoflagellates | 0 | 0 | 0 |
| EMB262_6_30_GC_48 | Diatoms | 1 | 35 | 2 |
| EMB262_6_30_GC_49 | Dinoflagellates | 0 | 0 | 0 |
| EMB262_6_30_GC_49 | Diatoms | 2 | 1255 | 6 |
| EMB262_6_30_GC_5 | Dinoflagellates | 0 | 0 | 0 |
| EMB262_6_30_GC_5 | Diatoms | 0 | 0 | 0 |
| EMB262_6_30_GC_50 | Dinoflagellates | 0 | 0 | 0 |
| EMB262_6_30_GC_50 | Diatoms | 2 | 16775 | 8 |
| EMB262_6_30_GC_51 | Dinoflagellates | 0 | 0 | 0 |
| EMB262_6_30_GC_51 | Diatoms | 2 | 21375 | 8 |
| EMB262_6_30_GC_52 | Dinoflagellates | 0 | 0 | 0 |
| EMB262_6_30_GC_52 | Diatoms | 1 | 125 | 2 |
| EMB262_6_30_GC_53 | Dinoflagellates | 0 | 0 | 0 |
| EMB262_6_30_GC_53 | Diatoms | 1 | 125 | 3 |
| EMB262_6_30_GC_54 | Dinoflagellates | 0 | 0 | 0 |
| EMB262_6_30_GC_54 | Diatoms | 2 | 7475 | 6 |

| | | | | |
|---|---|---|---|---|
| EMB262_6_30_GC_55 | Dinoflagellates | 0 | 0 | 0 |
| EMB262_6_30_GC_55 | Diatoms | 3 | 2735 | 9 |
| EMB262_6_30_GC_56 | Dinoflagellates | 0 | 0 | 0 |
| EMB262_6_30_GC_56 | Diatoms | 1 | 25 | 1 |
| EMB262_6_30_GC_57 | Dinoflagellates | 0 | 0 | 0 |
| EMB262_6_30_GC_57 | Diatoms | 0 | 0 | 0 |
| EMB262_6_30_GC_58 | Dinoflagellates | 0 | 0 | 0 |
| EMB262_6_30_GC_58 | Diatoms | 0 | 0 | 0 |
| EMB262_6_30_GC_59 | Dinoflagellates | 0 | 0 | 0 |
| EMB262_6_30_GC_59 | Diatoms | 0 | 0 | 0 |
| EMB262_6_30_GC_6 | Dinoflagellates | 0 | 0 | 0 |
| EMB262_6_30_GC_6 | Diatoms | 0 | 0 | 0 |
| EMB262_6_30_GC_60 | Dinoflagellates | 1 | 25 | 1 |
| EMB262_6_30_GC_60 | Diatoms | 0 | 0 | 0 |
| EMB262_6_30_GC_61 | Dinoflagellates | 0 | 0 | 0 |
| EMB262_6_30_GC_61 | Diatoms | 1 | 75 | 1 |
| EMB262_6_30_GC_62 | Dinoflagellates | 0 | 0 | 0 |
| EMB262_6_30_GC_62 | Diatoms | 0 | 0 | 0 |
| EMB262_6_30_GC_63 | Dinoflagellates | 0 | 0 | 0 |
| EMB262_6_30_GC_63 | Diatoms | 1 | 1925 | 2 |
| EMB262_6_30_GC_64 | Dinoflagellates | 0 | 0 | 0 |
| EMB262_6_30_GC_64 | Diatoms | 1 | 625 | 2 |
| EMB262_6_30_GC_65 | Dinoflagellates | 0 | 0 | 0 |
| EMB262_6_30_GC_65 | Diatoms | 1 | 21 | 2 |
| EMB262_6_30_GC_66 | Dinoflagellates | 0 | 0 | 0 |
| EMB262_6_30_GC_66 | Diatoms | 1 | 2 | 2 |
| EMB262_6_30_GC_7 | Dinoflagellates | 0 | 0 | 0 |
| EMB262_6_30_GC_7 | Diatoms | 0 | 0 | 0 |
| EMB262_6_30_GC_8 | Dinoflagellates | 0 | 0 | 0 |
| EMB262_6_30_GC_8 | Diatoms | 0 | 0 | 0 |

#### Droplet digital PCR

Table S6 presents the ddPCR concentration measured in copies/µL for both targets *A. malmogiense* (FAM) and *S. marinoi* (HEX). Each sample tagged by a unique identifier (tag) was run in three replicates. The concentration was normalized via its corresponding sediment weight and the average concentration of the three replicates.

**Table S6: ddPCR of Dinoflagellates and Diatoms from DNA extracts of EGB262 samples. Each sample from each location is uniquely tagged (tag).**

| tag | conc. FAM [copies/µL] | conc. HEX [copies/µL] | sediment weight [mg] | relative conc. FAM | $\bar{X}$ conc. Am [copies/µL] | relative conc. HEX | $\bar{X}$ conc. Sm [copies/µL] |
|---|---|---|---|---|---|---|---|
| EMB262_12_3_GC_41 | 19,6 | 2,9 | 421,2 | 23,27 | 22,61 | 3,44 | 5,22 |
| EMB262_12_3_GC_41 | 18,5 | 5,9 | 421,2 | 21,96 | 22,61 | 7 | 5,22 |
| EMB262_6_30_GC_15 | 0 | 4,9 | 433 | 0 | 0 | 5,66 | 6,27 |
| EMB262_6_30_GC_15 | 0 | 5,6 | 433 | 0 | 0 | 6,47 | 6,27 |

| | | | | | | | |
|---|---|---|---|---|---|---|---|
| EMB262_6_30_GC_15 | 0 | 5,8 | 433 | 0 | 0 | 6,7 | 6,27 |
| EMB262_6_30_GC_25 | 0 | 8,6 | 445,5 | 0 | 0 | 9,65 | 11,04 |
| EMB262_6_30_GC_25 | 0 | 11,7 | 445,5 | 0 | 0 | 13,13 | 11,04 |
| EMB262_6_30_GC_25 | 0 | 9,2 | 445,5 | 0 | 0 | 10,33 | 11,04 |
| EMB262_12_3_GC_27 | 0,16 | 2,3 | 432,7 | 0,18 | 0,18 | 2,66 | 2,45 |
| EMB262_12_3_GC_27 | 0,3 | 3,9 | 432,7 | 0,35 | 0,18 | 4,51 | 2,45 |
| EMB262_12_3_GC_27 | 0 | 0,16 | 432,7 | 0 | 0,18 | 0,18 | 2,45 |
| EMB262_6_30_GC_37 | 0 | 1,4 | 511,9 | 0 | 0 | 1,37 | 1,56 |
| EMB262_6_30_GC_37 | 0 | 1,6 | 511,9 | 0 | 0 | 1,56 | 1,56 |
| EMB262_6_30_GC_37 | 0 | 1,8 | 511,9 | 0 | 0 | 1,76 | 1,56 |
| EMB262_6_30_GC_51 | 0 | 15,3 | 485,7 | 0 | 0,13 | 15,75 | 17,16 |
| EMB262_6_30_GC_51 | 0,14 | 18 | 485,7 | 0,14 | 0,13 | 18,53 | 17,16 |
| EMB262_6_30_GC_51 | 0,23 | 16,7 | 485,7 | 0,24 | 0,13 | 17,19 | 17,16 |
| EMB262_12_3_GC_7 | 0,3 | 6,2 | 465,7 | 0,32 | 0,97 | 6,66 | 5,94 |
| EMB262_12_3_GC_7 | 1 | 4,1 | 465,7 | 1,07 | 0,97 | 4,4 | 5,94 |
| EMB262_12_3_GC_7 | 1,4 | 6,3 | 465,7 | 1,5 | 0,97 | 6,76 | 5,94 |
| EMB262_6_30_GC_62 | 0 | 1,1 | 627,5 | 0 | 0 | 0,88 | 0,73 |
| EMB262_6_30_GC_62 | 0 | 0,35 | 627,5 | 0 | 0 | 0,28 | 0,73 |
| EMB262_6_30_GC_62 | 0 | 1,3 | 627,5 | 0 | 0 | 1,04 | 0,73 |
| NTC | 0 | 0 | 0 | 0 | | 0 | |
| EMB262_3_10_MUC_4 | 68 | 54 | 546 | 62,27 | 63,49 | 49,45 | 52,5 |
| EMB262_3_10_MUC_4 | 85 | 70 | 546 | 77,84 | 63,49 | 64,1 | 52,5 |
| EMB262_3_10_MUC_4 | 55 | 48 | 546 | 50,37 | 63,49 | 43,96 | 52,5 |
| EMB262_3_10_MUC_5 | 31,8 | 1,1 | 539 | 29,5 | 19,94 | 1,02 | 0,69 |
| EMB262_3_10_MUC_5 | 17,4 | 0,9 | 539 | 16,14 | 19,94 | 0,83 | 0,69 |
| EMB262_3_10_MUC_5 | 15,3 | 0,23 | 539 | 14,19 | 19,94 | 0,21 | 0,69 |
| EMB262_3_10_MUC_6 | 14,3 | 0,27 | 594 | 12,04 | 8,28 | 0,23 | 0,11 |
| EMB262_3_10_MUC_6 | 7,3 | 0 | 594 | 6,14 | 8,28 | 0 | 0,11 |
| EMB262_3_10_MUC_6 | 7,9 | 0,12 | 594 | 6,65 | 8,28 | 0,1 | 0,11 |
| EMB262_3_10_MUC_14 | 5,4 | 0,23 | 758 | 3,56 | 2,62 | 0,15 | 0,11 |
| EMB262_3_10_MUC_14 | 4,4 | 0,14 | 758 | 2,9 | 2,62 | 0,09 | 0,11 |
| EMB262_3_10_MUC_14 | 2,1 | 0,15 | 758 | 1,39 | 2,62 | 0,1 | 0,11 |
| EMB262_3_10_MUC_12 | 212 | 1,5 | 620 | 170,97 | 128,49 | 1,21 | 0,83 |
| EMB262_3_10_MUC_12 | 143 | 0,9 | 620 | 115,32 | 128,49 | 0,73 | 0,83 |
| EMB262_3_10_MUC_12 | 123 | 0,7 | 620 | 99,19 | 128,49 | 0,56 | 0,83 |
| EMB262_3_10_MUC_10 | 10,7 | 1,1 | 252 | 21,23 | 16,8 | 2,18 | 1,65 |
| EMB262_3_10_MUC_10 | 7,4 | 0,6 | 252 | 14,68 | 16,8 | 1,19 | 1,65 |
| EMB262_3_10_MUC_10 | 7,3 | 0,8 | 252 | 14,48 | 16,8 | 1,59 | 1,65 |
| EMB262_6_28_MUC_14 | 0 | 4,3 | 738 | 0 | 0 | 2,91 | 1,92 |
| EMB262_6_28_MUC_14 | 0 | 2,4 | 738 | 0 | 0 | 1,63 | 1,92 |
| EMB262_6_28_MUC_14 | 0 | 1,8 | 738 | 0 | 0 | 1,22 | 1,92 |
| EMB262_6_28_MUC_1 | 23,9 | 12,9 | 485 | 24,64 | 27,46 | 13,3 | 12,2 |
| EMB262_6_28_MUC_1 | 35 | 9,6 | 485 | 36,08 | 27,46 | 9,9 | 12,2 |
| EMB262_6_28_MUC_1 | 21 | 13 | 485 | 21,65 | 27,46 | 13,4 | 12,2 |
| EMB262_6_28_MUC_5 | 51 | 0,7 | 599 | 42,57 | 35,2 | 0,58 | 1,09 |
| EMB262_6_28_MUC_5 | 41,3 | 1,4 | 599 | 34,47 | 35,2 | 1,17 | 1,09 |
| EMB262_6_28_MUC_5 | 34,2 | 1,8 | 599 | 28,55 | 35,2 | 1,5 | 1,09 |

| | | | | | | | |
|---|---|---|---|---|---|---|---|
| NTC | 0 | 0 | - | 0 | | 0 | |
| EMB262_6_28_MUC_12 | 0 | 1,9 | 675 | 0 | 0,04 | 1,41 | 0,94 |
| EMB262_6_28_MUC_12 | 0 | 1,4 | 675 | 0 | 0,04 | 1,04 | 0,94 |
| EMB262_6_28_MUC_12 | 0,17 | 0,5 | 675 | 0,13 | 0,04 | 0,37 | 0,94 |
| EMB262_12_2_MUC_11 | 9 | 2,6 | 723 | 6,22 | 4,5 | 1,8 | 1,01 |
| EMB262_12_2_MUC_11 | 6 | 0,9 | 723 | 4,15 | 4,5 | 0,62 | 1,01 |
| EMB262_12_2_MUC_11 | 4,5 | 0,9 | 723 | 3,11 | 4,5 | 0,62 | 1,01 |
| EMB262_12_2_MUC_12 | 0 | 1,7 | 598 | 0 | 0 | 1,42 | 1,64 |
| EMB262_12_2_MUC_12 | 0 | 2,1 | 598 | 0 | 0 | 1,76 | 1,64 |
| EMB262_12_2_MUC_12 | 0 | 2,1 | 598 | 0 | 0 | 1,76 | 1,64 |
| EMB262_12_2_MUC_9 | 0 | 0 | 578 | 0 | 62,57 | 0 | 1,99 |
| EMB262_12_2_MUC_9 | 127 | 4,3 | 578 | 109,86 | 62,57 | 3,72 | 1,99 |
| EMB262_12_2_MUC_9 | 90 | 2,6 | 578 | 77,85 | 62,57 | 2,25 | 1,99 |
| EMB262_13_8_MUC_21 | 0 | 0,8 | 957 | 0 | 0 | 0,42 | 0,24 |
| EMB262_13_8_MUC_21 | 0 | 0,1 | 957 | 0 | 0 | 0,05 | 0,24 |
| EMB262_13_8_MUC_21 | 0 | 0,45 | 957 | 0 | 0 | 0,24 | 0,24 |
| EMB262_13_8_MUC_22 | 0,08 | 0,7 | 761 | 0,05 | 0,07 | 0,46 | 0,48 |
| EMB262_13_8_MUC_22 | 0 | 0,9 | 761 | 0 | 0,07 | 0,59 | 0,48 |
| EMB262_13_8_MUC_22 | 0,23 | 0,6 | 761 | 0,15 | 0,07 | 0,39 | 0,48 |
| EMB262_13_8_MUC_26 | 253 | 1,4 | 649 | 194,92 | 137,9 | 1,08 | 0,85 |
| EMB262_13_8_MUC_26 | 156 | 1 | 649 | 120,18 | 137,9 | 0,77 | 0,85 |
| EMB262_13_8_MUC_26 | 128 | 0,9 | 649 | 98,61 | 137,9 | 0,69 | 0,85 |
| EMB262_13_8_MUC_23 | 0 | 1,4 | 983 | 0 | 0,04 | 0,71 | 0,51 |
| EMB262_13_8_MUC_23 | 0 | 1,1 | 983 | 0 | 0,04 | 0,56 | 0,51 |
| EMB262_13_8_MUC_23 | 0,25 | 0,5 | 983 | 0,13 | 0,04 | 0,25 | 0,51 |
| EMB262_13_8_MUC_25 | 10,2 | 0,8 | 797 | 6,4 | 3,72 | 0,5 | 0,46 |
| EMB262_13_8_MUC_25 | 4,9 | 0,52 | 797 | 3,07 | 3,72 | 0,33 | 0,46 |
| EMB262_13_8_MUC_25 | 2,7 | 0,9 | 797 | 1,69 | 3,72 | 0,56 | 0,46 |
| NTC | 0 | 0 | - | 0 | | 0 | |
| EMB262_6_28_MUC_16 | 0 | 0,4 | 741 | 0 | 0 | 0,27 | 0,34 |
| EMB262_6_28_MUC_16 | 0 | 0,7 | 741 | 0 | 0 | 0,47 | 0,34 |
| EMB262_6_28_MUC_16 | 0 | 0,4 | 741 | 0 | 0 | 0,27 | 0,34 |
| EMB262_6_28_MUC_7 | 6,8 | 0 | 769 | 4,42 | 5,2 | 0 | 0 |
| EMB262_6_28_MUC_7 | 7,2 | 0 | 769 | 4,68 | 5,2 | 0 | 0 |
| EMB262_6_28_MUC_7 | 10 | 0 | 769 | 6,5 | 5,2 | 0 | 0 |
| EMB262_3_10_MUC_9 | 90 | 4,3 | 781 | 57,62 | 41,19 | 2,75 | 2,3 |
| EMB262_3_10_MUC_9 | 50 | 3,4 | 781 | 32,01 | 41,19 | 2,18 | 2,3 |
| EMB262_3_10_MUC_9 | 53 | 3,1 | 781 | 33,93 | 41,19 | 1,98 | 2,3 |
| EMB262_12_2_MUC_13 | 0 | 2,3 | 748 | 0 | 0 | 1,54 | 1,09 |
| EMB262_12_2_MUC_13 | 0 | 1,2 | 748 | 0 | 0 | 0,8 | 1,09 |
| EMB262_12_2_MUC_13 | 0 | 1,4 | 748 | 0 | 0 | 0,94 | 1,09 |
| EMB262_3_10_MUC_15 | 18,8 | 4,6 | 972 | 9,67 | 6,45 | 2,37 | 1,39 |
| EMB262_3_10_MUC_15 | 11,7 | 2,2 | 972 | 6,02 | 6,45 | 1,13 | 1,39 |
| EMB262_3_10_MUC_15 | 7,1 | 1,3 | 972 | 3,65 | 6,45 | 0,67 | 1,39 |
| EMB262_3_10_MUC_16 | 14,6 | 1,2 | 472 | 15,47 | 42,44 | 1,27 | 0,99 |
| EMB262_3_10_MUC_16 | 11,6 | 0,8 | 472 | 12,29 | 42,44 | 0,85 | 0,99 |
| EMB262_3_10_MUC_16 | 94 | 0,8 | 472 | 99,58 | 42,44 | 0,85 | 0,99 |

| | | | | | | | |
|---|---|---|---|---|---|---|---|
| EMB262_3_10_MUC_3 | 36,6 | 109 | 380 | 48,16 | 29,12 | 143,42 | 92,59 |
| EMB262_3_10_MUC_3 | 15 | 58,1 | 380 | 19,74 | 29,12 | 76,45 | 92,59 |
| EMB262_3_10_MUC_3 | 14,8 | 44 | 380 | 19,47 | 29,12 | 57,89 | 92,59 |
| EMB262_12_2_MUC_1 | 263 | 58,7 | 737 | 178,43 | 116,01 | 39,82 | 27,48 |
| EMB262_12_2_MUC_1 | 141 | 36,8 | 737 | 95,66 | 116,01 | 24,97 | 27,48 |
| EMB262_12_2_MUC_1 | 109 | 26 | 737 | 73,95 | 116,01 | 17,64 | 27,48 |
| EMB262_6_28_MUC_13 | 0 | 1,1 | 859 | 0 | 0 | 0,64 | 0,44 |
| EMB262_6_28_MUC_13 | 0 | 0,8 | 859 | 0 | 0 | 0,47 | 0,44 |
| EMB262_6_28_MUC_13 | 0 | 0,35 | 859 | 0 | 0 | 0,2 | 0,44 |
| NTC | 0 | 0 | - | 0 | | 0 | |
| EMB262_13_8_MUC_19 | 23 | 5,4 | 561 | 20,5 | 6,88 | 4,81 | 3,54 |
| EMB262_13_8_MUC_19 | 0 | 3 | 561 | 0 | 6,88 | 2,67 | 3,54 |
| EMB262_13_8_MUC_19 | 0,15 | 3,5 | 561 | 0,13 | 6,88 | 3,12 | 3,54 |
| EMB262_3_10_MUC_19 | 207 | 1291 | 631 | 164,03 | 154,25 | 1022,98 | 975,44 |
| EMB262_3_10_MUC_19 | 223 | 1400 | 631 | 176,7 | 154,25 | 1109,35 | 975,44 |
| EMB262_3_10_MUC_19 | 154 | 1002 | 631 | 122,03 | 154,25 | 793,98 | 975,44 |
| EMB262_13_8_MUC_18 | 17,9 | 3,3 | 624 | 14,34 | 10,98 | 2,64 | 2,32 |
| EMB262_13_8_MUC_18 | 12,9 | 3,5 | 624 | 10,34 | 10,98 | 2,8 | 2,32 |
| EMB262_13_8_MUC_18 | 10,3 | 1,9 | 624 | 8,25 | 10,98 | 1,52 | 2,32 |
| EMB262_3_10_MUC_2 | 56 | 202 | 640 | 43,75 | 42,63 | 157,81 | 144,27 |
| EMB262_3_10_MUC_2 | 57,9 | 187 | 640 | 45,23 | 42,63 | 146,09 | 144,27 |
| EMB262_3_10_MUC_2 | 49,8 | 165 | 640 | 38,91 | 42,63 | 128,91 | 144,27 |
| EMB262_6_28_MUC_18 | 0,07 | 0,35 | 557 | 0,06 | 0,02 | 0,31 | 0,29 |
| EMB262_6_28_MUC_18 | 0 | 0,36 | 557 | 0 | 0,02 | 0,32 | 0,29 |
| EMB262_6_28_MUC_18 | 0 | 0,26 | 557 | 0 | 0,02 | 0,23 | 0,29 |
| EMB262_3_10_MUC_16 | 25,1 | 2,4 | 432 | 29,05 | 20,22 | 2,78 | 2,47 |
| EMB262_3_10_MUC_16 | 14,5 | 2,4 | 432 | 16,78 | 20,22 | 2,78 | 2,47 |
| EMB262_3_10_MUC_16 | 12,8 | 1,6 | 432 | 14,81 | 20,22 | 1,85 | 2,47 |
| EMB262_12_2_MUC_5 | 136 | 1,9 | 514 | 132,3 | 172,5 | 1,85 | 1,39 |
| EMB262_12_2_MUC_5 | 108 | 1 | 514 | 105,06 | 172,5 | 0,97 | 1,39 |
| EMB262_12_2_MUC_5 | 288 | 1,4 | 514 | 280,16 | 172,5 | 1,36 | 1,39 |
| EMB262_12_2_MUC_17 | 6,5 | 1,5 | 486 | 6,69 | 5,9 | 1,54 | 1,2 |
| EMB262_12_2_MUC_17 | 5,8 | 1,1 | 486 | 5,97 | 5,9 | 1,13 | 1,2 |
| EMB262_12_2_MUC_17 | 4,9 | 0,9 | 486 | 5,04 | 5,9 | 0,93 | 1,2 |
| EMB262_3_10_MUC_17 | 38 | 2,5 | 709 | 26,8 | 18,71 | 1,76 | 1,18 |
| EMB262_3_10_MUC_17 | 16,5 | 1 | 709 | 11,64 | 18,71 | 0,71 | 1,18 |
| EMB262_3_10_MUC_17 | 25,1 | 1,5 | 709 | 17,7 | 18,71 | 1,06 | 1,18 |
| NTC | 0 | 0 | - | 0 | | 0 | |
| EMB262_13_8_MUC_17 | 49,6 | 1 | 512 | 48,44 | 33,56 | 0,98 | 0,54 |
| EMB262_13_8_MUC_17 | 34,4 | 0,26 | 512 | 33,59 | 33,56 | 0,25 | 0,54 |
| EMB262_13_8_MUC_17 | 19,1 | 0,39 | 512 | 18,65 | 33,56 | 0,38 | 0,54 |
| EMB262_12_2_MUC_16 | 0,3 | 0,44 | 635 | 0,24 | 0,13 | 0,35 | 0,34 |
| EMB262_12_2_MUC_16 | 0,12 | 0,6 | 635 | 0,09 | 0,13 | 0,47 | 0,34 |
| EMB262_12_2_MUC_16 | 0,07 | 0,27 | 635 | 0,06 | 0,13 | 0,21 | 0,34 |
| EMB262_6_28_MUC_3 | 34,3 | 1,2 | 657 | 26,1 | 15,53 | 0,91 | 0,53 |
| EMB262_6_28_MUC_3 | 15,7 | 0,57 | 657 | 11,95 | 15,53 | 0,43 | 0,53 |
| EMB262_6_28_MUC_3 | 11,2 | 0,3 | 657 | 8,52 | 15,53 | 0,23 | 0,53 |

| | | | | | | | |
|---|---|---|---|---|---|---|---|
| EMB262_12_2_MUC_7 | 308 | 2,3 | 652 | 236,2 | 150,56 | 1,76 | 1,33 |
| EMB262_12_2_MUC_7 | 143 | 1,7 | 652 | 109,66 | 150,56 | 1,3 | 1,33 |
| EMB262_12_2_MUC_7 | 138 | 1,2 | 652 | 105,83 | 150,56 | 0,92 | 1,33 |
| EMB262_3_10_MUC_13 | 43,2 | 0,68 | 480 | 45 | 37,43 | 0,71 | 0,65 |
| EMB262_3_10_MUC_13 | 32,9 | 0,65 | 480 | 34,27 | 37,43 | 0,68 | 0,65 |
| EMB262_3_10_MUC_13 | 31,7 | 0,53 | 480 | 33,02 | 37,43 | 0,55 | 0,65 |
| EMB262_3_10_MUC_20 | 11,4 | 0,33 | 547 | 10,42 | 7,46 | 0,3 | 0,26 |
| EMB262_3_10_MUC_20 | 6,7 | 0,29 | 547 | 6,12 | 7,46 | 0,27 | 0,26 |
| EMB262_3_10_MUC_20 | 6,4 | 0,22 | 547 | 5,85 | 7,46 | 0,2 | 0,26 |
| EMB262_12_2_MUC_3 | 495 | 1 | 681 | 363,44 | 331,13 | 0,73 | 0,62 |
| EMB262_12_2_MUC_3 | 443 | 0,64 | 681 | 325,26 | 331,13 | 0,47 | 0,62 |
| EMB262_12_2_MUC_3 | 415 | 0,9 | 681 | 304,7 | 331,13 | 0,66 | 0,62 |
| EMB262_6_28_MUC_20 | 0 | 0,52 | 705 | 0 | 0 | 0,37 | 0,48 |
| EMB262_6_28_MUC_20 | 0 | 0,72 | 705 | 0 | 0 | 0,51 | 0,48 |
| EMB262_6_28_MUC_20 | 0 | 0,81 | 705 | 0 | 0 | 0,57 | 0,48 |
| EMB262_12_2_MUC_31 | 0 | 0,07 | 274 | 0 | 0 | 0,13 | 0,35 |
| EMB262_12_2_MUC_31 | 0 | 0,28 | 274 | 0 | 0 | 0,51 | 0,35 |
| EMB262_12_2_MUC_31 | 0 | 0,22 | 274 | 0 | 0 | 0,4 | 0,35 |
| NTC | 0 | 0 | - | 0 | | 0 | |
| EMB262_12_2_MUC_15 | 0 | 0,53 | 689 | 0 | 0 | 0,38 | 0,32 |
| EMB262_12_2_MUC_15 | 0 | 0,28 | 689 | 0 | 0 | 0,2 | 0,32 |
| EMB262_12_2_MUC_15 | 0 | 0,51 | 689 | 0 | 0 | 0,37 | 0,32 |
| EMB262_12_2_MUC_19 | 0 | 1,4 | 793 | 0 | 0 | 0,88 | 0,63 |
| EMB262_12_2_MUC_19 | 0 | 1 | 793 | 0 | 0 | 0,63 | 0,63 |
| EMB262_12_2_MUC_19 | 0 | 0,59 | 793 | 0 | 0 | 0,37 | 0,63 |
| EMB262_3_10_MUC_36 | 0 | 0,6 | 648 | 0 | 0 | 0,46 | 0,27 |
| EMB262_3_10_MUC_36 | 0 | 0,33 | 648 | 0 | 0 | 0,25 | 0,27 |
| EMB262_3_10_MUC_36 | 0 | 0,12 | 648 | 0 | 0 | 0,09 | 0,27 |
| EMB262_3_10_MUC_19 | 0 | 0,32 | 486 | 0 | 0 | 0,33 | 0,11 |
| EMB262_3_10_MUC_19 | 0 | 0 | 486 | 0 | 0 | 0 | 0,11 |
| EMB262_3_10_MUC_19 | 0 | 0 | 486 | 0 | 0 | 0 | 0,11 |
| EMB262_12_2_MUC_21 | 0 | 0 | 512 | 0 | 0,02 | 0 | 0,1 |
| EMB262_12_2_MUC_21 | 0 | 0,17 | 512 | 0 | 0,02 | 0,17 | 0,1 |
| EMB262_12_2_MUC_21 | 0,07 | 0,15 | 512 | 0,07 | 0,02 | 0,15 | 0,1 |
| EMB262_3_10_MUC_39 | 0 | 0,13 | 525 | 0 | 0 | 0,12 | 0,1 |
| EMB262_3_10_MUC_39 | 0 | 0,06 | 525 | 0 | 0 | 0,06 | 0,1 |
| EMB262_3_10_MUC_39 | 0 | 0,13 | 525 | 0 | 0 | 0,12 | 0,1 |
| EMB262_6_28_MUC_21 | 0 | 1,4 | 735 | 0 | 0 | 0,95 | 0,53 |
| EMB262_6_28_MUC_21 | 0 | 0,4 | 735 | 0 | 0 | 0,27 | 0,53 |
| EMB262_6_28_MUC_21 | 0 | 0,52 | 735 | 0 | 0 | 0,35 | 0,53 |
| EMB262_3_10_MUC_31 | 0 | 0,3 | 632 | 0 | 0 | 0,24 | 0,18 |
| EMB262_3_10_MUC_31 | 0 | 0,33 | 632 | 0 | 0 | 0,26 | 0,18 |
| EMB262_3_10_MUC_31 | 0 | 0,07 | 632 | 0 | 0 | 0,06 | 0,18 |
| EMB262_12_2_MUC_28 | 0,08 | 0,16 | 574 | 0,07 | 0,02 | 0,14 | 0,12 |
| EMB262_12_2_MUC_28 | 0 | 0,08 | 574 | 0 | 0,02 | 0,07 | 0,12 |
| EMB262_12_2_MUC_28 | 0 | 0,16 | 574 | 0 | 0,02 | 0,14 | 0,12 |
| NTC | 0 | 0 | - | 0 | | 0 | |

| | | | | | | | |
|---|---|---|---|---|---|---|---|
| EMB262_3_10_MUC_35 | 0 | 0,46 | 384 | 0 | 0 | 0,6 | 0,51 |
| EMB262_3_10_MUC_35 | 0 | 0,42 | 384 | 0 | 0 | 0,55 | 0,51 |
| EMB262_3_10_MUC_35 | 0 | 0,3 | 384 | 0 | 0 | 0,39 | 0,51 |
| EMB262_13_8_MUC_14 | 0,34 | 7,6 | 847 | 0,2 | 0,18 | 4,49 | 3,78 |
| EMB262_13_8_MUC_14 | 0,22 | 6 | 847 | 0,13 | 0,18 | 3,54 | 3,78 |
| EMB262_13_8_MUC_14 | 0,35 | 5,6 | 847 | 0,21 | 0,18 | 3,31 | 3,78 |
| EMB262_6_28_MUC_17 | 0 | 3,3 | 825 | 0 | 0 | 2 | 1,27 |
| EMB262_6_28_MUC_17 | 0 | 1,4 | 825 | 0 | 0 | 0,85 | 1,27 |
| EMB262_6_28_MUC_17 | 0 | 1,6 | 825 | 0 | 0 | 0,97 | 1,27 |
| EMB262_3_10_MUC_38 | 0 | 0,47 | 873 | 0 | 0 | 0,27 | 0,19 |
| EMB262_3_10_MUC_38 | 0 | 0,22 | 873 | 0 | 0 | 0,13 | 0,19 |
| EMB262_3_10_MUC_38 | 0 | 0,31 | 873 | 0 | 0 | 0,18 | 0,19 |
| EMB262_3_10_MUC_26 | 0 | 0,07 | 418 | 0 | 0 | 0,08 | 0,03 |
| EMB262_3_10_MUC_26 | 0 | 0 | 418 | 0 | 0 | 0 | 0,03 |
| EMB262_3_10_MUC_26 | 0 | 0 | 418 | 0 | 0 | 0 | 0,03 |
| EMB262_12_2_MUC_24 | 0 | 0,13 | 619 | 0 | 0 | 0,11 | 0,06 |
| EMB262_12_2_MUC_24 | 0 | 0,08 | 619 | 0 | 0 | 0,06 | 0,06 |
| EMB262_12_2_MUC_24 | 0 | 0 | 619 | 0 | 0 | 0 | 0,06 |
| EMB262_12_2_MUC_23 | 0 | 0,14 | 919 | 0 | 0 | 0,08 | 0,05 |
| EMB262_12_2_MUC_23 | 0 | 0,07 | 919 | 0 | 0 | 0,04 | 0,05 |
| EMB262_12_2_MUC_23 | 0 | 0,07 | 919 | 0 | 0 | 0,04 | 0,05 |
| EMB262_12_2_MUC_22 | 0 | 0,07 | 529 | 0 | 0 | 0,07 | 0,02 |
| EMB262_12_2_MUC_22 | 0 | 0 | 529 | 0 | 0 | 0 | 0,02 |
| EMB262_12_2_MUC_22 | 0 | 0 | 529 | 0 | 0 | 0 | 0,02 |
| EMB262_3_10_MUC_29 | 0 | 0 | 603 | 0 | 0 | 0 | 0,02 |
| EMB262_3_10_MUC_29 | 0 | 0 | 603 | 0 | 0 | 0 | 0,02 |
| EMB262_3_10_MUC_29 | 0 | 0,06 | 603 | 0 | 0 | 0,05 | 0,02 |
| NTC | 0 | 0 | - | 0 | | 0 | |
| EMB262_6_28_MUC_23 | 0 | 0,2 | 889 | 0 | 0 | 0,11 | 0,12 |
| EMB262_6_28_MUC_23 | 0 | 0,39 | 889 | 0 | 0 | 0,22 | 0,12 |
| EMB262_6_28_MUC_23 | 0 | 0,06 | 889 | 0 | 0 | 0,03 | 0,12 |
| EMB262_3_10_MUC_30 | 0 | 0 | 886 | 0 | 0 | 0 | 0,05 |
| EMB262_3_10_MUC_30 | 0 | 0,25 | 886 | 0 | 0 | 0,14 | 0,05 |
| EMB262_3_10_MUC_30 | 0 | 0 | 886 | 0 | 0 | 0 | 0,05 |
| EMB262_6_28_MUC_22 | 0 | 0,48 | 800 | 0 | 0 | 0,3 | 0,19 |
| EMB262_6_28_MUC_22 | 0 | 0,08 | 800 | 0 | 0 | 0,05 | 0,19 |
| EMB262_6_28_MUC_22 | 0 | 0,33 | 800 | 0 | 0 | 0,21 | 0,19 |
| EMB262_12_2_MUC_34 | 0 | 0 | 511 | 0 | 0 | 0 | 0,04 |
| EMB262_12_2_MUC_34 | 0 | 0,06 | 511 | 0 | 0 | 0,06 | 0,04 |
| EMB262_12_2_MUC_34 | 0 | 0,07 | 511 | 0 | 0 | 0,07 | 0,04 |
| EMB262_13_8_MUC_9 | 0 | 0,07 | 882 | 0 | 0 | 0,04 | 0,03 |
| EMB262_13_8_MUC_9 | 0 | 0 | 882 | 0 | 0 | 0 | 0,03 |
| EMB262_13_8_MUC_9 | 0 | 0,07 | 882 | 0 | 0 | 0,04 | 0,03 |
| EMB262_6_28_MUC_25 | 0 | 1,6 | 740 | 0 | 0 | 1,08 | 0,6 |
| EMB262_6_28_MUC_25 | 0 | 0,7 | 740 | 0 | 0 | 0,47 | 0,6 |
| EMB262_6_28_MUC_25 | 0 | 0,36 | 740 | 0 | 0 | 0,24 | 0,6 |
| EMB262_6_28_MUC_34 | 0 | 0,29 | 839 | 0 | 0 | 0,17 | 0,09 |

| | | | | | | | |
|---|---|---|---|---|---|---|---|
| EMB262_6_28_MUC_34 | 0 | 0,16 | 839 | 0 | 0 | 0,1 | 0,09 |
| EMB262_6_28_MUC_34 | 0 | 0 | 839 | 0 | 0 | 0 | 0,09 |
| EMB262_6_28_MUC_11 | 0 | 0,1 | 767 | 0 | 0 | 0,07 | 0,07 |
| EMB262_6_28_MUC_11 | 0 | 0,1 | 767 | 0 | 0 | 0,07 | 0,07 |
| EMB262_6_28_MUC_11 | 0 | 0,1 | 767 | 0 | 0 | 0,07 | 0,07 |
| EMB262_3_10_MUC_37 | 0 | 0 | 799 | 0 | 0 | 0 | 0 |
| EMB262_3_10_MUC_37 | 0 | 0 | 799 | 0 | 0 | 0 | 0 |
| EMB262_3_10_MUC_37 | 0 | 0 | 799 | 0 | 0 | 0 | 0 |
| NTC | 0 | 0 | - | 0 | | 0 | |
| EMB262_3_10_MUC_27 | 0 | 0 | 614 | 0 | 0 | 0 | 0 |
| EMB262_3_10_MUC_27 | 0 | 0 | 614 | 0 | 0 | 0 | 0 |
| EMB262_3_10_MUC_27 | 0 | 0 | 614 | 0 | 0 | 0 | 0 |
| EMB262_3_10_MUC_8 | 32,1 | 1,2 | 534 | 30,06 | 28,93 | 1,12 | 0,59 |
| EMB262_3_10_MUC_8 | 31,8 | 0,4 | 534 | 29,78 | 28,93 | 0,37 | 0,59 |
| EMB262_3_10_MUC_8 | 28,8 | 0,29 | 534 | 26,97 | 28,93 | 0,27 | 0,59 |
| EMB262_12_2_MUC_26 | 0 | 0,22 | 577 | 0 | 0 | 0,19 | 0,27 |
| EMB262_12_2_MUC_26 | 0 | 0,28 | 577 | 0 | 0 | 0,24 | 0,27 |
| EMB262_12_2_MUC_26 | 0 | 0,42 | 577 | 0 | 0 | 0,36 | 0,27 |
| EMB262_3_10_MUC_32 | 0 | 0,27 | 754 | 0 | 0 | 0,18 | 0,1 |
| EMB262_3_10_MUC_32 | 0 | 0,13 | 754 | 0 | 0 | 0,09 | 0,1 |
| EMB262_3_10_MUC_32 | 0 | 0,07 | 754 | 0 | 0 | 0,05 | 0,1 |
| EMB262_13_8_MUC_10 | 0 | 0,13 | 864 | 0 | 0 | 0,08 | 0,03 |
| EMB262_13_8_MUC_10 | 0 | 0 | 864 | 0 | 0 | 0 | 0,03 |
| EMB262_13_8_MUC_10 | 0 | 0 | 864 | 0 | 0 | 0 | 0,03 |
| EMB262_3_10_MUC_11 | 124 | 25,5 | 297 | 208,75 | 177,05 | 42,93 | 37,43 |
| EMB262_3_10_MUC_11 | 101 | 20,7 | 297 | 170,03 | 177,05 | 34,85 | 37,43 |
| EMB262_3_10_MUC_11 | 90,5 | 20,5 | 297 | 152,36 | 177,05 | 34,51 | 37,43 |
| EMB262_3_10_MUC_25 | 0 | 0,4 | 540 | 0 | 0 | 0,37 | 0,28 |
| EMB262_3_10_MUC_25 | 0 | 0,06 | 540 | 0 | 0 | 0,06 | 0,28 |
| EMB262_3_10_MUC_25 | 0 | 0,44 | 540 | 0 | 0 | 0,41 | 0,28 |
| EMB262_12_2_MUC_6 | 387 | 1 | 864 | 223,96 | 130,02 | 0,58 | 0,34 |
| EMB262_12_2_MUC_6 | 156 | 0,42 | 864 | 90,28 | 130,02 | 0,24 | 0,34 |
| EMB262_12_2_MUC_6 | 131 | 0,33 | 864 | 75,81 | 130,02 | 0,19 | 0,34 |
| EMB262_3_10_MUC_28 | 0,06 | 0,64 | 769 | 0,04 | 0,01 | 0,42 | 0,24 |
| EMB262_3_10_MUC_28 | 0 | 0,13 | 769 | 0 | 0,01 | 0,08 | 0,24 |
| EMB262_3_10_MUC_28 | 0 | 0,32 | 769 | 0 | 0,01 | 0,21 | 0,24 |
| NTC | 0 | 0 | - | 0 | | 0 | |
| EMB262_13_8_MUC_16 | 0,12 | 15,5 | 707 | 0,08 | 0,06 | 10,96 | 8,11 |
| EMB262_13_8_MUC_16 | 0,07 | 10,1 | 707 | 0,05 | 0,06 | 7,14 | 8,11 |
| EMB262_13_8_MUC_16 | 0,06 | 8,8 | 707 | 0,04 | 0,06 | 6,22 | 8,11 |
| EMB262_13_8_MUC_24 | 0 | 1,4 | 762 | 0 | 0 | 0,92 | 0,54 |
| EMB262_13_8_MUC_24 | 0 | 0,37 | 762 | 0 | 0 | 0,24 | 0,54 |
| EMB262_13_8_MUC_24 | 0 | 0,68 | 762 | 0 | 0 | 0,45 | 0,54 |
| EMB262_12_2_MUC_33 | 0,12 | 0,81 | 757 | 0,08 | 0,03 | 0,54 | 0,33 |
| EMB262_12_2_MUC_33 | 0 | 0,7 | 757 | 0 | 0,03 | 0,46 | 0,33 |
| EMB262_12_2_MUC_33 | 0 | 0 | 757 | 0 | 0,03 | 0 | 0,33 |
| EMB262_6_28_MUC_9 | 0 | 0,73 | 629 | 0 | 0 | 0,58 | 0,29 |

| | | | | | | | |
|---|---|---|---|---|---|---|---|
| EMB262_6_28_MUC_9 | 0 | 0,31 | 629 | 0 | 0 | 0,25 | 0,29 |
| EMB262_6_28_MUC_9 | 0 | 0,06 | 629 | 0 | 0 | 0,05 | 0,29 |
| EMB262_12_2_MUC_27 | 0,14 | 1 | 426 | 0,16 | 0,11 | 1,17 | 0,7 |
| EMB262_12_2_MUC_27 | 0,07 | 0,34 | 426 | 0,08 | 0,11 | 0,4 | 0,7 |
| EMB262_12_2_MUC_27 | 0,06 | 0,45 | 426 | 0,07 | 0,11 | 0,53 | 0,7 |
| EMB262_13_8_MUC_12 | 0 | 48,2 | 840 | 0 | 0 | 28,69 | 18,39 |
| EMB262_13_8_MUC_12 | 0 | 25,6 | 840 | 0 | 0 | 15,24 | 18,39 |
| EMB262_13_8_MUC_12 | 0 | 18,9 | 840 | 0 | 0 | 11,25 | 18,39 |
| EMB262_12_2_MUC_25 | 0,13 | 1,1 | 594 | 0,11 | 0,04 | 0,93 | 0,36 |
| EMB262_12_2_MUC_25 | 0 | 0,2 | 594 | 0 | 0,04 | 0,17 | 0,36 |
| EMB262_12_2_MUC_25 | 0 | 0 | 594 | 0 | 0,04 | 0 | 0,36 |
| EMB262_3_10_MUC_22 | 14,2 | 1,3 | 794 | 8,94 | 6,99 | 0,82 | 0,64 |
| EMB262_3_10_MUC_22 | 10,5 | 1,1 | 794 | 6,61 | 6,99 | 0,69 | 0,64 |
| EMB262_3_10_MUC_22 | 8,6 | 0,66 | 794 | 5,42 | 6,99 | 0,42 | 0,64 |
| EMB262_3_10_MUC_33 | 0 | 0,78 | 822 | 0 | 0 | 0,47 | 0,3 |
| EMB262_3_10_MUC_33 | 0 | 0,49 | 822 | 0 | 0 | 0,3 | 0,3 |
| EMB262_3_10_MUC_33 | 0 | 0,2 | 822 | 0 | 0 | 0,12 | 0,3 |
| NTC | 0 | 0 | - | 0 | 0 | 0 | 0 |
| EMB262_12_2_MUC_30 | 0 | 0,45 | 533 | 0 | 0 | 0,42 | 0,34 |
| EMB262_12_2_MUC_30 | 0 | 0,44 | 533 | 0 | 0 | 0,41 | 0,34 |
| EMB262_12_2_MUC_30 | 0 | 0,21 | 533 | 0 | 0 | 0,2 | 0,34 |
| EMB262_6_28_MUC_19 | 0 | 0,62 | 660 | 0 | 0,02 | 0,47 | 0,34 |
| EMB262_6_28_MUC_19 | 0,06 | 0,57 | 660 | 0,05 | 0,02 | 0,43 | 0,34 |
| EMB262_6_28_MUC_19 | 0 | 0,14 | 660 | 0 | 0,02 | 0,11 | 0,34 |
| EMB262_12_2_MUC_14 | 4,3 | 0,39 | 669 | 3,21 | 2,42 | 0,29 | 0,39 |
| EMB262_12_2_MUC_14 | 2,8 | 0,75 | 669 | 2,09 | 2,42 | 0,56 | 0,39 |
| EMB262_12_2_MUC_14 | 2,6 | 0,43 | 669 | 1,94 | 2,42 | 0,32 | 0,39 |
| EMB262_3_10_MUC_7 | 26,6 | 0,18 | 472 | 28,18 | 22,88 | 0,19 | 0,17 |
| EMB262_3_10_MUC_7 | 20,2 | 0,06 | 472 | 21,4 | 22,88 | 0,06 | 0,17 |
| EMB262_3_10_MUC_7 | 18 | 0,25 | 472 | 19,07 | 22,88 | 0,26 | 0,17 |
| EMB262_6_28_MUC_38 | 0 | 0,08 | 633 | 0 | 0 | 0,06 | 0,07 |
| EMB262_6_28_MUC_38 | 0 | 0,18 | 633 | 0 | 0 | 0,14 | 0,07 |
| EMB262_6_28_MUC_38 | 0 | 0 | 633 | 0 | 0 | 0 | 0,07 |
| EMB262_6_28_MUC_31 | 0 | 0,17 | 396 | 0 | 0 | 0,21 | 0,26 |
| EMB262_6_28_MUC_31 | 0 | 0,23 | 396 | 0 | 0 | 0,29 | 0,26 |
| EMB262_6_28_MUC_31 | 0 | 0,21 | 396 | 0 | 0 | 0,27 | 0,26 |
| EMB262_6_28_MUC_39 | 0 | 0,14 | 695 | 0 | 0 | 0,1 | 0,07 |
| EMB262_6_28_MUC_39 | 0 | 0,07 | 695 | 0 | 0 | 0,05 | 0,07 |
| EMB262_6_28_MUC_39 | 0 | 0,07 | 695 | 0 | 0 | 0,05 | 0,07 |
| EMB262_13_8_MUC_15 | 0 | 0,13 | 766 | 0 | 0 | 0,08 | 0,04 |
| EMB262_13_8_MUC_15 | 0 | 0,07 | 766 | 0 | 0 | 0,05 | 0,04 |
| EMB262_13_8_MUC_15 | 0 | 0 | 766 | 0 | 0 | 0 | 0,04 |
| EMB262_6_28_MUC_28 | 0 | 2,6 | 719 | 0 | 0 | 1,81 | 1,12 |
| EMB262_6_28_MUC_28 | 0 | 0,73 | 719 | 0 | 0 | 0,51 | 1,12 |
| EMB262_6_28_MUC_28 | 0 | 1,5 | 719 | 0 | 0 | 1,04 | 1,12 |
| NTC | 0 | 0 | - | 0 | 0 | 0 | 0 |
| EMB262_6_28_MUC_35 | 0 | 0,4 | 677 | 0 | 0 | 0,3 | 0,31 |

| | | | | | | | |
|---|---|---|---|---|---|---|---|
| EMB262_6_28_MUC_35 | 0 | 0,41 | 677 | 0 | 0 | 0,3 | 0,31 |
| EMB262_6_28_MUC_35 | 0 | 0,46 | 677 | 0 | 0 | 0,34 | 0,31 |
| EMB262_6_28_MUC_27 | 0 | 0,28 | 679 | 0 | 0 | 0,21 | 0,22 |
| EMB262_6_28_MUC_27 | 0 | 0,18 | 679 | 0 | 0 | 0,13 | 0,22 |
| EMB262_6_28_MUC_27 | 0 | 0,42 | 679 | 0 | 0 | 0,31 | 0,22 |
| EMB262_12_2_MUC_32 | 0 | 0,47 | 538 | 0 | 0 | 0,44 | 0,25 |
| EMB262_12_2_MUC_32 | 0 | 0,17 | 538 | 0 | 0 | 0,16 | 0,25 |
| EMB262_12_2_MUC_32 | 0 | 0,16 | 538 | 0 | 0 | 0,15 | 0,25 |
| EMB262_3_10_MUC_24 | 0 | 0,17 | 717 | 0 | 0 | 0,12 | 0,12 |
| EMB262_3_10_MUC_24 | 0 | 0,23 | 717 | 0 | 0 | 0,16 | 0,12 |
| EMB262_3_10_MUC_24 | 0 | 0,11 | 717 | 0 | 0 | 0,08 | 0,12 |
| EMB262_12_2_MUC_35 | 0,08 | 0,46 | 447 | 0,09 | 0,07 | 0,51 | 0,35 |
| EMB262_12_2_MUC_35 | 0,06 | 0,24 | 447 | 0,07 | 0,07 | 0,27 | 0,35 |
| EMB262_12_2_MUC_35 | 0,06 | 0,23 | 447 | 0,07 | 0,07 | 0,26 | 0,35 |
| EMB262_6_28_MUC_30 | 0,69 | 1,1 | 751 | 0,46 | 0,37 | 0,73 | 0,76 |
| EMB262_6_28_MUC_30 | 0,65 | 1,4 | 751 | 0,43 | 0,37 | 0,93 | 0,76 |
| EMB262_6_28_MUC_30 | 0,33 | 0,92 | 751 | 0,22 | 0,37 | 0,61 | 0,76 |
| EMB262_6_28_MUC_44 | 0 | 0,29 | 730 | 0 | 0 | 0,2 | 0,11 |
| EMB262_6_28_MUC_44 | 0 | 0,13 | 730 | 0 | 0 | 0,09 | 0,11 |
| EMB262_6_28_MUC_44 | 0 | 0,06 | 730 | 0 | 0 | 0,04 | 0,11 |
| EMB262_6_28_MUC_26 | 0,06 | 0,55 | 717 | 0,04 | 0,01 | 0,38 | 0,52 |
| EMB262_6_28_MUC_26 | 0 | 0,72 | 717 | 0 | 0,01 | 0,5 | 0,52 |
| EMB262_6_28_MUC_26 | 0 | 0,95 | 717 | 0 | 0,01 | 0,66 | 0,52 |
| EMB262_6_28_MUC_36 | 0 | 0,63 | 526 | 0 | 0 | 0,6 | 0,52 |
| EMB262_6_28_MUC_36 | 0 | 0,58 | 526 | 0 | 0 | 0,55 | 0,52 |
| EMB262_6_28_MUC_36 | 0 | 0,42 | 526 | 0 | 0 | 0,4 | 0,52 |
| NTC | 0 | 0 | - | 0 | 0 | 0 | 0 |
| EMB262_13_8_MUC_4 | 1,6 | 1,9 | 534 | 1,5 | 1,53 | 1,78 | 1,31 |
| EMB262_13_8_MUC_4 | 1,2 | 1,5 | 534 | 1,12 | 1,53 | 1,4 | 1,31 |
| EMB262_13_8_MUC_4 | 2,1 | 0,8 | 534 | 1,97 | 1,53 | 0,75 | 1,31 |
| EMB262_6_28_MUC_32 | 0 | 0 | 539 | 0 | 0 | 0 | 0,09 |
| EMB262_6_28_MUC_32 | 0 | 0,2 | 539 | 0 | 0 | 0,19 | 0,09 |
| EMB262_6_28_MUC_32 | 0 | 0,1 | 539 | 0 | 0 | 0,09 | 0,09 |
| EMB262_6_28_MUC_15 | 0 | 2,8 | 479 | 0 | 0 | 2,92 | 1,57 |
| EMB262_6_28_MUC_15 | 0 | 1,7 | 479 | 0 | 0 | 1,77 | 1,57 |
| EMB262_6_28_MUC_15 | 0 | 0 | 479 | 0 | 0 | 0 | 1,57 |
| EMB262_3_10_MUC_21 | 5,9 | 1,5 | 508 | 5,81 | 3,94 | 1,48 | 0,77 |
| EMB262_3_10_MUC_21 | 3,4 | 0,39 | 508 | 3,35 | 3,94 | 0,38 | 0,77 |
| EMB262_3_10_MUC_21 | 2,7 | 0,45 | 508 | 2,66 | 3,94 | 0,44 | 0,77 |
| EMB262_6_28_MUC_45 | 0,13 | 0 | 531 | 0,12 | 0,04 | 0 | 0,03 |
| EMB262_6_28_MUC_45 | 0 | 0 | 531 | 0 | 0,04 | 0 | 0,03 |
| EMB262_6_28_MUC_45 | 0 | 0,11 | 531 | 0 | 0,04 | 0,1 | 0,03 |
| EMB262_13_8_MUC_3 | 7 | 2,7 | 623 | 5,62 | 3,29 | 2,17 | 1,23 |
| EMB262_13_8_MUC_3 | 3,6 | 1,2 | 623 | 2,89 | 3,29 | 0,96 | 1,23 |
| EMB262_13_8_MUC_3 | 1,7 | 0,7 | 623 | 1,36 | 3,29 | 0,56 | 1,23 |
| EMB262_6_28_MUC_24 | 0 | 1,2 | 631 | 0 | 0 | 0,95 | 0,64 |
| EMB262_6_28_MUC_24 | 0 | 0,8 | 631 | 0 | 0 | 0,63 | 0,64 |

| | | | | | | | |
|---|---|---|---|---|---|---|---|
| EMB262_6_28_MUC_24 | 0 | 0,42 | 631 | 0 | 0 | 0,33 | 0,64 |
| EMB262_12_2_MUC_29 | 0 | 0,5 | 650 | 0 | 0,03 | 0,38 | 0,42 |
| EMB262_12_2_MUC_29 | 0,1 | 0,7 | 650 | 0,08 | 0,03 | 0,54 | 0,42 |
| EMB262_12_2_MUC_29 | 0 | 0,43 | 650 | 0 | 0,03 | 0,33 | 0,42 |
| EMB262_6_28_MUC_48 | 0 | 0,34 | 521 | 0 | 0,04 | 0,33 | 0,28 |
| EMB262_6_28_MUC_48 | 0 | 0,31 | 521 | 0 | 0,04 | 0,3 | 0,28 |
| EMB262_6_28_MUC_48 | 0,12 | 0,24 | 521 | 0,12 | 0,04 | 0,23 | 0,28 |
| NTC | 0 | 0 | - | 0 | 0 | 0 | 0 |
| EMB262_6_28_MUC_42 | 0 | 0 | 619 | 0 | 0 | 0 | 0 |
| EMB262_6_28_MUC_42 | 0 | 0 | 619 | 0 | 0 | 0 | 0 |
| EMB262_6_28_MUC_42 | 0 | 0 | 619 | 0 | 0 | 0 | 0 |
| EMB262_6_28_MUC_29 | 0,06 | 1,2 | 563 | 0,05 | 0,04 | 1,07 | 0,76 |
| EMB262_6_28_MUC_29 | 0,06 | 0,68 | 563 | 0,05 | 0,04 | 0,6 | 0,76 |
| EMB262_6_28_MUC_29 | 0 | 0,68 | 563 | 0 | 0,04 | 0,6 | 0,76 |
| EMB262_13_8_MUC_5 | 0 | 148 | 406 | 0 | 0 | 182,27 | 120,81 |
| EMB262_13_8_MUC_5 | 0 | 81,5 | 406 | 0 | 0 | 100,37 | 120,81 |
| EMB262_13_8_MUC_5 | 0 | 64,8 | 406 | 0 | 0 | 79,8 | 120,81 |
| EMB262_3_10_MUC_34 | 0 | 0,19 | 739 | 0 | 0 | 0,13 | 0,18 |
| EMB262_3_10_MUC_34 | 0 | 0,49 | 739 | 0 | 0 | 0,33 | 0,18 |
| EMB262_3_10_MUC_34 | 0 | 0,12 | 739 | 0 | 0 | 0,08 | 0,18 |
| EMB262_6_28_MUC_41 | 0 | 0,88 | 629 | 0 | 0 | 0,7 | 0,42 |
| EMB262_6_28_MUC_41 | 0 | 0,36 | 629 | 0 | 0 | 0,29 | 0,42 |
| EMB262_6_28_MUC_41 | 0 | 0,36 | 629 | 0 | 0 | 0,29 | 0,42 |
| EMB262_13_8_MUC_13 | 0,56 | 1,8 | 807 | 0,35 | 0,31 | 1,12 | 0,97 |
| EMB262_13_8_MUC_13 | 0,37 | 1,3 | 807 | 0,23 | 0,31 | 0,81 | 0,97 |
| EMB262_13_8_MUC_13 | 0,55 | 1,6 | 807 | 0,34 | 0,31 | 0,99 | 0,97 |
| EMB262_6_28_MUC_49 | 0 | 1,1 | 578 | 0 | 0 | 0,95 | 0,67 |
| EMB262_6_28_MUC_49 | 0 | 0,66 | 578 | 0 | 0 | 0,57 | 0,67 |
| EMB262_6_28_MUC_49 | 0 | 0,57 | 578 | 0 | 0 | 0,49 | 0,67 |
| EMB262_13_8_MUC_8 | 0 | 0,17 | 848 | 0 | 0 | 0,1 | 0,03 |
| EMB262_13_8_MUC_8 | 0 | 0 | 848 | 0 | 0 | 0 | 0,03 |
| EMB262_13_8_MUC_8 | 0 | 0 | 848 | 0 | 0 | 0 | 0,03 |
| EMB262_6_28_MUC_47 | 0 | 1,2 | 629 | 0 | 0 | 0,95 | 0,77 |
| EMB262_6_28_MUC_47 | 0 | 0,96 | 629 | 0 | 0 | 0,76 | 0,77 |
| EMB262_6_28_MUC_47 | 0 | 0,76 | 629 | 0 | 0 | 0,6 | 0,77 |
| NTC | 0 | 0 | - | 0 | 0 | 0 | 0 |
| EMB262_3_10_MUC_23 | 0 | 0,35 | 593 | 0 | 0 | 0,3 | 0,28 |
| EMB262_3_10_MUC_23 | 0 | 0,31 | 593 | 0 | 0 | 0,26 | 0,28 |
| EMB262_3_10_MUC_23 | 0 | 0,34 | 593 | 0 | 0 | 0,29 | 0,28 |
| EMB262_6_28_MUC_43 | 0 | 0,31 | 918 | 0 | 0 | 0,17 | 0,11 |
| EMB262_6_28_MUC_43 | 0 | 0,15 | 918 | 0 | 0 | 0,08 | 0,11 |
| EMB262_6_28_MUC_43 | 0 | 0,14 | 918 | 0 | 0 | 0,08 | 0,11 |
| EMB262_12_2_MUC_18 | 0 | 2,3 | 522 | 0 | 0 | 2,2 | 1,33 |
| EMB262_12_2_MUC_18 | 0 | 1 | 522 | 0 | 0 | 0,96 | 1,33 |
| EMB262_12_2_MUC_18 | 0 | 0,85 | 522 | 0 | 0 | 0,81 | 1,33 |
| EMB262_12_2_MUC_20 | 0,06 | 1,5 | 363 | 0,08 | 0,08 | 2,07 | 1,8 |
| EMB262_12_2_MUC_20 | 0,12 | 0,91 | 363 | 0,17 | 0,08 | 1,25 | 1,8 |

| | | | | | | | |
|---|---|---|---|---|---|---|---|
| EMB262_12_2_MUC_20 | 0 | 1,5 | 363 | 0 | 0,08 | 2,07 | 1,8 |
| EMB262_13_8_MUC_6 | 1,5 | 10,9 | 774 | 0,97 | 1,12 | 7,04 | 8,42 |
| EMB262_13_8_MUC_6 | 1,2 | 16,3 | 774 | 0,78 | 1,12 | 10,53 | 8,42 |
| EMB262_13_8_MUC_6 | 2,5 | 11,9 | 774 | 1,61 | 1,12 | 7,69 | 8,42 |
| EMB262_13_8_MUC_1 | 0 | 12,5 | 633 | 0 | 0 | 9,87 | 7,19 |
| EMB262_13_8_MUC_1 | 0 | 8,6 | 633 | 0 | 0 | 6,79 | 7,19 |
| EMB262_13_8_MUC_1 | 0 | 6,2 | 633 | 0 | 0 | 4,9 | 7,19 |
| EMB262_13_8_MUC_20 | 0,48 | 4,1 | 564 | 0,43 | 0,4 | 3,63 | 3,49 |
| EMB262_13_8_MUC_20 | 0,35 | 3,6 | 564 | 0,31 | 0,4 | 3,19 | 3,49 |
| EMB262_13_8_MUC_20 | 0,51 | 4,1 | 564 | 0,45 | 0,4 | 3,63 | 3,49 |
| EMB262_13_8_MUC_2 | 22 | 2,6 | 613 | 17,94 | 15,36 | 2,12 | 2,37 |
| EMB262_13_8_MUC_2 | 16,4 | 3 | 613 | 13,38 | 15,36 | 2,45 | 2,37 |
| EMB262_13_8_MUC_2 | 18,1 | 3,1 | 613 | 14,76 | 15,36 | 2,53 | 2,37 |
| EMB262_6_28_MUC_40 | 0 | 1,1 | 653 | 0 | 0 | 0,84 | 0,56 |
| EMB262_6_28_MUC_40 | 0 | 0,71 | 653 | 0 | 0 | 0,54 | 0,56 |
| EMB262_6_28_MUC_40 | 0 | 0,4 | 653 | 0 | 0 | 0,31 | 0,56 |
| NTC | 0 | 0 | - | 0 | 0 | 0 | 0 |
| EMB262_13_8_MUC_7 | 0 | 0,06 | 581 | 0 | 0 | 0,05 | 0,11 |
| EMB262_13_8_MUC_7 | 0 | 0,12 | 581 | 0 | 0 | 0,1 | 0,11 |
| EMB262_13_8_MUC_7 | 0 | 0,19 | 581 | 0 | 0 | 0,16 | 0,11 |
| EMB262_13_8_MUC_11 | 2,7 | 4 | 819 | 1,65 | 1,06 | 2,44 | 1,63 |
| EMB262_13_8_MUC_11 | 1,8 | 2,4 | 819 | 1,1 | 1,06 | 1,47 | 1,63 |
| EMB262_13_8_MUC_11 | 0,71 | 1,6 | 819 | 0,43 | 1,06 | 0,98 | 1,63 |
| EMB262_6_28_MUC_33 | 0 | 2,2 | 477 | 0 | 0 | 2,31 | 1,82 |
| EMB262_6_28_MUC_33 | 0 | 1,7 | 477 | 0 | 0 | 1,78 | 1,82 |
| EMB262_6_28_MUC_33 | 0 | 1,3 | 477 | 0 | 0 | 1,36 | 1,82 |
| Extraction Blank | 0 | 0 | - | 0 | 0 | 0 | 0 |
| Extraction Blank | 0 | 0 | - | 0 | 0 | 0 | 0 |
| Extraction Blank | 0 | 0 | - | 0 | 0 | 0 | 0 |
| EMB262_12_2_GC_24 | 0 | 0,06 | 304 | 0 | 0,07 | 0,1 | 0,25 |
| EMB262_12_2_GC_24 | 0,06 | 0,17 | 304 | 0,1 | 0,07 | 0,28 | 0,25 |
| EMB262_12_2_GC_24 | 0,06 | 0,23 | 304 | 0,1 | 0,07 | 0,38 | 0,25 |
| EMB262_6_30_GC_22 | 0 | 6,3 | 689 | 0 | 0 | 4,57 | 3,99 |
| EMB262_6_30_GC_22 | 0 | 5 | 689 | 0 | 0 | 3,63 | 3,99 |
| EMB262_6_30_GC_22 | 0 | 5,2 | 689 | 0 | 0 | 3,77 | 3,99 |
| EMB262_12_2_GC_12 | 1,7 | 0,67 | 573 | 1,48 | 1,31 | 0,58 | 0,65 |
| EMB262_12_2_GC_12 | 1,5 | 0,86 | 573 | 1,31 | 1,31 | 0,75 | 0,65 |
| EMB262_12_2_GC_12 | 1,3 | 0,69 | 573 | 1,13 | 1,31 | 0,6 | 0,65 |
| EMB262_12_2_GC_11 | 1,9 | 0,78 | 866 | 1,1 | 1,04 | 0,45 | 0,35 |
| EMB262_12_2_GC_11 | 1,6 | 0,57 | 866 | 0,92 | 1,04 | 0,33 | 0,35 |
| EMB262_12_2_GC_11 | 1,9 | 0,48 | 866 | 1,1 | 1,04 | 0,28 | 0,35 |
| EMB262_12_2_MUC_10 | 22,5 | 0,61 | 581 | 19,36 | 20,08 | 0,52 | 0,66 |
| EMB262_12_2_MUC_10 | 26,2 | 1,06 | 581 | 22,55 | 20,08 | 0,91 | 0,66 |
| EMB262_12_2_MUC_10 | 21,3 | 0,64 | 581 | 18,33 | 20,08 | 0,55 | 0,66 |
| - | 0 | 0 | - | 0 | 0 | 0 | 0 |
| EMB262_6_28_MUC_46 | 0 | 0,89 | 808 | 0 | 0 | 0,55 | 0,49 |
| EMB262_6_28_MUC_46 | 0 | 0,69 | 808 | 0 | 0 | 0,43 | 0,49 |

| | | | | | | | |
|---|---|---|---|---|---|---|---|
| EMB262_6_28_MUC_46 | 0 | 0,78 | 808 | 0 | 0 | 0,48 | 0,49 |
| EMB262_6_30_GC_17 | 0 | 0,66 | 759 | 0 | 0,02 | 0,43 | 0,28 |
| EMB262_6_30_GC_17 | 0,08 | 0,31 | 759 | 0,05 | 0,02 | 0,2 | 0,28 |
| EMB262_6_30_GC_17 | 0 | 0,29 | 759 | 0 | 0,02 | 0,19 | 0,28 |
| EMB262_6_30_GC_42 | 0 | 2 | 721 | 0 | 0,01 | 1,39 | 1,16 |
| EMB262_6_30_GC_42 | 0,06 | 1,9 | 721 | 0,04 | 0,01 | 1,32 | 1,16 |
| EMB262_6_30_GC_42 | 0 | 1,1 | 721 | 0 | 0,01 | 0,76 | 1,16 |
| EMB262_6_30_GC_23 | 0 | 1,7 | 644 | 0 | 0 | 1,32 | 1,14 |
| EMB262_6_30_GC_23 | 0 | 1,1 | 644 | 0 | 0 | 0,85 | 1,14 |
| EMB262_6_30_GC_23 | 0 | 1,6 | 644 | 0 | 0 | 1,24 | 1,14 |
| EMB262_6_28_MUC_37 | 0 | 0,12 | 477 | 0 | 0 | 0,13 | 0,1 |
| EMB262_6_28_MUC_37 | 0 | 0,06 | 477 | 0 | 0 | 0,06 | 0,1 |
| EMB262_6_28_MUC_37 | 0 | 0,11 | 477 | 0 | 0 | 0,12 | 0,1 |
| EMB262_6_28_MUC_50 | 0 | 6,8 | 666 | 0 | 0 | 5,11 | 3,58 |
| EMB262_6_28_MUC_50 | 0 | 3,5 | 666 | 0 | 0 | 2,63 | 3,58 |
| EMB262_6_28_MUC_50 | 0 | 4 | 666 | 0 | 0 | 3 | 3,58 |
| EMB262_12_2_GC_28 | 0,06 | 1,6 | 323 | 0,09 | 0,17 | 2,48 | 2,17 |
| EMB262_12_2_GC_28 | 0,06 | 1,4 | 323 | 0,09 | 0,17 | 2,17 | 2,17 |
| EMB262_12_2_GC_28 | 0,2 | 1,2 | 323 | 0,31 | 0,17 | 1,86 | 2,17 |
| EMB262_6_30_GC_15 | 0 | 0 | 429 | 0 | 0 | 0 | 0,67 |
| EMB262_6_30_GC_15 | 0 | 1 | 429 | 0 | 0 | 1,17 | 0,67 |
| EMB262_6_30_GC_15 | 0 | 0,72 | 429 | 0 | 0 | 0,84 | 0,67 |
| EMB262_12_2_MUC_6 | 368 | 1,3 | 817 | 225,21 | 147,08 | 0,8 | 0,55 |
| EMB262_12_2_MUC_6 | 182 | 0,64 | 817 | 111,38 | 147,08 | 0,39 | 0,55 |
| EMB262_12_2_MUC_6 | 171 | 0,77 | 817 | 104,65 | 147,08 | 0,47 | 0,55 |
| NTC | 0 | 0 | - | 0 | 0 | 0 | 0 |
| EMB262_12_2_GC_29 | 8,5 | 2,1 | 589 | 7,22 | 4,33 | 1,78 | 1,25 |
| EMB262_12_2_GC_29 | 4,2 | 1,5 | 589 | 3,57 | 4,33 | 1,27 | 1,25 |
| EMB262_12_2_GC_29 | 2,6 | 0,82 | 589 | 2,21 | 4,33 | 0,7 | 1,25 |
| EMB262_12_2_GC_32 | 0,06 | 0,76 | 775 | 0,04 | 0,04 | 0,49 | 0,46 |
| EMB262_12_2_GC_32 | 0,06 | 0,52 | 775 | 0,04 | 0,04 | 0,34 | 0,46 |
| EMB262_12_2_GC_32 | 0,06 | 0,87 | 775 | 0,04 | 0,04 | 0,56 | 0,46 |
| EMB262_12_2_GC_13 | 0,06 | 0,9 | 304 | 0,1 | 0,14 | 1,48 | 1,02 |
| EMB262_12_2_GC_13 | 0,12 | 0,47 | 304 | 0,2 | 0,14 | 0,77 | 1,02 |
| EMB262_12_2_GC_13 | 0,07 | 0,49 | 304 | 0,12 | 0,14 | 0,81 | 1,02 |
| Extraction Blank | 0 | 0 | - | 0 | 0 | 0 | 0 |
| Extraction Blank | 0 | 0 | - | 0 | 0 | 0 | 0 |
| Extraction Blank | 0 | 0 | - | 0 | 0 | 0 | 0 |
| EMB262_6_30_GC_27 | 0 | 0,64 | 445 | 0 | 0 | 0,72 | 0,75 |
| EMB262_6_30_GC_27 | 0 | 0,65 | 445 | 0 | 0 | 0,73 | 0,75 |
| EMB262_6_30_GC_27 | 0 | 0,7 | 445 | 0 | 0 | 0,79 | 0,75 |
| EMB262_12_2_GC_33 | 0,12 | 2,5 | 680 | 0,09 | 0,03 | 1,84 | 1,16 |
| EMB262_12_2_GC_33 | 0 | 0,93 | 680 | 0 | 0,03 | 0,68 | 1,16 |
| EMB262_12_2_GC_33 | 0 | 1,3 | 680 | 0 | 0,03 | 0,96 | 1,16 |
| EMB262_12_2_GC_18 | 2,2 | 0,89 | 793 | 1,39 | 1,37 | 0,56 | 0,65 |
| EMB262_12_2_GC_18 | 2,5 | 0,82 | 793 | 1,58 | 1,37 | 0,52 | 0,65 |
| EMB262_12_2_GC_18 | 1,8 | 1,4 | 793 | 1,13 | 1,37 | 0,88 | 0,65 |

| | | | | | | | |
|---|---|---|---|---|---|---|---|
| EMB262_6_30_GC_19 | 0 | 1,05 | 552 | 0 | 0 | 0,95 | 1,05 |
| EMB262_6_30_GC_19 | 0 | 1,6 | 552 | 0 | 0 | 1,45 | 1,05 |
| EMB262_6_30_GC_19 | 0 | 0,84 | 552 | 0 | 0 | 0,76 | 1,05 |
| NTC | 0 | 0 | - | 0 | 0 | 0 | 0 |
| EMB262_6_30_GC_29 | 0 | 0,29 | 308 | 0 | 0 | 0,47 | 0,47 |
| EMB262_6_30_GC_29 | 0 | 0,3 | 308 | 0 | 0 | 0,49 | 0,47 |
| EMB262_6_30_GC_29 | 0 | 0,28 | 308 | 0 | 0 | 0,45 | 0,47 |
| EMB262_12_2_GC_16 | 0 | 0,38 | 265 | 0 | 0,08 | 0,72 | 1,13 |
| EMB262_12_2_GC_16 | 0,06 | 0,65 | 265 | 0,11 | 0,08 | 1,23 | 1,13 |
| EMB262_12_2_GC_16 | 0,06 | 0,76 | 265 | 0,11 | 0,08 | 1,43 | 1,13 |
| EMB262_6_28_MUC_28 | 0 | 2,2 | 650 | 0 | 0,02 | 1,69 | 1,33 |
| EMB262_6_28_MUC_28 | 0,06 | 1,7 | 650 | 0,05 | 0,02 | 1,31 | 1,33 |
| EMB262_6_28_MUC_28 | 0 | 1,3 | 650 | 0 | 0,02 | 1 | 1,33 |
| EMB262_6_30_GC_31 | 0 | 0,49 | 725 | 0 | 0 | 0,34 | 0,19 |
| EMB262_6_30_GC_31 | 0 | 0,11 | 725 | 0 | 0 | 0,08 | 0,19 |
| EMB262_6_30_GC_31 | 0 | 0,22 | 725 | 0 | 0 | 0,15 | 0,19 |
| EMB262_6_30_GC_31 | 0 | 0,55 | 725 | 0 | 0 | 0,38 | 0,25 |
| EMB262_6_30_GC_31 | 0 | 0 | 725 | 0 | 0 | 0 | 0,25 |
| EMB262_6_30_GC_31 | 0 | 0,52 | 725 | 0 | 0 | 0,36 | 0,25 |
| EMB262_6_30_GC_32 | 0 | 0,43 | 672 | 0 | 0 | 0,32 | 0,42 |
| EMB262_6_30_GC_32 | 0 | 0,66 | 672 | 0 | 0 | 0,49 | 0,42 |
| EMB262_6_30_GC_32 | 0 | 0,59 | 672 | 0 | 0 | 0,44 | 0,42 |
| EMB262_6_30_GC_33 | 0 | 0,76 | 779 | 0 | 0 | 0,49 | 0,38 |
| EMB262_6_30_GC_33 | 0 | 0,4 | 779 | 0 | 0 | 0,26 | 0,38 |
| EMB262_6_30_GC_33 | 0 | 0,6 | 779 | 0 | 0 | 0,39 | 0,38 |
| EMB262_6_30_GC_26 | 0,11 | 2,6 | 766 | 0,07 | 0,02 | 1,7 | 1,52 |
| EMB262_6_30_GC_26 | 0 | 2,1 | 766 | 0 | 0,02 | 1,37 | 1,52 |
| EMB262_6_30_GC_26 | 0 | 2,3 | 766 | 0 | 0,02 | 1,5 | 1,52 |
| EMB262_12_2_GC_14 | 0,17 | 1,1 | 409 | 0,21 | 0,09 | 1,34 | 1,16 |
| EMB262_12_2_GC_14 | 0 | 1,2 | 409 | 0 | 0,09 | 1,47 | 1,16 |
| EMB262_12_2_GC_14 | 0,06 | 0,55 | 409 | 0,07 | 0,09 | 0,67 | 1,16 |
| NTC | 0 | 0 | - | 0 | 0 | 0 | 0 |
| EMB262_12_2_GC_15 | 4,9 | 0,76 | 797 | 3,07 | 2,55 | 0,48 | 0,37 |
| EMB262_12_2_GC_15 | 3,7 | 0,75 | 797 | 2,32 | 2,55 | 0,47 | 0,37 |
| EMB262_12_2_GC_15 | 3,6 | 0,28 | 797 | 2,26 | 2,55 | 0,18 | 0,37 |
| EMB262_6_30_GC_16 | 0 | 0,39 | 611 | 0 | 0 | 0,32 | 0,33 |
| EMB262_6_30_GC_16 | 0 | 0,58 | 611 | 0 | 0 | 0,47 | 0,33 |
| EMB262_6_30_GC_16 | 0 | 0,24 | 611 | 0 | 0 | 0,2 | 0,33 |
| EMB262_6_30_GC_25 | 0 | 5,8 | 855 | 0 | 0 | 3,39 | 2,42 |
| EMB262_6_30_GC_25 | 0 | 3,6 | 855 | 0 | 0 | 2,11 | 2,42 |
| EMB262_6_30_GC_25 | 0 | 3 | 855 | 0 | 0 | 1,75 | 2,42 |
| EMB262_12_2_GC_35 | 0 | 0,88 | 714 | 0 | 0,01 | 0,62 | 0,51 |
| EMB262_12_2_GC_35 | 0 | 0,85 | 714 | 0 | 0,01 | 0,6 | 0,51 |
| EMB262_12_2_GC_35 | 0,06 | 0,44 | 714 | 0,04 | 0,01 | 0,31 | 0,51 |
| EMB262_6_30_GC_20 | 0 | 2,6 | 476 | 0 | 0 | 2,73 | 2,45 |
| EMB262_6_30_GC_20 | 0 | 1,9 | 476 | 0 | 0 | 2 | 2,45 |
| EMB262_6_30_GC_20 | 0 | 2,5 | 476 | 0 | 0 | 2,63 | 2,45 |

| | | | | | | | |
|---|---|---|---|---|---|---|---|
| EMB262_12_2_GC_31 | 0 | 0,41 | 819 | 0 | 0 | 0,25 | 0,43 |
| EMB262_12_2_GC_31 | 0 | 0,75 | 819 | 0 | 0 | 0,46 | 0,43 |
| EMB262_12_2_GC_31 | 0 | 0,94 | 819 | 0 | 0 | 0,57 | 0,43 |
| EMB262_12_2_MUC_4 | 5230 | 1,9 | 692 | 3778,9 | 2820,33 | 1,37 | 0,88 |
| EMB262_12_2_MUC_4 | 3330 | 0,78 | 692 | 2406,07 | 2820,33 | 0,56 | 0,88 |
| EMB262_12_2_MUC_4 | 3150 | 0,98 | 692 | 2276,01 | 2820,33 | 0,71 | 0,88 |
| Extraction Blank | 0 | 0 | - | 0 | 0 | 0 | 0 |
| Extraction Blank | 0 | 0 | - | 0 | 0 | 0 | 0 |
| Extraction Blank | 0 | 0 | - | 0 | 0 | 0 | 0 |
| EMB262_12_2_GC_30 | 0 | 0,97 | 661 | 0 | 0 | 0,73 | 0,88 |
| EMB262_12_2_GC_30 | 0 | 1,4 | 661 | 0 | 0 | 1,06 | 0,88 |
| EMB262_12_2_GC_30 | 0 | 1,13 | 661 | 0 | 0 | 0,85 | 0,88 |
| NTC | 0 | 0 | - | 0 | 0 | 0 | 0 |
| EMB262_6_30_GC_39 | 0 | 1,2 | 422 | 0 | 0 | 1,42 | 0,87 |
| EMB262_6_30_GC_39 | 0 | 0,59 | 422 | 0 | 0 | 0,7 | 0,87 |
| EMB262_6_30_GC_39 | 0 | 0,42 | 422 | 0 | 0 | 0,5 | 0,87 |
| EMB262_12_2_GC_21 | 0 | 0,96 | 478 | 0 | 0 | 1 | 0,66 |
| EMB262_12_2_GC_21 | 0 | 0,46 | 478 | 0 | 0 | 0,48 | 0,66 |
| EMB262_12_2_GC_21 | 0 | 0,46 | 478 | 0 | 0 | 0,48 | 0,66 |
| EMB262_12_2_GC_40 | 0 | 1,7 | 859 | 0 | 0 | 0,99 | 1,18 |
| EMB262_12_2_GC_40 | 0 | 2 | 859 | 0 | 0 | 1,16 | 1,18 |
| EMB262_12_2_GC_40 | 0 | 2,4 | 859 | 0 | 0 | 1,4 | 1,18 |
| EMB262_6_30_GC_29 | 0,06 | 8,7 | 756 | 0,04 | 0,01 | 5,75 | 4,7 |
| EMB262_6_30_GC_29 | 0 | 6,7 | 756 | 0 | 0,01 | 4,43 | 4,7 |
| EMB262_6_30_GC_29 | 0 | 5,9 | 756 | 0 | 0,01 | 3,9 | 4,7 |
| EMB262_12_2_GC_50 | 0,06 | 0,76 | 503 | 0,06 | 0,02 | 0,76 | 0,67 |
| EMB262_12_2_GC_50 | 0 | 0,61 | 503 | 0 | 0,02 | 0,61 | 0,67 |
| EMB262_12_2_GC_50 | 0 | 0,64 | 503 | 0 | 0,02 | 0,64 | 0,67 |
| EMB262_12_2_GC_23 | 0,06 | 1,3 | 473 | 0,06 | 0,02 | 1,37 | 0,95 |
| EMB262_12_2_GC_23 | 0 | 0,67 | 473 | 0 | 0,02 | 0,71 | 0,95 |
| EMB262_12_2_GC_23 | 0 | 0,73 | 473 | 0 | 0,02 | 0,77 | 0,95 |
| EMB262_12_2_GC_36 | 0 | 0,24 | 830 | 0 | 0 | 0,14 | 0,16 |
| EMB262_12_2_GC_36 | 0 | 0,45 | 830 | 0 | 0 | 0,27 | 0,16 |
| EMB262_12_2_GC_36 | 0 | 0,12 | 830 | 0 | 0 | 0,07 | 0,16 |
| EMB262_12_2_GC_30 | 0,06 | 4 | 761 | 0,04 | 0,03 | 2,63 | 2,04 |
| EMB262_12_2_GC_30 | 0,06 | 2,6 | 761 | 0,04 | 0,03 | 1,71 | 2,04 |
| EMB262_12_2_GC_30 | 0 | 2,7 | 761 | 0 | 0,03 | 1,77 | 2,04 |
| EMB262_12_2_GC_19 | 0 | 0,31 | 629 | 0 | 0 | 0,25 | 0,16 |
| EMB262_12_2_GC_19 | 0 | 0,19 | 629 | 0 | 0 | 0,15 | 0,16 |
| EMB262_12_2_GC_19 | 0 | 0,12 | 629 | 0 | 0 | 0,1 | 0,16 |
| NTC | 0 | 0 | - | 0 | 0 | 0 | 0 |
| EMB262_12_2_GC_38 | 0 | 0,5 | 719 | 0 | 0 | 0,35 | 0,16 |
| EMB262_12_2_GC_38 | 0 | 0,2 | 719 | 0 | 0 | 0,14 | 0,16 |
| EMB262_12_2_GC_38 | 0 | 0 | 719 | 0 | 0 | 0 | 0,16 |
| EMB262_6_30_GC_38 | 0 | 0,75 | 662 | 0 | 0 | 0,57 | 0,73 |
| EMB262_6_30_GC_38 | 0 | 1,2 | 662 | 0 | 0 | 0,91 | 0,73 |
| EMB262_6_30_GC_38 | 0 | 0,93 | 662 | 0 | 0 | 0,7 | 0,73 |

| | | | | | | | |
|---|---|---|---|---|---|---|---|
| EMB262_6_30_GC_14 | 0 | 1,6 | 625 | 0 | 0 | 1,28 | 0,99 |
| EMB262_6_30_GC_14 | 0 | 1,1 | 625 | 0 | 0 | 0,88 | 0,99 |
| EMB262_6_30_GC_14 | 0 | 1 | 625 | 0 | 0 | 0,8 | 0,99 |
| EMB262_6_30_GC_43 | 0 | 0,22 | 644 | 0 | 0 | 0,17 | 0,25 |
| EMB262_6_30_GC_43 | 0 | 0,54 | 644 | 0 | 0 | 0,42 | 0,25 |
| EMB262_6_30_GC_43 | 0 | 0,21 | 644 | 0 | 0 | 0,16 | 0,25 |
| EMB262_6_30_GC_18 | 0 | 1,4 | 579 | 0 | 0 | 1,21 | 0,89 |
| EMB262_6_30_GC_18 | 0 | 0,86 | 579 | 0 | 0 | 0,74 | 0,89 |
| EMB262_6_30_GC_18 | 0 | 0,82 | 579 | 0 | 0 | 0,71 | 0,89 |
| Extraction Blank | 0 | 0 | - | 0 | 0 | 0 | 0 |
| Extraction Blank | 0 | 0 | - | 0 | 0 | 0 | 0 |
| Extraction Blank | 0 | 0 | - | 0 | 0 | 0 | 0 |
| EMB262_12_2_GC_42 | 0 | 0,4 | 606 | 0 | 0 | 0,33 | 0,3 |
| EMB262_12_2_GC_42 | 0 | 0,31 | 606 | 0 | 0 | 0,26 | 0,3 |
| EMB262_12_2_GC_42 | 0 | 0,37 | 606 | 0 | 0 | 0,31 | 0,3 |
| EMB262_12_2_GC_17 | 1,8 | 0,49 | 721 | 1,25 | 0,93 | 0,34 | 0,35 |
| EMB262_12_2_GC_17 | 0,84 | 0,66 | 721 | 0,58 | 0,93 | 0,46 | 0,35 |
| EMB262_12_2_GC_17 | 1,4 | 0,38 | 721 | 0,97 | 0,93 | 0,26 | 0,35 |
| EMB262_12_2_GC_26 | 0,08 | 0,23 | 220 | 0,18 | 0,06 | 0,52 | 0,34 |
| EMB262_12_2_GC_26 | 0 | 0 | 220 | 0 | 0,06 | 0 | 0,34 |
| EMB262_12_2_GC_26 | 0 | 0,22 | 220 | 0 | 0,06 | 0,5 | 0,34 |
| NTC | 0 | 0 | - | 0 | 0 | 0 | 0 |
| EMB262_6_30_GC_21 | 0,08 | 5,4 | 430 | 0,09 | 0,03 | 6,28 | 4,73 |
| EMB262_6_30_GC_21 | 0 | 3,2 | 430 | 0 | 0,03 | 3,72 | 4,73 |
| EMB262_6_30_GC_21 | 0 | 3,6 | 430 | 0 | 0,03 | 4,19 | 4,73 |
| EMB262_12_2_GC_22 | 0 | 0,23 | 432 | 0 | 0 | 0,27 | 0,27 |
| EMB262_12_2_GC_22 | 0 | 0,13 | 432 | 0 | 0 | 0,15 | 0,27 |
| EMB262_12_2_GC_22 | 0 | 0,33 | 432 | 0 | 0 | 0,38 | 0,27 |
| EMB262_6_30_GC_41 | 0 | 0,18 | 798 | 0 | 0 | 0,11 | 0,17 |
| EMB262_6_30_GC_41 | 0 | 0,25 | 798 | 0 | 0 | 0,16 | 0,17 |
| EMB262_6_30_GC_41 | 0 | 0,37 | 798 | 0 | 0 | 0,23 | 0,17 |
| EMB262_12_2_GC_25 | 0,13 | 1,1 | 596 | 0,11 | 0,05 | 0,92 | 0,87 |
| EMB262_12_2_GC_25 | 0,06 | 1 | 596 | 0,05 | 0,05 | 0,84 | 0,87 |
| EMB262_12_2_GC_25 | 0 | 1 | 596 | 0 | 0,05 | 0,84 | 0,87 |
| EMB262_6_30_GC_13 | 0,08 | 2,7 | 725 | 0,06 | 0,02 | 1,86 | 1,26 |
| EMB262_6_30_GC_13 | 0 | 1,3 | 725 | 0 | 0,02 | 0,9 | 1,26 |
| EMB262_6_30_GC_13 | 0 | 1,5 | 725 | 0 | 0,02 | 1,03 | 1,26 |
| EMB262_12_2_GC_20 | 0 | 0,13 | 857 | 0 | 0,01 | 0,08 | 0,13 |
| EMB262_12_2_GC_20 | 0,07 | 0,36 | 857 | 0,04 | 0,01 | 0,21 | 0,13 |
| EMB262_12_2_GC_20 | 0 | 0,2 | 857 | 0 | 0,01 | 0,12 | 0,13 |
| EMB262_12_2_GC_51 | 0,15 | 0,53 | 340 | 0,22 | 0,07 | 0,78 | 0,58 |
| EMB262_12_2_GC_51 | 0 | 0,28 | 340 | 0 | 0,07 | 0,41 | 0,58 |
| EMB262_12_2_GC_51 | 0 | 0,38 | 340 | 0 | 0,07 | 0,56 | 0,58 |
| EMB262_12_2_GC_39 | 0 | 1,1 | 873 | 0 | 0 | 0,63 | 0,46 |
| EMB262_12_2_GC_39 | 0 | 0,73 | 873 | 0 | 0 | 0,42 | 0,46 |
| EMB262_12_2_GC_39 | 0 | 0,59 | 873 | 0 | 0 | 0,34 | 0,46 |
| EMB262_12_2_GC_37 | 1,3 | 0,25 | 753 | 0,86 | 0,63 | 0,17 | 0,11 |

| | | | | | | | |
|---|---|---|---|---|---|---|---|
| EMB262_12_2_GC_37 | 0,9 | 0,12 | 753 | 0,6 | 0,63 | 0,08 | 0,11 |
| EMB262_12_2_GC_37 | 0,63 | 0,13 | 753 | 0,42 | 0,63 | 0,09 | 0,11 |
| NTC | 0 | 0 | - | 0 | 0 | 0 | 0 |
| EMB262_6_30_GC_24 | 0 | 8,9 | 882 | 0 | 0 | 5,05 | 3,42 |
| EMB262_6_30_GC_24 | 0 | 4,7 | 882 | 0 | 0 | 2,66 | 3,42 |
| EMB262_6_30_GC_24 | 0 | 4,5 | 882 | 0 | 0 | 2,55 | 3,42 |
| EMB262_6_30_GC_35 | 0 | 0,61 | 500 | 0 | 0 | 0,61 | 0,4 |
| EMB262_6_30_GC_35 | 0 | 0,44 | 500 | 0 | 0 | 0,44 | 0,4 |
| EMB262_6_30_GC_35 | 0 | 0,15 | 500 | 0 | 0 | 0,15 | 0,4 |
| EMB262_6_30_GC_40 | 0 | 0,81 | 612 | 0 | 0 | 0,66 | 0,33 |
| EMB262_6_30_GC_40 | 0 | 0,26 | 612 | 0 | 0 | 0,21 | 0,33 |
| EMB262_6_30_GC_40 | 0 | 0,14 | 612 | 0 | 0 | 0,11 | 0,33 |
| Extraction Blank | 0 | 0 | - | 0 | 0 | 0 | 0 |
| Extraction Blank | 0 | 0 | - | 0 | 0 | 0 | 0 |
| Extraction Blank | 0 | 0 | - | 0 | 0 | 0 | 0 |
| EMB262_12_2_GC_45 | 0 | 0,32 | 926 | 0 | 0,02 | 0,17 | 0,19 |
| EMB262_12_2_GC_45 | 0 | 0,48 | 926 | 0 | 0,02 | 0,26 | 0,19 |
| EMB262_12_2_GC_45 | 0,13 | 0,26 | 926 | 0,07 | 0,02 | 0,14 | 0,19 |
| EMB262_6_30_GC_12 | 0,06 | 0,67 | 918 | 0,03 | 0,01 | 0,36 | 0,42 |
| EMB262_6_30_GC_12 | 0 | 0,64 | 918 | 0 | 0,01 | 0,35 | 0,42 |
| EMB262_6_30_GC_12 | 0 | 1 | 918 | 0 | 0,01 | 0,54 | 0,42 |
| EMB262_12_2_GC_46 | 0 | 0,82 | 480 | 0 | 0 | 0,85 | 0,93 |
| EMB262_12_2_GC_46 | 0 | 1,1 | 480 | 0 | 0 | 1,15 | 0,93 |
| EMB262_12_2_GC_46 | 0 | 0,76 | 480 | 0 | 0 | 0,79 | 0,93 |
| EMB262_6_30_GC_51 | 0 | 11,1 | 818 | 0 | 0 | 6,78 | 5,85 |
| EMB262_6_30_GC_51 | 0 | 8,5 | 818 | 0 | 0 | 5,2 | 5,85 |
| EMB262_6_30_GC_51 | 0 | 9,1 | 818 | 0 | 0 | 5,56 | 5,85 |
| EMB262_6_30_GC_64 | 0 | 0,42 | 591 | 0 | 0 | 0,36 | 0,23 |
| EMB262_6_30_GC_64 | 0 | 0,28 | 591 | 0 | 0 | 0,24 | 0,23 |
| EMB262_6_30_GC_64 | 0 | 0,13 | 591 | 0 | 0 | 0,11 | 0,23 |
| NTC | 0 | 0 | - | 0 | 0 | 0 | 0 |
| EMB262_6_30_GC_62 | 0,07 | 2,1 | 643 | 0,05 | 0,02 | 1,63 | 1,22 |
| EMB262_6_30_GC_62 | 0 | 1,4 | 643 | 0 | 0,02 | 1,09 | 1,22 |
| EMB262_6_30_GC_62 | 0 | 1,2 | 643 | 0 | 0,02 | 0,93 | 1,22 |
| EMB262_12_2_GC_4 | 1 | 0,39 | 571 | 0,88 | 1,08 | 0,34 | 0,33 |
| EMB262_12_2_GC_4 | 1,3 | 0,44 | 571 | 1,14 | 1,08 | 0,39 | 0,33 |
| EMB262_12_2_GC_4 | 1,4 | 0,29 | 571 | 1,23 | 1,08 | 0,25 | 0,33 |
| EMB262_6_30_GC_7 | 0 | 0 | 939 | 0 | 0 | 0 | 0 |
| EMB262_6_30_GC_7 | 0 | 0 | 939 | 0 | 0 | 0 | 0 |
| EMB262_6_30_GC_7 | 0 | 0 | 939 | 0 | 0 | 0 | 0 |
| EMB262_6_30_GC_48 | 0 | 2,9 | 458 | 0 | 0,02 | 3,17 | 3,64 |
| EMB262_6_30_GC_48 | 0,06 | 3,8 | 458 | 0,07 | 0,02 | 4,15 | 3,64 |
| EMB262_6_30_GC_48 | 0 | 3,3 | 458 | 0 | 0,02 | 3,6 | 3,64 |
| EMB262_6_28_MUC_51 | 0 | 0,45 | 537 | 0 | 0 | 0,42 | 0,34 |
| EMB262_6_28_MUC_51 | 0 | 0,31 | 537 | 0 | 0 | 0,29 | 0,34 |
| EMB262_6_28_MUC_51 | 0 | 0,35 | 537 | 0 | 0 | 0,33 | 0,34 |
| EMB262_6_30_GC_47 | 0 | 8,6 | 802 | 0 | 0 | 5,36 | 3,93 |

| | | | | | | | |
|---|---|---|---|---|---|---|---|
| EMB262_6_30_GC_47 | 0 | 4,5 | 802 | 0 | 0 | 2,81 | 3,93 |
| EMB262_6_30_GC_47 | 0 | 5,8 | 802 | 0 | 0 | 3,62 | 3,93 |
| EMB262_6_30_GC_65 | 0,06 | 0 | 730 | 0,04 | 0,01 | 0 | 0 |
| EMB262_6_30_GC_65 | 0 | 0 | 730 | 0 | 0,01 | 0 | 0 |
| EMB262_6_30_GC_65 | 0 | 0 | 730 | 0 | 0,01 | 0 | 0 |
| EMB262_6_30_GC_2 | 0,15 | 0 | 281 | 0,27 | 0,13 | 0 | 0 |
| EMB262_6_30_GC_2 | 0,07 | 0 | 281 | 0,12 | 0,13 | 0 | 0 |
| EMB262_6_30_GC_2 | 0 | 0 | 281 | 0 | 0,13 | 0 | 0 |
| EMB262_6_30_GC_59 | 0 | 0 | 760 | 0 | 0 | 0 | 0 |
| EMB262_6_30_GC_59 | 0 | 0 | 760 | 0 | 0 | 0 | 0 |
| EMB262_6_30_GC_59 | 0 | 0 | 760 | 0 | 0 | 0 | 0 |
| NTC | 0 | 0 | - | 0 | 0 | 0 | 0 |
| EMB262_12_2_MUC_2 | 374 | 4,3 | 605 | 309,09 | 257,85 | 3,55 | 3,55 |
| EMB262_12_2_MUC_2 | 327 | 4,6 | 605 | 270,25 | 257,85 | 3,8 | 3,55 |
| EMB262_12_2_MUC_2 | 235 | 4 | 605 | 194,21 | 257,85 | 3,31 | 3,55 |
| Extraction Blank | 0 | 0 | - | 0 | 0 | 0 | 0 |
| Extraction Blank | 0 | 0 | - | 0 | 0 | 0 | 0 |
| Extraction Blank | 0 | 0 | - | 0 | 0 | 0 | 0 |
| EMB262_6_30_GC_57 | 0 | 0 | 722 | 0 | 0 | 0 | 0 |
| EMB262_6_30_GC_57 | 0 | 0 | 722 | 0 | 0 | 0 | 0 |
| EMB262_6_30_GC_57 | 0 | 0 | 722 | 0 | 0 | 0 | 0 |
| EMB262_12_2_GC_49 | 0,12 | 0,93 | 517 | 0,12 | 0,08 | 0,9 | 0,44 |
| EMB262_12_2_GC_49 | 0 | 0,06 | 517 | 0 | 0,08 | 0,06 | 0,44 |
| EMB262_12_2_GC_49 | 0,12 | 0,36 | 517 | 0,12 | 0,08 | 0,35 | 0,44 |
| EMB262_6_30_GC_56 | 0 | 0 | 478 | 0 | 0,02 | 0 | 0 |
| EMB262_6_30_GC_56 | 0 | 0 | 478 | 0 | 0,02 | 0 | 0 |
| EMB262_6_30_GC_56 | 0,07 | 0 | 478 | 0,07 | 0,02 | 0 | 0 |
| EMB262_12_2_GC_5 | 6,2 | 0,47 | 495 | 6,26 | 5,93 | 0,47 | 0,42 |
| EMB262_12_2_GC_5 | 5,5 | 0,37 | 495 | 5,56 | 5,93 | 0,37 | 0,42 |
| EMB262_12_2_GC_5 | 5,9 | 0,41 | 495 | 5,96 | 5,93 | 0,41 | 0,42 |
| EMB262_6_30_GC_63 | 0 | 0,15 | 965 | 0 | 0 | 0,08 | 0,08 |
| EMB262_6_30_GC_63 | 0 | 0,21 | 965 | 0 | 0 | 0,11 | 0,08 |
| EMB262_6_30_GC_63 | 0 | 0,13 | 965 | 0 | 0 | 0,07 | 0,08 |
| EMB262_13_8_MUC_27 | 121 | 0,08 | 648 | 93,36 | 71,45 | 0,06 | 0,06 |
| EMB262_13_8_MUC_27 | 84,1 | 0,07 | 648 | 64,89 | 71,45 | 0,05 | 0,06 |
| EMB262_13_8_MUC_27 | 72,7 | 0,08 | 648 | 56,1 | 71,45 | 0,06 | 0,06 |
| EMB262_6_30_GC_44 | 0 | 3 | 632 | 0 | 0 | 2,37 | 2,14 |
| EMB262_6_30_GC_44 | 0 | 2,9 | 632 | 0 | 0 | 2,29 | 2,14 |
| EMB262_6_30_GC_44 | 0 | 2,2 | 632 | 0 | 0 | 1,74 | 2,14 |
| NTC | 0 | 0 | - | 0 | 0 | 0 | 0 |
| EMB262_6_30_GC_61 | 0 | 0 | 412 | 0 | 0 | 0 | 0 |
| EMB262_6_30_GC_61 | 0 | 0 | 412 | 0 | 0 | 0 | 0 |
| EMB262_6_30_GC_61 | 0 | 0 | 412 | 0 | 0 | 0 | 0 |
| EMB262_6_30_GC_46 | 0 | 3,3 | 514 | 0 | 0 | 3,21 | 2,56 |
| EMB262_6_30_GC_46 | 0 | 2,6 | 514 | 0 | 0 | 2,53 | 2,56 |
| EMB262_6_30_GC_46 | 0 | 2 | 514 | 0 | 0 | 1,95 | 2,56 |
| EMB262_6_30_GC_53 | 0 | 3,1 | 657 | 0 | 0 | 2,36 | 1,45 |

| | | | | | | | |
|---|---|---|---|---|---|---|---|
| EMB262_6_30_GC_53 | 0 | 1,3 | 657 | 0 | 0 | 0,99 | 1,45 |
| EMB262_6_30_GC_53 | 0 | 1,3 | 657 | 0 | 0 | 0,99 | 1,45 |
| EMB262_12_2_GC_9 | 0,06 | 0,25 | 870 | 0,03 | 0,01 | 0,14 | 0,3 |
| EMB262_12_2_GC_9 | 0 | 0,54 | 870 | 0 | 0,01 | 0,31 | 0,3 |
| EMB262_12_2_GC_9 | 0 | 0,79 | 870 | 0 | 0,01 | 0,45 | 0,3 |
| EMB262_6_30_GC_54 | 0 | 11,3 | 459 | 0 | 0 | 12,31 | 7,73 |
| EMB262_6_30_GC_54 | 0 | 10 | 459 | 0 | 0 | 10,89 | 7,73 |
| EMB262_6_30_GC_54 | 0 | 0 | 459 | 0 | 0 | 0 | 7,73 |
| EMB262_6_30_GC_50 | 0 | 16,6 | 490 | 0 | 0 | 16,94 | 12,21 |
| EMB262_6_30_GC_50 | 0 | 11,6 | 490 | 0 | 0 | 11,84 | 12,21 |
| EMB262_6_30_GC_50 | 0 | 7,7 | 490 | 0 | 0 | 7,86 | 12,21 |
| EMB262_12_2_GC_2 | 31,7 | 3,3 | 584 | 27,14 | 25,34 | 2,83 | 2,43 |
| EMB262_12_2_GC_2 | 29,1 | 2,6 | 584 | 24,91 | 25,34 | 2,23 | 2,43 |
| EMB262_12_2_GC_2 | 28 | 2,6 | 584 | 23,97 | 25,34 | 2,23 | 2,43 |
| EMB262_12_2_GC_10 | 0,18 | 1,4 | 836 | 0,11 | 0,09 | 0,84 | 0,53 |
| EMB262_12_2_GC_10 | 0,12 | 0,76 | 836 | 0,07 | 0,09 | 0,45 | 0,53 |
| EMB262_12_2_GC_10 | 0,13 | 0,52 | 836 | 0,08 | 0,09 | 0,31 | 0,53 |
| Extraction Blank | 0 | 0 | - | 0 | 0 | 0 | 0 |
| Extraction Blank | 0 | 0 | - | 0 | 0 | 0 | 0 |
| Extraction Blank | 0 | 0 | - | 0 | 0 | 0 | 0 |
| NTC | 0 | 0 | | 0 | 0 | 0 | 0 |
| EMB262_6_30_GC_1 | 0,1 | 0 | 945 | 0,05 | 0,02 | 0 | 0 |
| EMB262_6_30_GC_1 | 0 | 0 | 945 | 0 | 0,02 | 0 | 0 |
| EMB262_6_30_GC_1 | 0 | 0 | 945 | 0 | 0,02 | 0 | 0 |
| EMB262_12_2_GC_3 | 0 | 1,2 | 443 | 0 | 0 | 1,35 | 1 |
| EMB262_12_2_GC_3 | 0 | 0,8 | 443 | 0 | 0 | 0,9 | 1 |
| EMB262_12_2_GC_3 | 0 | 0,67 | 443 | 0 | 0 | 0,76 | 1 |
| EMB262_6_30_GC_49 | 0 | 7,3 | 582 | 0 | 0,02 | 6,27 | 5,99 |
| EMB262_6_30_GC_49 | 0 | 7,8 | 582 | 0 | 0,02 | 6,7 | 5,99 |
| EMB262_6_30_GC_49 | 0,08 | 5,8 | 582 | 0,07 | 0,02 | 4,98 | 5,99 |
| EMB262_6_30_GC_5 | 0 | 0 | 598 | 0 | 0 | 0 | 0 |
| EMB262_6_30_GC_5 | 0 | 0 | 598 | 0 | 0 | 0 | 0 |
| EMB262_6_30_GC_5 | 0 | 0 | 598 | 0 | 0 | 0 | 0 |
| EMB262_12_2_GC_47 | 0,07 | 0,59 | 717 | 0,05 | 0,02 | 0,41 | 0,47 |
| EMB262_12_2_GC_47 | 0 | 0,86 | 717 | 0 | 0,02 | 0,6 | 0,47 |
| EMB262_12_2_GC_47 | 0 | 0,59 | 717 | 0 | 0,02 | 0,41 | 0,47 |
| EMB262_12_2_GC_8 | 0,21 | 0,85 | 507 | 0,21 | 0,11 | 0,84 | 0,6 |
| EMB262_12_2_GC_8 | 0,13 | 0,6 | 507 | 0,13 | 0,11 | 0,59 | 0,6 |
| EMB262_12_2_GC_8 | 0 | 0,36 | 507 | 0 | 0,11 | 0,36 | 0,6 |
| EMB262_12_2_GC_1 | 0,07 | 0,33 | 388 | 0,09 | 0,09 | 0,43 | 0,57 |
| EMB262_12_2_GC_1 | 0,07 | 0,56 | 388 | 0,09 | 0,09 | 0,72 | 0,57 |
| EMB262_12_2_GC_1 | 0,07 | 0,43 | 388 | 0,09 | 0,09 | 0,55 | 0,57 |
| EMB262_6_30_GC_3 | 0,07 | 0 | 494 | 0,07 | 0,02 | 0 | 0 |
| EMB262_6_30_GC_3 | 0 | 0 | 494 | 0 | 0,02 | 0 | 0 |
| EMB262_6_30_GC_3 | 0 | 0 | 494 | 0 | 0,02 | 0 | 0 |
| EMB262_6_28_MUC_52 | 0 | 2,1 | 474 | 0 | 0 | 2,22 | 2,04 |
| EMB262_6_28_MUC_52 | 0 | 2,5 | 474 | 0 | 0 | 2,64 | 2,04 |

| | | | | | | | |
|---|---|---|---|---|---|---|---|
| EMB262_6_28_MUC_52 | 0 | 1,2 | 474 | 0 | 0 | 1,27 | 2,04 |
| NTC | 0 | 0 | | 0 | 0 | 0 | 0 |
| EMB262_6_30_GC_8 | 0 | 0 | 200 | 0 | 0 | 0 | 0 |
| EMB262_6_30_GC_8 | 0 | 0 | 200 | 0 | 0 | 0 | 0 |
| EMB262_6_30_GC_8 | 0 | 0 | 200 | 0 | 0 | 0 | 0 |
| EMB262_12_2_GC_44 | 0 | 0,87 | 484 | 0 | 0,08 | 0,9 | 0,84 |
| EMB262_12_2_GC_44 | 0,16 | 1,4 | 484 | 0,17 | 0,08 | 1,45 | 0,84 |
| EMB262_12_2_GC_44 | 0,08 | 0,16 | 484 | 0,08 | 0,08 | 0,17 | 0,84 |
| EMB262_6_30_GC_4 | 0 | 0 | 479 | 0 | 0 | 0 | 0 |
| EMB262_6_30_GC_4 | 0 | 0 | 479 | 0 | 0 | 0 | 0 |
| EMB262_6_30_GC_4 | 0 | 0 | 479 | 0 | 0 | 0 | 0 |
| EMB262_6_30_GC_10 | 0 | 0 | 744 | 0 | 0 | 0 | 0 |
| EMB262_6_30_GC_10 | 0 | 0 | 744 | 0 | 0 | 0 | 0 |
| EMB262_6_30_GC_10 | 0 | 0 | 744 | 0 | 0 | 0 | 0 |
| EMB262_6_30_GC_60 | 0 | 0 | 190 | 0 | 0 | 0 | 0 |
| EMB262_6_30_GC_60 | 0 | 0 | 190 | 0 | 0 | 0 | 0 |
| EMB262_6_30_GC_60 | 0 | 0 | 190 | 0 | 0 | 0 | 0 |
| EMB262_6_30_GC_9 | 0 | 0 | 529 | 0 | 0 | 0 | 0 |
| EMB262_6_30_GC_9 | 0 | 0 | 529 | 0 | 0 | 0 | 0 |
| EMB262_6_30_GC_9 | 0 | 0 | 529 | 0 | 0 | 0 | 0 |
| Extraction Blank | 0 | 0 | - | 0 | 0 | 0 | 0 |
| Extraction Blank | 0 | 0 | - | 0 | 0 | 0 | 0 |
| Extraction Blank | 0 | 0 | - | 0 | 0 | 0 | 0 |
| EMB262_6_30_GC_52 | 0 | 9,2 | 547 | 0 | 0 | 8,41 | 6,12 |
| EMB262_6_30_GC_52 | 0 | 5,6 | 547 | 0 | 0 | 5,12 | 6,12 |
| EMB262_6_30_GC_52 | 0 | 5,3 | 547 | 0 | 0 | 4,84 | 6,12 |
| EMB262_6_30_GC_6 | 0 | 0 | 641 | 0 | 0 | 0 | 0 |
| EMB262_6_30_GC_6 | 0 | 0 | 641 | 0 | 0 | 0 | 0 |
| EMB262_6_30_GC_6 | 0 | 0 | 641 | 0 | 0 | 0 | 0 |
| NTC | 0 | 0 | | 0 | 0 | 0 | 0 |
| EMB262_6_30_GC_36 | 0 | 10,9 | 531 | 0 | 0 | 10,26 | 6,56 |
| EMB262_6_30_GC_36 | 0 | 5,9 | 531 | 0 | 0 | 5,56 | 6,56 |
| EMB262_6_30_GC_36 | 0 | 4,1 | 531 | 0 | 0 | 3,86 | 6,56 |
| EMB262_6_30_GC_43 | 0 | 0,45 | 518 | 0 | 0 | 0,43 | 0,37 |
| EMB262_6_30_GC_43 | 0 | 0,48 | 518 | 0 | 0 | 0,46 | 0,37 |
| EMB262_6_30_GC_43 | 0 | 0,21 | 518 | 0 | 0 | 0,2 | 0,37 |
| EMB262_12_2_GC_7 | 0,07 | 1,7 | 484 | 0,07 | 0,07 | 1,76 | 1,45 |
| EMB262_12_2_GC_7 | 0 | 1,3 | 484 | 0 | 0,07 | 1,34 | 1,45 |
| EMB262_12_2_GC_7 | 0,13 | 1,2 | 484 | 0,13 | 0,07 | 1,24 | 1,45 |
| EMB262_12_2_MUC_8 | 415 | 2,6 | 508 | 408,46 | 300,52 | 2,56 | 2,2 |
| EMB262_12_2_MUC_8 | 269 | 2,4 | 508 | 264,76 | 300,52 | 2,36 | 2,2 |
| EMB262_12_2_MUC_8 | 232 | 1,7 | 508 | 228,35 | 300,52 | 1,67 | 2,2 |
| EMB262_6_30_GC_66 | 0 | 1,4 | 258 | 0 | 0,05 | 2,71 | 2,65 |
| EMB262_6_30_GC_66 | 0 | 1 | 258 | 0 | 0,05 | 1,94 | 2,65 |
| EMB262_6_30_GC_66 | 0,07 | 1,7 | 258 | 0,14 | 0,05 | 3,29 | 2,65 |
| EMB262_6_30_GC_45 | 0 | 1,5 | 541 | 0 | 0 | 1,39 | 0,91 |
| EMB262_6_30_GC_45 | 0 | 1 | 541 | 0 | 0 | 0,92 | 0,91 |

| | | | | | | | |
|---|---|---|---|---|---|---|---|
| EMB262_6_30_GC_45 | 0 | 0,46 | 541 | 0 | 0 | 0,43 | 0,91 |
| EMB262_12_2_GC_48 | 1,3 | 1,8 | 498 | 1,31 | 0,84 | 1,81 | 1,04 |
| EMB262_12_2_GC_48 | 1,2 | 1,3 | 498 | 1,2 | 0,84 | 1,31 | 1,04 |
| EMB262_12_2_GC_48 | 0 | 0 | 498 | 0 | 0,84 | 0 | 1,04 |
| EMB262_6_30_GC_55 | 0 | 12,4 | 674 | 0 | 0 | 9,2 | 6,58 |
| EMB262_6_30_GC_55 | 0 | 6,7 | 674 | 0 | 0 | 4,97 | 6,58 |
| EMB262_6_30_GC_55 | 0 | 7,5 | 674 | 0 | 0 | 5,56 | 6,58 |
| EMB262_12_2_GC_6 | 0 | 0 | 398 | 0 | 0,26 | 0 | 0,29 |
| EMB262_12_2_GC_6 | 0 | 0 | 398 | 0 | 0,26 | 0 | 0,29 |
| EMB262_12_2_GC_6 | 0,61 | 0,69 | 398 | 0,77 | 0,26 | 0,87 | 0,29 |
| NTC | 0,16 | 0,9 | - | 0 | 0 | 0 | 0 |
| EMB262_6_30_GC_37 | 0,22 | 0,51 | 576 | 0,19 | 0,06 | 0,44 | 0,15 |
| EMB262_6_30_GC_37 | 0 | 0 | 576 | 0 | 0,06 | 0 | 0,15 |
| EMB262_6_30_GC_37 | 0 | 0 | 576 | 0 | 0,06 | 0 | 0,15 |
| EMB262_6_30_GC_58 | 0 | 0 | 460 | 0 | 8,55 | 0 | 2,93 |
| EMB262_6_30_GC_58 | 14,8 | 4,8 | 460 | 16,09 | 8,55 | 5,22 | 2,93 |
| EMB262_6_30_GC_58 | 8,8 | 3,3 | 460 | 9,57 | 8,55 | 3,59 | 2,93 |
| EMB262_12_2_GC_43 | 9,7 | 3,8 | 585 | 8,29 | 2,76 | 3,25 | 1,23 |
| EMB262_12_2_GC_43 | 0 | 0 | 585 | 0 | 2,76 | 0 | 1,23 |
| EMB262_12_2_GC_43 | 0 | 0,53 | 585 | 0 | 2,76 | 0,45 | 1,23 |
| Extraction Blank | 0 | 0 | - | 0 | 0 | 0 | 0 |
| Extraction Blank | 0 | 0 | - | 0 | 0 | 0 | 0 |
| NTC | 0 | 0 | - | 0 | 0 | 0 | 0 |

### Analyses

#### ddPCR

The consistency between the replicates of FAM and Hex concentration was analyzed by performing a Pearson correlation. All replicates are significantly correlated with each other. They all show R-values higher than 0.99.

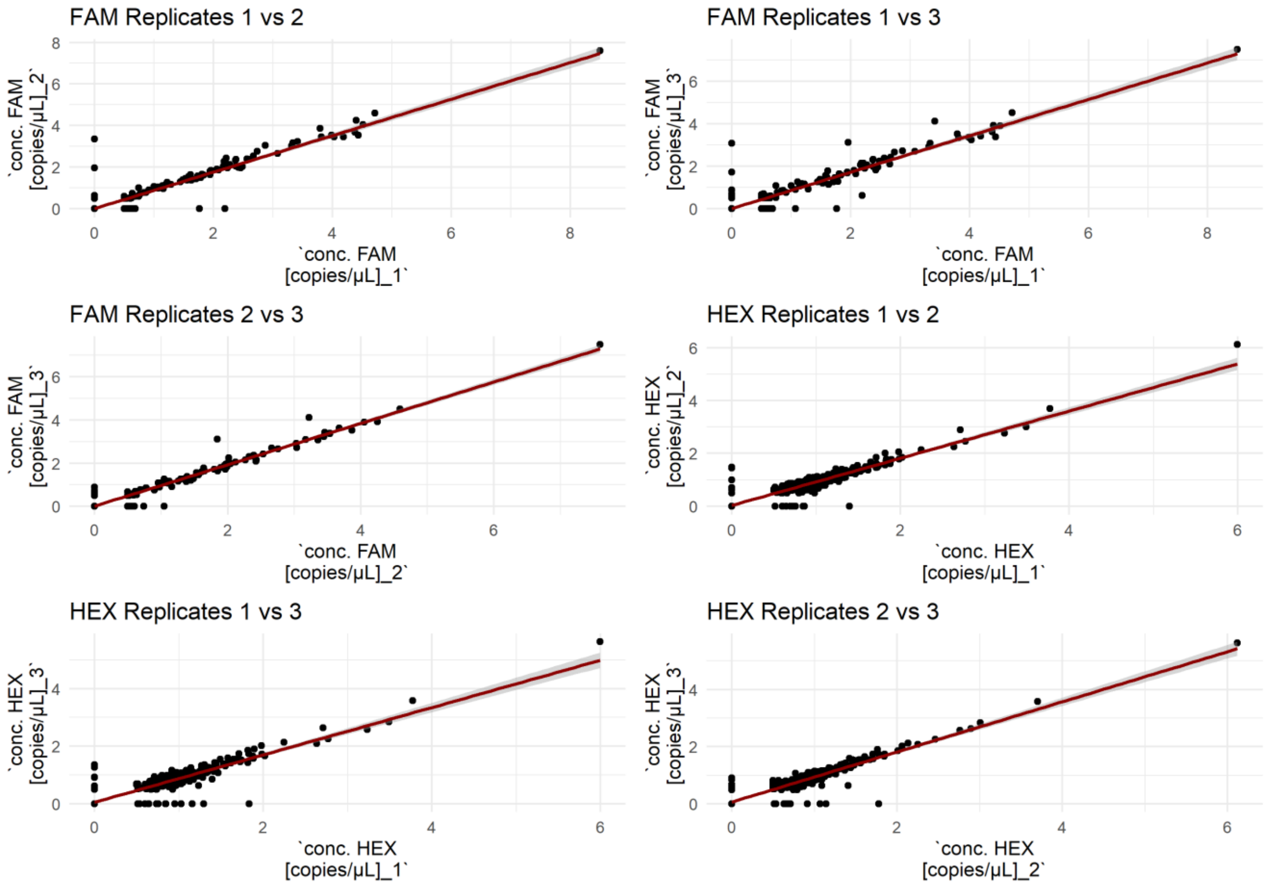

**Figure S4: Consistency between replicates of FAM and HEX concentrations. Each subplot represents a pair of replicates, with the double square root transformed concentrations plotted against each other. A linear regression line (red) is included in each subplot to visualize the trend. Pearson correlation and p-values for: FAM replicates: between Replicate 1 and Replicate 2 is R=0.9971 with a p=1.68e-302, between Replicate 1 and Replicate 3 is R=0.9955 with a p=1.82e-277, between Replicate 2 and Replicate 3 is R=0.9974 with a p=1.14e-308; HEX replicates: between Replicate 1 and Replicate 2 is R=0.9969 with p=8.75e-300, between Replicate 1 and Replicate 3 is R=0.9975 with p=6.30e-312, between Replicate 2 and Replicate 3 is R=0.9994 with a p=0.000. The high degree of correlation, as indicated by the correlation coefficients and p-values, suggests a high level of consistency between the replicates.**

#### Metabarcoding

The ordination of the community composition in all replicates of the metabarcoding dataset at each location resulted in similar replicate patterns, with the exception of a few individual replicates (as indicated in Figure S5).

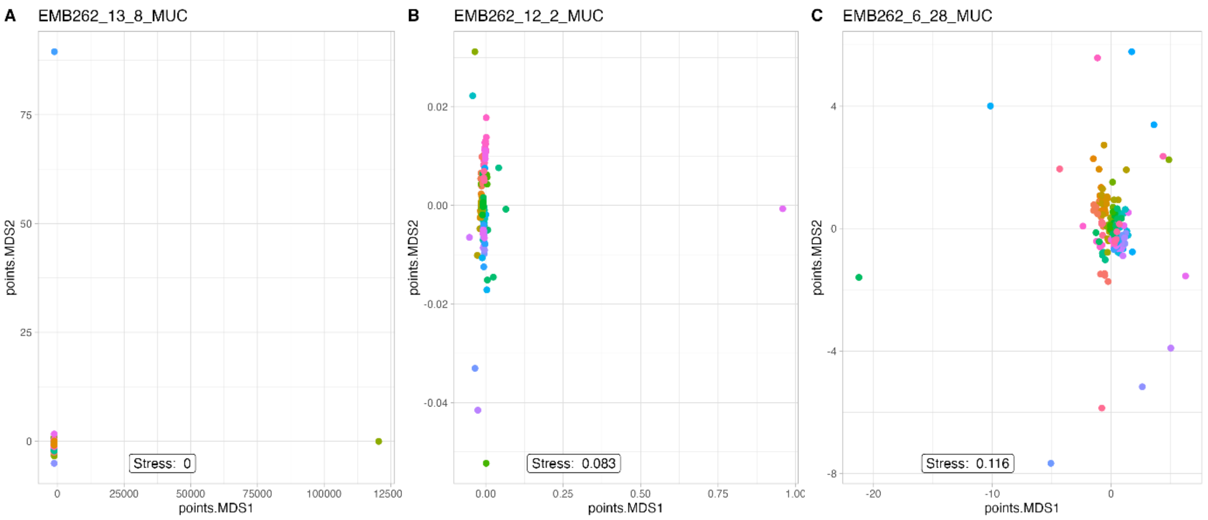

**Figure S5: Nonmetric multidimensional scaling analysis of Dinoflagellate and Diatom communities. The analysis is based on the Bray-Curtis distance and covers the time series from all short cores. Different colors represent different times.**

The relative abundance of the four datasets over time is shown in Figure S6. It can be observed that biomonitoring data has only been available since the 1980s, and that the relative abundance of the datasets varies dynamically. To investigate these dynamics further and to see if they are coherent, the datasets were compared and analyzed further. The occurrence of *S. marinoi* and *A. malmogiense* and its respective phylum are presented in the four different datasets: 1) ddPCR concentration; 2) Biomonitoring phylum; 3) Biomonitoring species; 4) Metabarcoding.

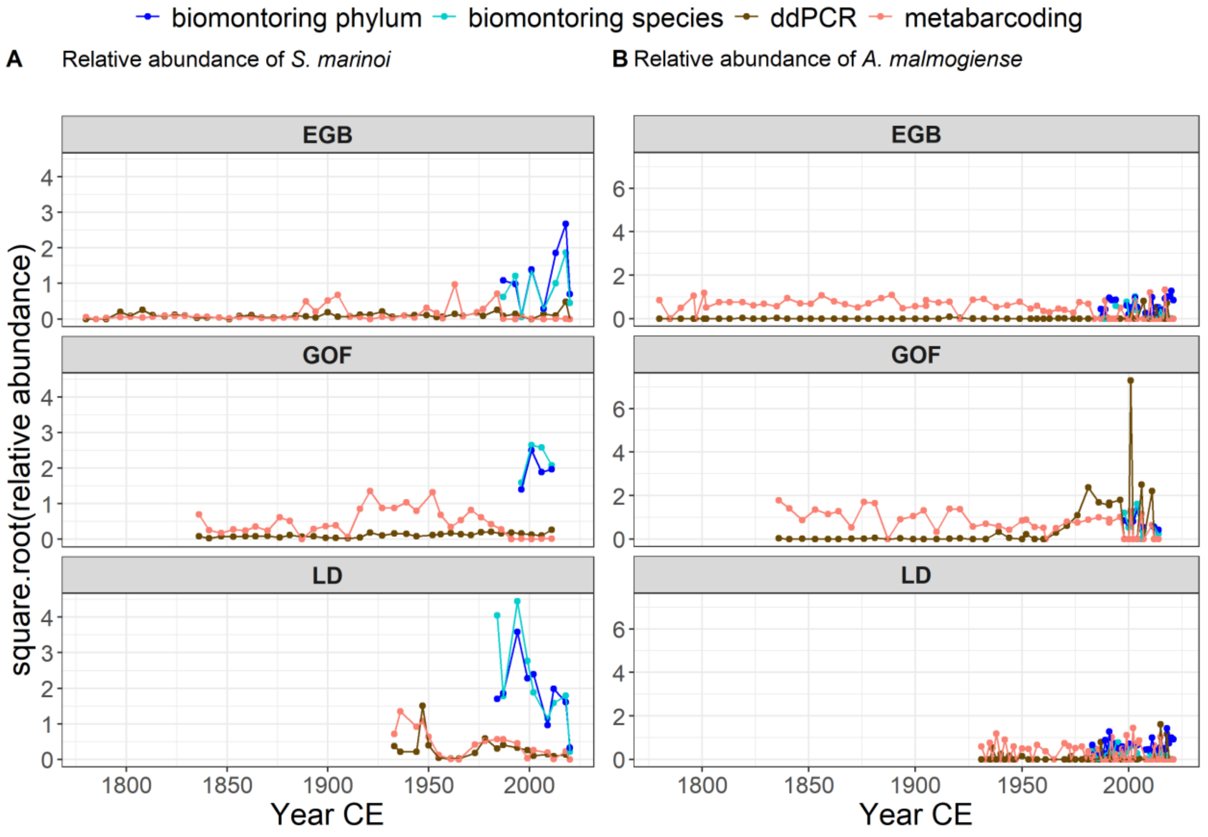

**Figure S6: Relative abundances of *S. marinoi* (A) and *A. malmogiense* (B) over time. Sampling sites are: Eastern Gotland Basin (EGB), Golf of Finland (GOF) and Landort Deep (LD).**

The Pearson's correlation analysis of the four datasets (ddPCR concentration, Biomonitoring phylum, Biomonitoring species, Metabarcoding) revealed significant relationships between the datasets for the two target groups, *A. malmogiense*/Dinoflagellates and *S. marinoi*/Diatoms.

For *A. malmogiense*/Dinoflagellates, the strongest correlation was observed between ddPCR concentration and Biomonitoring species (R = 0.89, p = 3.22E-07). A moderate correlation was also found between Metabarcoding and Biomonitoring phylum (R = 0.44, p = 0.059), and Biomonitoring species and Biomonitoring phylum (R = 0.47, p = 0.045).

For *S. marinoi*/Diatoms, the strongest correlation was found between Biomonitoring species and Biomonitoring phylum (R = 0.71, p = 0.0006). Moderate correlations were observed between ddPCR concentration and Biomonitoring phylum (R = 0.50, p = 0.029), Metabarcoding and Biomonitoring species (R = 0.62, p = 0.005), and ddPCR concentration and Metabarcoding (R = 0.49, p = 0.032).

These results suggest that the four datasets are interconnected and provide complementary information for the study of *A. malmogiense*/Dinoflagellates and *S. marinoi*/Diatoms.

**Table S7: Peason's correlation of the four datasets: 1) ddPCR concentration; 2) Biomonitoring phylum; 3) Biomonitoring species; 4) Metabarcoding. Datasets and their correlation results (R- and p-value) are shown for the two target groups (*A. malmogiense*/Dinoflagellates and *S. marinoi*/Diatoms)**

| Datasets | value | *A. malmogiense/* Dinoflagellates | *S. marinoi/* Diatoms |
|---|---|---|---|
| ddPCR & biomonitoring genus | R | 0,89060782 | 0,362923918 |
| | p | 3,21628E-07 | 0,126718133 |
| ddPCR & biomonitoring phylum | R | 0,413786183 | 0,501296307 |
| | p | 0,078216604 | 0,028775386 |
| metabarcoding & biomonitoring genus | R | 0,366870697 | 0,617927734 |
| | p | 0,122336776 | 0,004809492 |
| metabarcoding & biomonitoring phylum | R | 0,441315828 | 0,287460153 |
| | p | 0,058559247 | 0,23273728 |
| ddPCR & metabarcoding | R | 0,262939579 | 0,492375696 |
| | p | 0,276772233 | 0,032226012 |
| biomonitoring genus & biomonitoring phylum | R | 0,465203655 | 0,714625508 |
| | p | 0,044744266 | 0,00058597 |

A weak negative correlation (R = -0.0041) between the ddPCR concentration of *A. malmogiense* and *S. marinoi* is depicted in Figure S5. However, the p-value (p = 0.96) suggests that this correlation is not statistically significant. Therefore, there is no relationship between the concentrations.

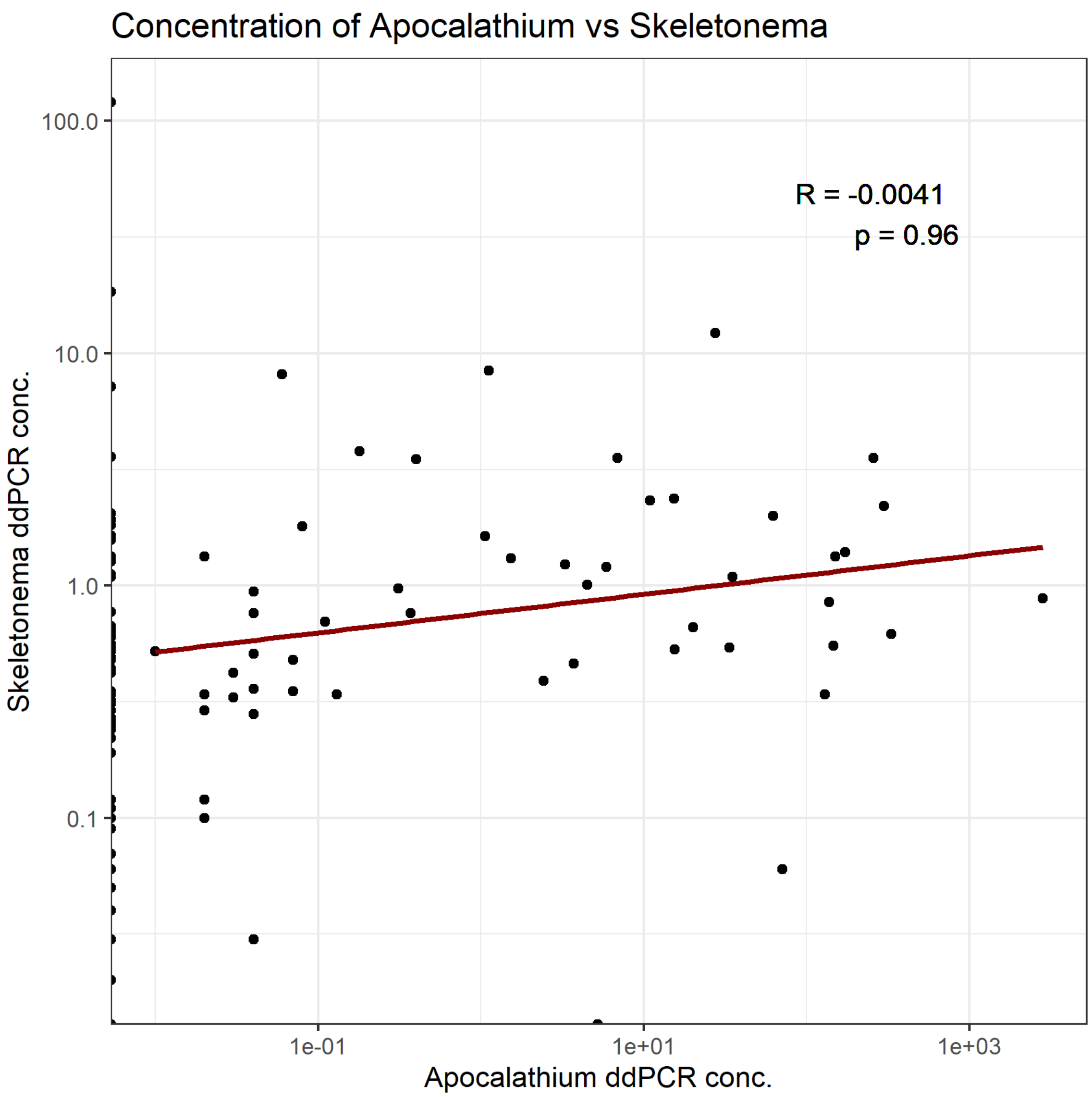

**Figure S7: Pearson's correlation between the ddPCR concentration of *S. marinoi* and *A. malmogiense* (R=-0.0041, p=0.96). Linear regression on the logarithm of the concentration of *S. marinoi* and *A. malmogiense*.**

#### Rscript availability

GitHub https://github.com/Alex-132/EI_paper_rep
